# Identification and partial reconstitution of the biosynthetic pathway of bioactive meroterpenoids from *Hericium erinaceus* (Lion’s Mane mushroom)

**DOI:** 10.1101/2024.07.16.603773

**Authors:** Riccardo Iacovelli, Fons Poon, Kristina Haslinger

## Abstract

*Hericium erinaceus* (Lion’s Mane mushroom) is widely consumed for its numerous reported benefits for brain health. A growing body of evidence suggests that these benefits are likely attributable to aromatics contained in its fruiting bodies, including the meroterpenoids hericenones. Here, we report the identification and reconstitution of the first two steps of the biosynthetic pathway of hericenones via heterologous expression of the PKS HerA and the carboxylic acid reductase HerB in *Aspergillus oryzae*. Furthermore, we investigate a putative prenyltransferase that might be responsible for the following biosynthetic step. Ongoing efforts to reconstitute the full pathway will enable large scale production of hericenones and other meroterpenoids in heterologous hosts.

---

Fungal natural products (NPs)—also known as secondary metabolites—are an important source for the discovery of bioactive compounds with pharmaceutical applications [1,2]. Traditionally, research in the field has focused on NPs from ascomycetes, particularly molds from the genera *Aspergillus* and *Penicillium*, given that these are genetically tractable and generally easy to cultivate in the laboratory [3–7]. Recent advances in -omics sciences allowed researchers to expand the scope of NP discovery to other groups of fungi, including edible mushrooms and medicinal mushrooms belonging to the phylum *Basidiomycota* [8–10]. Among these, the Lion’s Mane fungus—*Hericium erinaceus*—has gained considerable attention due to its reported health-promoting effects, attributable to the antioxidative, anti-inflammatory, and immunostimulating properties of its bioactive constituents [11,12]. These mainly include complex polysaccharides [13,14], the diterpenes erinacines, and orsellinic acid (ORA)-derived meroterpenoids (hericenones, hericerins, and erinacerins) (Fig. 1), mainly produced in the fruiting bodies [12,15–18]. The meroterpenoids belong to a broader class of compounds that exhibit potent biological activities with important applications in medicine [19,20]. For example, mycophenolic acid produced by *Penicillium brevicompactum* has long been a widely prescribed immunosuppressant medication [21,22]; cannabidiol extracted from *Cannabis* plants was recently approved for the treatment of seizures [23,24]; and daurichromenic acid isolated from the plant *Rhododendron dauricum* is being extensively studied for his remarkable anti-HIV activity [25,26]. Recent research has shown that the meroterpenoids from *H. erinaceus* exhibit neuroprotective and neuro-regenerative effects on isolated neuronal cells and in mouse models [15,17,27–30], which make these molecules interesting candidates to develop potential treatments for neurodegenerative diseases, such as dementia and Alzheimer’s, and for neuronal injuries. Although hericenones were first isolated from *H. erinaceus* more than 30 years ago [31], the biosynthetic machinery responsible for the synthesis of meroterpenoids in *H. erinaceus* remains yet to be identified. Knowing the responsible enzymes would provide access to enhanced compound production—for example by heterologous overexpression of the biosynthetic genes in model fungi or by metabolic engineering of the native producer. Thus, we set out to investigate the genome of *H. erinaceus* to identify the biosynthetic gene cluster (BGC) responsible for the biosynthesis of ORA-derived meroterpenoids and characterize it by means of heterologous expression.

**Figure 1.**
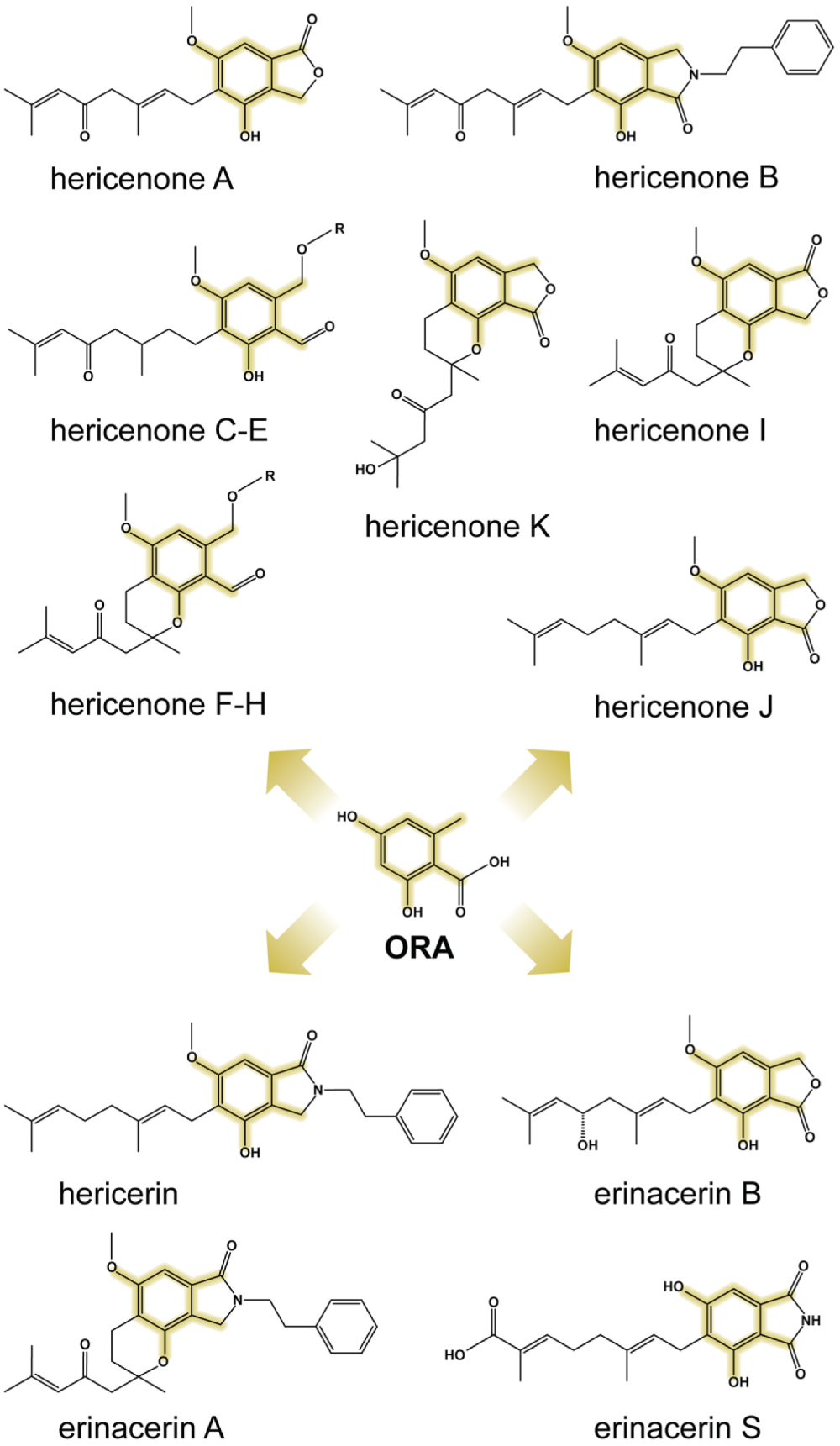
Revised structures [18] of some ORA-derived meroterpenoids from *H. erinaceus*. The orsellinic acid core (depicted in the center) is highlighted in yellow. For hericenones, R = palmitoyl (C, F), stearoyl (D, G), or linoleoyl (E, H).

Orsellinic acid—the core structure shared by hericenones, hericerin, and erinacerins—is one of the simplest aromatic polyketides, and it is biosynthesized by type III PKS in plants and by type I PKS in bacteria and fungi [32]. In several meroterpenoids of *H. erinaceus*, the carboxylic acid moiety is reduced to an aldehyde, indicating that an aromatic carboxylic acid reductase (CAR) might also be involved in the biosynthesis [33]. Meroterpenoid-producing BGCs that encode these enzymes have been recently identified in the ascomycetes *Stachybotrys bisbyi* [34] and *Acremonium egyptiacum* [35]. In both cases, a type I PKS (ORA synthase), a prenyltransferase (PT), and a CAR are involved in the first 3 biosynthetic steps that yield ilicicolin B (also called LL-Z 1272β), a farnesylated orsellinic aldehyde. Thus, we set out to identify a similar BGC in *H. erinaceus*. We retrieved the genome assembly of *H. erinaceus* 0605 from the NCBI database [36] and directly submitted it to the webtool fungiSMASH v7.0 [37] for BGC prediction. Out of the 12 predicted BGCs (additional file 2), we identified only 1 BGC encoding a type I non-reducing PKS and two putative NRPS-like enzymes, the enzyme family that CARs belong to (Fig. 2). Interestingly, this PKS had previously been reported to be overexpressed in the meroterpenoid-producing fruiting bodies compared to the mycelium [38].

**Figure 2.**
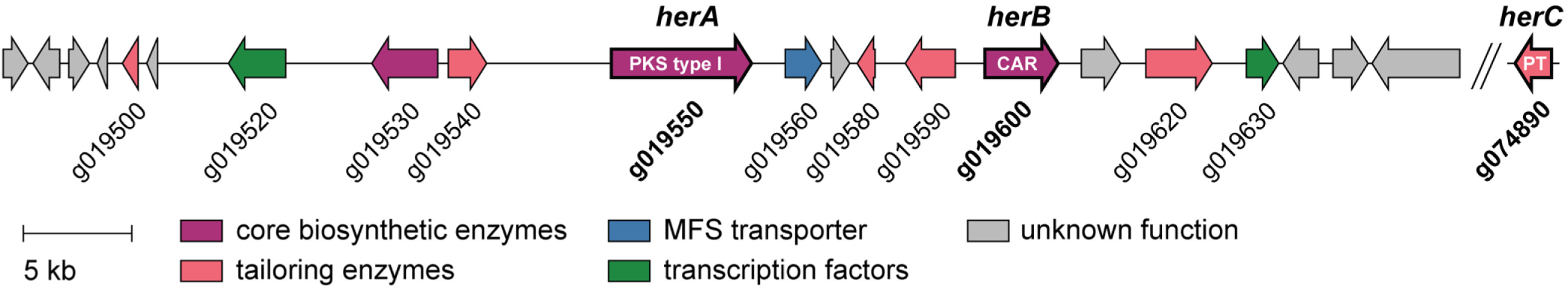
Hericenones BGC from *H. erinaceus*. The genes subjects of this study are highlighted. The putative prenyl-transferase HerC is not part of the cluster, and it was identified via a homology-based search strategy. The putative functions of the BGC genes are shown in Table 1.

**Table 1.** Predicted function of putative hericenone BGC genes. For *herA*, *herB*, and *herC*, homology to the corresponding proteins from meroterpenoid-producing *stb* BGC (MIBiG no. BGC0001390) [34] and *asc* BGCs (MIBiG no. BGC0001923 and BGC0001924) [35] is shown.

| Gene | Predicted function | *stb* BGC homolog (aa % identity; % similarity) | *asc* BGC homolog (aa % identity; % similarity) |
| --- | --- | --- | --- |
| g019500 | Flavoprotein |  |  |
| g019520 | Fungal specific transcription factor |  |  |
| g019530 | NRPS-like reductase |  |  |
| g019540 | Aldehyde dehydrogenase |  |  |
| **g019540_herA** | **Type I PKS** | BAV19379.1_StbA (24.89; 41.98) | BBF25315.1_AscC (25.02; 41.92) |
| g019560 | MFS 1 family transporter |  |  |
| g019580 | short-chain dehydrogenase/reductase SDR |  |  |
| g019590 | Monooxygenase FAD-binding |  |  |
| **g019600_herB** | **Carboxylic acid reductase** | BAV19380.1_StbB (31.71; 50.75) | BBF25314.1_AscB (32.10; 50.13) |
| g019620 | Phenylalanine-specific permease |  |  |
| g019630 | Serine/threonine protein kinase |  |  |
| **g074890_herC** | **Prenyltransferase** | BAV19381.1_StbC (25.79; 44.99) | BBF25313.1_AscA (26.52; 41.44) |

Interestingly, upon further inspection of the BGC, we did not find a prenyltransferase-encoding gene. It is common in basidiomycete fungi that enzymes underlying a specific biosynthetic pathway are encoded on different loci in the genome [10]. Thus, we performed an additional structural annotation of the *H. erinaceus* 0605 genome using GenSAS v6.0 [39] (additional file 3) and carried out a homology-based search within its proteome using the StbC and AscA PTs as queries. We identified gene g074890—on a different contig—as the most likely PT candidate (Tables S1 and S2, additional file 2). We then proceeded with the in vivo characterization efforts. First, we cloned *herA* into the pTYargB plasmid [40], under control of the maltose-inducible *amyB* promoter, and overexpressed it in the model host *A. oryzae* NSAR1 [41]. Following cultivation on solid medium, we analyzed extra- and intra-cellular metabolites via liquid chromatography-mass spectrometry (LC-MS) analysis and detected a prominent new peak (**1**) in the fungal extracts with an m/z of 167 (ESI-), corresponding to the expected value for orsellinic acid (Fig. 3a, Fig. S1). High-resolution tandem mass spectrometry (HRMS2) analysis of the same sample confirmed this observation (Fig. 3b).

**Figure 3.**
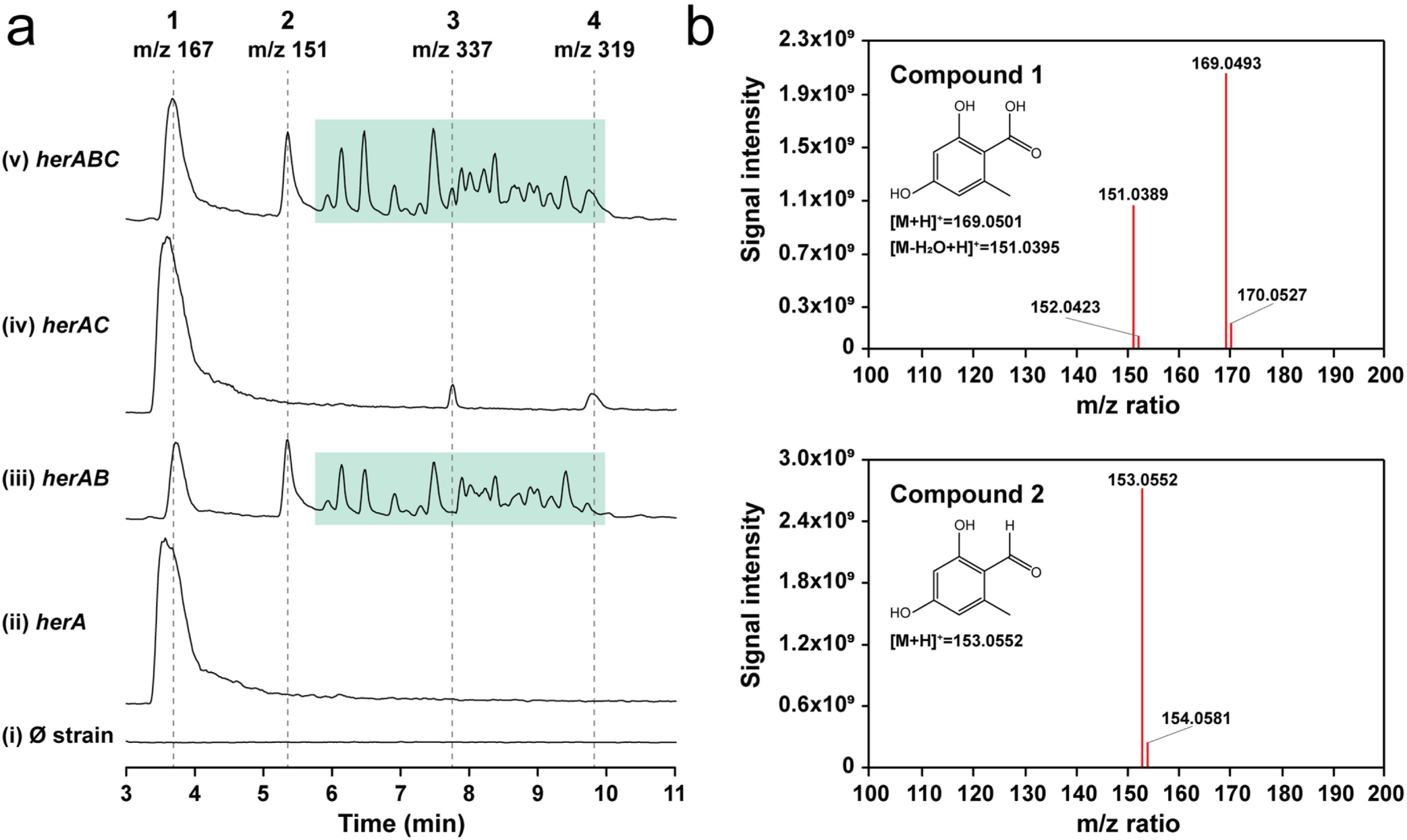
Heterologous production of orsellinic acid and orsellinic aldehyde in *A. oryzae* NSAR1. **(a)** Extracted ion chromatograms (ESI-) of fungal extracts displaying m/z 167 (**1**), 151 (**2**), 337 (**3**), 319 (**4**) and a series of m/z values ranging between 259 and 983 detected only in *herA* and *herB* co-expression strains (additional file 3), highlighted by a green-shaded box. (i) *A. oryzae* NSAR1, background strain; (ii)-(v) *her* overexpression strains. **(b-c)** HRMS2 spectra of orsellinic acid (**1**) and orsellinic aldehyde (**2**) in the fungal extracts. Spectra were recorded in positive ionization mode.

Around the same time, Han and coworkers [41] also reported the successful expression of *herA* in *A. oryzae*, which was confirmed to be an orsellinic acid synthase. Interestingly, the sequence they published was 155 nt shorter than the one we obtained via PCR. When we examined the sequence of *herA* that was predicted by GenSAS, we realized that this was a perfect match with the published sequence (Table S1). Thus, we hypothesize that the 155 nt—and in general, the full genome—were poorly annotated by the basic annotation tool of fungiSMASH, as opposed to the more accurate GenSAS pipeline that we applied. This prompted us to run BGC prediction on the fungal genome again, this time supplying the annotation file we generated. Indeed, the webtool predicted more BGCs on the genome (20 vs 12, additional file 2), and several features within the *her* BGC showed significant differences, including redefined gene boundaries and a previously undetected transcription factor (Fig. S2). The revised *her* BGC is shown in Fig. 2, while the putative functions of the genes are listed in Table 1. Two striking differences were the predicted size and genomic location of the NRPS-like–encoding gene 39—now labeled as g019600 (*herB*) (Fig. S2, Table S3).

To prioritize our cloning and overexpression efforts, we performed bioinformatic analyses on the two NRPS-like–encoding genes. We found that g019600 was likely the functional CAR because it shows the overall domain arrangement and the conserved sequence motif in the adenylation domain of fungal CARs, while g019530 lacks some of these important features (Fig. S3). Thus, we transformed the *herA* overexpression strain with a plasmid harboring g019600, under control of the *amyB* promoter. The resulting co-expression strain showed a slightly stunted growth and a different pigmentation compared to the background and *herA* overexpression strains (Fig. S4). When we analyzed the fungal extracts via LC-MS, we observed consumption of orsellinic acid and detected a second peak (**2**) with an m/z value of 151 (ESI-), corresponding to the expected value of orsellinic aldehyde (Fig. 3a, Fig. S1). Once again, HRMS2 analysis confirmed the chemical identity of the compound (Fig. 3b). Interestingly, we also detected several other peaks in the chromatograms, with m/z values ranging from 259 to 983 (ESI-) (green box in Fig. 3a, Fig. S1). Fungal strains that overexpress *herB* alone do not produce either orsellinic aldehyde, or any other metabolites compared to background strains (Fig. S1). From HRMS2 data (additional file 3), it appears that all these compounds are derivatives of orsellinic acid/aldehyde, since their MS2 spectra show low molecular weight MS fragments in the m/z value range of 120-170 compatible with (sub)structures of orsellinic acid. When we inspected their combined spectra (Fig. S5), we found that they show repeated mass differences of 136.0530 and 43.9900 Da, corresponding to chemical formulas of C_8_H_8_O_2_ (orsellinic acid minus 2 oxygen atoms) and CO_2_, respectively. We hypothesize that orsellinic aldehyde molecules react with orsellinic acid leading to the formation of polymers. The 43.9900 Da difference can be explained by decarboxylation of these molecules, which would happen spontaneously in solution and during MS analysis, as it does with orsellinic acid. Despite these observations, we cannot confirm our hypothesis without further chemical characterization by NMR. Furthermore, we do not know whether polymerization happens *in vivo* or after extraction in organic solvents. Nevertheless, our results confirm that HerB is the functional CAR within the BGC, and that it can act in concert with HerA to produce orsellinic aldehyde, core structure of the bioactive meroterpenoids from *H. erinaceus*.

Lastly, we proceeded to clone and overexpress the putative PT encoded by g074890—designated as *herC*—in both the *herA* and *herAB* overexpression strains. From the LC-MS analysis of the fungal extracts, we detected two new peaks (**3** and **4**), with respective m/z values of 337 and 319 (ESI-) (Fig. 3a). These compounds are only formed when *herA* and *herC* are co-expressed (Fig. 3a, Fig. S1) and independently of *herB* co-expression, indicating that they are derived from orsellinic acid. Unfortunately, neither **3** nor **4** show the expected m/z value of geranylated orsellinc acid—304 (ESI-)—which is the modification that we expected to be carried out by HerC. Based on HRMS2 analysis and its MS2 spectrum, however, we propose that compound **3** might indeed be a geranylated variant of orsellinic acid, albeit with two additional hydroxyl groups (Fig. S6). Compound **4** shows similarities with **3** in its MS2 spectrum (Fig. S7) and, based on its m/z value of 319 (ESI-), we could hypothesize that it is also a geranylated variant of orsellinic acid, but with only one additional hydroxyl group. Unfortunately, we do not see the corresponding ion in positive ionization mode (m/z 321), which makes any further hypothesis challenging. These observations need to be confirmed via isolation of **3** and **4** and NMR analysis which, given their limited abundance in the fungal extracts, will require scaling up the cultivations and developing a suitable purification procedure.

In conclusion, we employed bioinformatic analysis and heterologous expression to identify the first two steps of the biosynthetic pathway of meroterpenoids in *H. erinaceus*. Namely, we cloned and successfully overexpressed the non-reducing PKS HerA—an orsellinic acid synthase—and the CAR HerB, necessary to deliver the building block orsellinic aldehyde, which is the central core structure of hericenones, erinacerins, and hericerin (Fig. 1). With these results, we are able to link for the first time the biosynthesis of these compounds in *H. erinaceus* to a candidate BGC. We also identified and overexpressed a putative geranyltransferase located outside of the BGC, which could be involved in the third biosynthetic step, although we could not confirm its exact function yet based on HRMS2 analysis alone. It is also possible that another—yet to be identified—geranyltransferase is responsible for the decoration of orsellinic aldehyde, or that *herC* is active on another intermediate generated further along the pathway. Lastly, several genes found in the cluster are predicted to encode tailoring enzymes—including a flavoprotein, a monooxygenase, an SDR and an aldehyde dehydrogenase—which are all likely required to functionalize the orsellinic core structure and generate the different meroterpenoids. Efforts to reconstitute the full BGC and confirm the function of these tailoring enzymes are ongoing. Overall, our work provides a first look into the biosynthesis of bioactive meroterpenoids from the medicinal mushroom *H. erinaceus*, enabling future endeavors to produce these molecules in heterologous hosts and exploit them for pharmaceutical applications.

## EXPERIMENTAL SECTION

### Bioinformatic analyses

Structural annotation of *H. erinaceus* strain 0605 (NCBI acc. no. GCA_016906435.1) was carried out using the Genome Sequence Annotation Server (GenSAS) v6.0 [39]. Standard settings were used unless otherwise mentioned. In brief, low complexity regions and repeats were masked using RepeatModeler v2.0.3 and RepeatMasker v4.1.1 [42], setting the DNA source to ‘Fungi’. The generated masked consensus sequence was used for ab initio gene prediction using the following tools: (I) Augustus v3.4.0 [43], selecting *Coprinus cinereus* (*Coprinopsis cinerea*) as a trained organism; (II) GeneMarkES v4.48 [44]. The NCBI reference transcript and protein databases for Fungi were used for homology-based prediction, using the tools (III) blastn v2.12.0 [45]and (IV) DIAMOND v2.0.11 [46], respectively. Lastly, EvidenceModeler v1.1.1 [47] was used on the above-mentioned predictions to generate the final consensus model, weighted as follows: (I)–five, (II)-five, (III)-ten, (IV)-ten.

Secondary metabolite biosynthetic gene clusters in the genome of *H. erinaceus* were predicted using the fungal suite of antiSMASH web server v7.0 with default settings. The assembly (FASTA) file was used as only input with the first prediction, and later in combination with the annotation file (GFF3, additional file 4) generated with the GenSAS pipeline.

For the identification of the putative PT-encoding genes, the sequences of enzymes StbC (UniProt acc. no. A0A193PS58) [48] and AscA (UniProt acc. no. A0A455R413) [49] were retrieved from the UniProt database [50], and submitted to phmmer (HMMER v3.3.2) [51] to search against the total proteome of *H. erinaceus* 0605. The cutoff was set at E-value of 0.01 (additional file 2). The Sequence Manipulation Suite [52] was used to calculate identity (Ident tool) and similarity (Sim tool) percentages between HerA, HerB, and HerC, and homologous proteins from the *stb* and *asc* BGCs.

To analyze the domain architecture of candidate CAR genes g019530 and g019600, their amino acid sequences were submitted to the HMMER v3.3.2 web-tool (Fig. S3a). The multiple sequence alignment analysis between the amino acid sequences of known fungal aromatic CARs—retrieved from the UniProt database and described previously [33]—and the sequences of g019530 and g019600 to identify conserved motifs, was performed with MEGA 11 [53] using the MUSCLE algorithm [54] with default settings, and visualized in Jalview [55].

### Fungal material

*H. erinaceus* CBS 302.89 (reisolated from an infected culture originating from Taiwan) was obtained from the Westerdijk Institute strain collection (Utrecht, The Netherlands). The fungus was routinely maintained on malt extract agar (MEA: malt etraxct 30 g/L; peptone 5 g/L; microagar 15 g/L in ddH_2_O), incubated at 20 °C in the dark. For extraction of genomic DNA, *H. erinaceus* was grown in liquid malt extract medium (as MEA, without agar) incubated in static conditions at 20 °C in the dark for 14 days. The biomass was subsequently harvested and freeze-dried, and genomic DNA was extracted with the Nucleospin Microbial DNA kit (Qiagen, Venlo, the Netherlands) as described previously [56].

*A. oryzae* NSAR1 (ΔargB, adeA-, sC-, and niaD-) [41] was kindly provided by Prof. Jun-ichi Maruyama from the University of Tokyo, Japan. The fungus was cultivated on Dextrose-Peptone-Yeast extract (DPY) agar plates for routine passages (20 g/L glucose; 10 g/L peptone; 5 g/L yeast extract; 0.5 g/L MgSO_4_ · 7H_2_O; 5 g/L KH_2_PO_4_; microagar 15 g/L in ddH_2_O; pH 5.5), and on DPY-KCl agar plates (5 g/L glucose; 10 g/L peptone; 5 g/L yeast extract; 0.5 g/L MgSO_4_ · 7H_2_O; 5 g/L KH_2_PO_4_; 45 g/L KCl; 1 mL/L Hutner’s trace element solution [57]; microagar 15 g/L in ddH_2_O; pH 5.5) to induce sporulation. The plates were incubated at 30 °C in the dark for 5-7 days.

### Amplification and cloning of *her* BGC genes

The integrative expression vectors pTYargB-*eGFPac*, pTYadeA-*eGFPac*, and pTYsC-*eGFPac* were kindly provided by Dr. Colin Lazarus from the University of Bristol, UK. These were digested with FastDigest NotI and PacI and dephosphorylated with FastAP alkaline phosphatase (Thermo Fisher Scientific, Waltham, MA, USA). The ready-to-use vector fragments were subsequently separated from the *eGFPac* inserts by gel purification using the QIAquick Gel Extraction Kit (Qiagen, Venlo, the Netherlands). The *herA*, *herB*, and *herC* genes were amplified from the genomic DNA of *H. erinaceus* CBS 302.89 and cloned by sticky-end ligation in the inducible amyB promoter of the pTYargB, pTYadeA, and pTYsC vectors, respectively (Fig. S8). PCR reactions were carried out using 2x Q5 PCR master mix (New England Biolabs, Ipswich, MA, USA) and 1 µL of template (∼10 ng genomic DNA) in a total volume of 25 µL, according to manufacturer’s instructions. Primers and other gene-specific parameters are listed in Table S4. Routine procedures were used for transforming chemically competent *E. coli* DH10β cells with the assembled constructs. Positive clones were selected on LB agar plates supplemented with ampicillin 100 µg/mL. Direct-colony PCR was used to pick positive transformants, and the corresponding plasmids extracted using QIAprep Spin Miniprep Kit (Qiagen, Venlo, the Netherlands) were sent to Macrogen Europe (Amsterdam, the Netherlands) for verification by Sanger sequencing.

### Genetic transformation of *A. oryzae* NSAR1

Protoplasts of *A. oryzae* NSAR1 were obtained from spore suspensions and genetically transformed as previously described [58]. To generate overexpression strains of single genes, protoplasts were mixed with 1–2 µg (max 10 µL) of pTYargB-*herA*, pTYadeA-*herB*, or pTYsC-*herC*. For co-expression of multiple genes, protoplasts were prepared from the *herA* overexpression strain and transformed with either pTYadeA-*herB* or pTYsC-*herC* alone (double transformants), or with both plasmids (triple transformant). In all cases, the total amount of DNA used was approximately 1-2 µg (max 10 µL volume). Lastly, the protoplasts were regenerated on selective media at 30 °C in the dark, for 5 days. Next, individual colonies were picked and transferred to fresh DPY-KCl plates for sporulation and isolation of genetically pure clones. For each strain, 3 individual clones were selected and used for the following experiments.

### Cultivation and extraction of fungal metabolites

The *her* overexpression strains and *A. oryzae* NSAR1 were inoculated with sterile cotton sticks from spore suspensions onto MPY agar plates (30 g/L maltose; 10 g/L peptone; 5 g/L yeast extract; 0.5 g/L MgSO_4_ · 7H_2_O; 5 g/L KH_2_PO_4_; microagar 15 g/L in ddH_2_O; pH 5.5), where maltose acts as inducer for the amyB promoter [59]. Next, the plates were incubated in the dark at 30 °C for 5 days until sufficient growth was observed (Fig. S4). For extraction of intra- and extracellular metabolites, the whole agar pads (medium and mycelium) were sliced into pieces of roughly 1 cm^3^ and transferred to 50 mL polypropylene tubes, then extracted once with 25 mL of 9:1 ethyl acetate–methanol (v/v) supplemented with 0.1% formic acid. Extraction was carried out in a sonication bath for 1 h. Prior to extraction, all samples were spiked with 10 µL of caffeine standard solution (10 mg/mL) to validate the extraction procedure. The organic extracts were collected in clean glass vials and dried under a gentle stream of N_2_ at room temperature. The dry residues were carefully resuspended in 1 mL of 1:1 MeOH-ultrapure water (v/v) supplemented with 0.1% formic acid by pipetting and vortexing, filtered with 0.45 μm PTFE filters into clean HPLC glass vials, and stored at -20 °C until further analysis.

### Mass spectrometry analyses

Low resolution LC-MS analysis of the fungal extracts was carried out using a Waters Acquity Arc HPLC system coupled to a 2998 PDA detector and a QDa single-quadrupole mass detector (Waters, Milford, MA, USA). A Waters XBridge BEH C18 reversed-phase column was applied for separation (50 mm × 2.1 mm I.D., 3.5 μm, 130 Å particles) which was maintained at 40 °C. The mobile phase consisted of a gradient of solution A (0.1% formic acid in ultrapure water) and solution B (0.1% formic acid in acetonitrile). A split gradient was used: 0–2 min 5% B, 2–10 min linear increase to 50% B, 10-15 min linear increase to 90% B, 15-17 min held at 90% B, 17–17.01 min decrease to 5% B, and 17.01-20 min held at 5% B. The injection volume was 2 µL, and the flow rate was set to 0.5 mL/min. MS analysis was carried out in both negative and positive ionization modes (ESI), with the following parameters: probe temperature of 600 °C; capillary voltage of ± 1.0 kV; cone voltage of ± 15 V; scan range 100–1250 m/z. Data obtained via these experiments was analyzed using the proprietary software MassLynx.

HRMS2 analyses of fungal extracts were carried out using a Shimadzu Nexera X2 high performance liquid chromatography (HPLC) system with a binary LC20ADXR pump coupled to a Thermo Scientific Q Exactive plus hybrid quadrupole-orbitrap mass spectrometer (Thermo Fisher Scientific, Waltham, MA, USA). A Kinetex EVO C18 reversed-phase column was applied for HPLC separations (100 mm × 2.1 mm I.D., 2.6 μm, 100 Å particles) (Phenomenex, Torrance, CA, USA), which was maintained at 40 °C. The mobile phase consisted of a gradient of solution A (0.1% formic acid in ultrapure water) and solution B (0.1% formic acid in acetonitrile). A linear gradient was used: 0–2 min 5% B, 2–21 min linear increase to 50% B, 21–27.5 min linear increase to 90% B, 27.5-30 min held at 90% B, 30–30.5 min decrease to 5% B, and 30.5-40 min held at 5% B. The injection volume was 2 µL, and the flow rate was set to 0.4 mL/min. MS and MS/MS analyses were performed with heated electrospray ionization (HESI) in positive mode at a spray voltage of 3.5 kV, and sheath and auxiliary gas flow set at 47.5 and 11.25, respectively. The ion transfer tube temperature was 256.25°C. Spectra were acquired in data-dependent mode with a survey scan at m/z 100–1500 at a resolution of 70,000, followed by MS/MS fragmentation of the top 5 precursor ions at a resolution of 17,500. A normalized collision energy of 30 was used for fragmentation, and fragmented precursor ions were dynamically excluded for 4 s. MZmine 3 [60] was used to analyze and export chromatograms and MS spectra obtained in these experiments.

## ASSOCIATED CONTENT

**Additional file 1.** Supplementary figures and tables for molecular biology procedures, BGC prediction and gene annotation, and mass spectrometry analysis (PDF).

**Additional file 2.** BGC predictions in the genome of *H. erinaceus* 0605 by fungiSMASH using genomic fasta file (GeneBank acc. no. GCA_016906435.1) with or without gene annotations (sheet 1). Output of pHMMER-based search of putative prenyltransferase (sheet 2).

**Additional file 3.** HRMS and MS2 spectra of unidentified compounds detected in *herAB* overexpression strains (PDF).

**Additional file 4.** Annotation file (GFF3 format, compressed) for *H. erinaceus* 0605 assembly (GeneBank acc. no. GCA_016906435.1) (ZIP).

## AUTHOR INFORMATION

### Authors

**F. Poon** – Department of Chemical and Pharmaceutical Biology, Groningen Research Institute of Pharmacy, University of Groningen, 9713 AV Groningen, The Netherlands.

### Author Contributions

The manuscript was written through contributions of all authors. All authors have given approval to the final version of the manuscript.

### Notes

The authors declare no competing financial interests.

## ACKNOWLEDGMENTS

The authors are grateful to the staff of the Interfaculty Mass Spectrometry Center of the University of Groningen and University Medical Center Groningen for their services in HRMS and MS/MS analysis. We would also like to thank Prof. Jun-ichi Maruyama from the University of Tokyo (Tokyo, Japan) for the *A. oryzae* NSAR1 strain; Dr. Colin Lazarus from the University of Bristol (Bristol, UK) for the fungal pTYxxx vectors; and Jesper Martens for performing part of the molecular biology work.

