## Additional file 1 for "Identification and partial reconstitution of the biosynthetic pathway of bioactive meroterpenoids from *Hericium erinaceus* (Lion’s Mane mushroom)"

**Contents**

**Supplementary tables** 2

**Supplementary figures** 8

**Supplementary references** 14

**Table S1. DNA and protein sequences of the three genes cloned in this work.**

| Gene name | Nucleotide sequence (5’ → 3’)[a] | Predicted aa sequence[b] |
|---|---|---|
| *herA* | atgtcctccatcgctgatacgcagcatttcaatgtcccggtcttcgcaggccacggtacaacggccatcaacacacc | MSSIADTQHFNVPVFAGHGTTAI |
| (g019550)[c] | ccagacgcgtgagcgcgcccttcgcgatgcttcctcgccatcaggctctattctactctcctcttgctttgatagtttcc | NTPQTRERALRDASSPSGSILLSS |
| | aagaagagctcgctacattcactgatgaggagcgcaaggctgcaggcgtcgaagctggggacttcgataagcctg | CFDSFQEELATFTDEERKAAGVE |
| | agtctctcctctctctatctcaggagcgctacctctccaatccagtcatctctgggatcacgctcttcctcatccaaac | AGDFDKPESLLSLSQERYLSNPVI |
| | gctgaggtacctctccttcgtcgagtcctactcttcgccgtcgtctccgcgcttcgcggccatcctctcccagaacctc | SGITLFLIQTLRYLSFVESYSSPSS |
| | gcacaccagcttggtgttcttggattttcttctggtattctcccagcatcagtcgtcggcacctcagtttcggcattgga | PRFAAILSQNLAHQLGVLGFSSGI |
| | gttcatctcaaacgctgtcgaggcattccgcctcgcgttctggatcggagttcgcgcccaactttatcgcgtcgcagc | LPASVVGTSVSALEFISNAVEAFR |
| | ctttgaatccgcaaatagtctcggagatgatgcggcgctcccatggagccttgtgttcctggggattgggcggcaag | LAFWIGVRAQLYRVAAFESANSL |
| | aagcagatgaggcagtgcgcaagtaccgtgaatcggtgagtccctcgcgttctagcctcgcgttgtgatgctgaca | GDDAALPWSLVFLGIGRQEADE |
| | acccttcgatgcgcgtctgttatgcccctgcttatctcgattcattcctgcatagaatgaaaatgcagagtctctgcac | AVRKYRESNENAESLHVTAVMD |
| | gtgaccgctgtgatggatgagacttgcgtcaccatatctggacgtcctgacgttctcgctgccttcgcgtcccgactt | ETCVTISGRPDVLAAFASRLPTSA |
| | ccgacttctgcgcctccgcacaagacgaccgtagatacccttttatcactccccgattcacacaggcacgacacgcg | PPHKTTVDTLYHSPIHTGTTRDL |
| | atctggtcctggaagatgtgtcccgccgtgggattcagttcccttcgttctcggacatcaagatccccatccgctcca | VLEDVSRRGIQFPSFSDIKIPIRSM |
| | tgcacaccggcgagctgctcgattcatctcgcgagggatcgttcgtagaggcggtcgtcgatatggtcctcacgca | HTGELLDSSREGSFVEAVVDMVL |
| | gccagtgaactgggatcgcgtcgtcagttcgcttgtgtcagccgcgccagagggcgaagccgtgcgcttgatcaac | TQPVNWDRVVSSLVSAAPEGEA |
| | gtcgggcctggtgctgggcttactcgcagcatggagcgggcattccccacgcgcaatgcattgtctgtagatctgac | VRLINVGPGAGLTRSMERAFPTR |
| | tgccccggagaagaattccgccgcgaacaagacgtctcccgttcaggagccgattgctattgttggcatggcggtt | NALSVDLTAPEKNSAANKTSPV |
| | aacatgcccggagcaccgagcgtcgagaagctgtgggaggtgctcgagaagggcatcaacactattgcggaggt | QEPIAIVGMAVNMPGAPSVEKL |
| | aagtgttcttttcctagactccctggacgtacgatctcatagtcctgtcaatcactccagatccctgagcaccggttca | WEVLEKGINTIAEIPEHRFKVSD |
| | aggtctcagactacaacaaccccgccgatgccaagagtgcccggtcgatgaaggcacatacgggtaacttcctcg | YNNPADAKSARSMKAHTGNFLD |
| | acgatcctgatgcatttgacaacaagttcttcaagatctcgccgcgtgaggcacgcagcatggatcctcaaggccg | DPDAFDNKFFKISPREARSMDP |
| | cgtgctgcttcacaccgcgtacgaggcgctggaggattcaggctatgttcccaacgccacgccgacattccagccg | QGRVLLHTAYEALEDSGYVPNA |
| | gactcgttcgggtgctacatcggttgcgccacgggcgattacgttgagaatctccgaaacgatattgacgtatatta | TPTFQPDSFGCYIGCATGDYVEN |
| | cagcactggtgagagacaaatgaagccaatttcttacctcaactaacgcattttctggataggcactctgcgcgcct | LRNDIDVYYSTGTLRAFLSGRISY |
| | tcctcagtggtcggatttcatacgctatgaaacttagcggcccatctgtggtcattgatactgcgtgctcgtcttcgat | AMKLSGPSVVIDTACSSSIIAVYQ |
| | catcgccgtgtaccaggcgtgtcgtgcactcatgaacggggactgcactgcagccatggcaggcggtgtcaacgt | ACRALMNGDCTAAMAGGVNVIA |
| | gatcgctgctcccgacgtacgtacatttaatcgaaatttctaccggctggactgattgtgcggcgtatcagatgttca | APDMFMGLDRGHFLSPTGQCKA |
| | tgggtctagatcgtggtcatttcctcagccctacgggccagtgcaaggcgttcgatgactctgccgatggatactctc | FDDSADGYSRSEGCGIFVLKRLS |
| | gaagcgaaggctgtggaattttcgtcctgaagcgactgtcagatgccatcgctgaggacgacaacatccttggcgt | DAIAEDDNILGVIRSIEVNQSGLA |
| | catccgcagcattgaggttaaccagagcggtctcgcgagctccatcactcatccgcattcgccgacgcagcagatc | SSITHPHSPTQQILFKKALEKSGI |
| | cttttcaagaaggcgctcgagaagtctggcattgatgcgcgtcgcatcaacgtcgtcgaagctcacggcactggca | DARRINVVEAHGTGTQAGDPNE |
| | cgcaggctggcgatcccaacgagctcgatagtattcgcggcgtctttgctgtcggccgtacccccgcgaacccgctt | LDSIRGVFAVGRTPANPLHITSV |
| | cacatcacgtccgtgaaggccaacatcgggcatctcgaggctgcgtcgggctccgcaggcctggcaaagcttctgc | KANIGHLEAASGSAGLAKLLLML |
| | tcatgcttaagcaccgcacgattcccgcccagatctcgttgaagaatctgaacccgaagatcgtggctctcgagaa | KHRTIPAQISLKNLNPKIVALEK |
| | ggatcacacggtcatcgacagggagcacgcgccttggaacccgtcggaagagggactgacgagaattgccatgc | DHTVIDREHAPWNPSEEGLTRI |

| | | |
|---|---|---|
| | tcaacaacttcggcgccgctggttccaatggggctctgttgttggaggaatatgtcccggcgggctcgaaggcacct | AMLNNFGAAGSNGALLLEEYVP |
| | gaggtcgagtctgctgcggccttcatcgttggtctgtcagcgaagacggacgaggcactcaatgccctgcgcatgc | AGSKAPEVESAAAFIVGLSAKTD |
| | gttacatcgagtggctcggcgatgccaagaatgcatccatctccctcgccgatttcgcatacactgctacagctcgc | EALNALRMRYIEWLGDAKNASIS |
| | aggcagttgtacggccagcgtctcgctgtctctgcaggcaccaaggaggagcttgtcgagaagttgagagccgcg | LADFAYTATARRQLYGQRLAVS |
| | tcgcaggtcgctgtgactgagcggccagcgaaagttgctttcgtcttctctggccagggcagccagtacctcgggat | AGTKEELVEKLRAASQVAVTERP |
| | gggatctgccctctacaagaccgtgcctctcttcaagcgtaccgtcgacgagtgccacgctatcttaactgcgtccg | AKVAFVFSGQGSQYLGMGSALYK |
| | gcttcccgggcgttctcgccatcatcaatcctgctggtgagaccagtggtcttactcagctggaggagttcgaggca | TVPLFKRTVDECHAILTASGFPG |
| | tatcaggcggcgatcttctcccttgagtacgcgctcgccaagttgtggatgtcatggggactggttcccgaggtggt | VLAIINPAGETSGLTQLEEFEAYQ |
| | cgtaggccacaggtacgttcggtttcccgctcgaattgacagtattctgaccgtctatctgtagtttgggcgagtacg | AAIFSLEYALAKLWMSWGLVPE |
| | cagcgcaggtcatcgctggcgtcttgacgctcaagggcgcgctcactctcattgccaaccgcgtccgcttcatggtc | VVVGHSLGEYAAQVIAGVLTLKG |
| | agcaagtgcgccgtggagacgaccggcatgatcgcgataaaccagggctccgaggcggttgcgaagctgctggc | ALTLIANRVRFMVSKCAVETTG |
| | agcttcgatggacttccccgacacttctgttgcatgctttaacagcaacactgactgcgtcgtttcgggccctattcct | MIAINQGSEAVAKLLAASMDFPD |
| | cagctcaaggcactgaaggcacacctcgacagtgaagtacgctgcaagaacgtcctcttgacggtgccgttcggg | TSVACFNSNTDCVVSGPIPQLKA |
| | taccatagctccgccatgcaccccttctcgacgacctttccactattgctaagcatgttaccatccgcgcgcccact | LKAHLDSEVRCKNVLLTVPFGYH |
| | attccgattatctccaatgtcaccggtgaggtcgttatgcccggcgatgagggcgtattcgactcggagtactactct | SSAMHPLLDDLSTIAKHVTIRAP |
| | cgccactgcgcgcagccagtactcttcgagaagggcctcacttcactcgcggctattccggagcttgcgaacatcg | TIPIISNVTGEVVMPGDEGVFDSE |
| | acgcatggatcgagattggtccccacagcatcacgcttcccatgttcaaggttcacccgtcgatctcgaagagcacc | YYSRHCAQPVLFEKGLTSLAAIP |
| | atgctcttgggctccctgaagaagaaccaggacccgtgggcaatcgtctcttcgactttggctcagctgtacacgtc | ELANIDAWIEIGPHSITLPMFKV |
| | cccgattcaactcaggtggcgtgaggtctttgcccatgtctcttctccttcgacgctttccctcccgtcgtaccccttca | HPSISKSTMLLGSLKKNQDPWAI |
| | cgaagtccaagttctgggtctcgttcaaggaggaggcacctggcgcggttgtatcggcgtccacctccgtcccgctc | VSSTLAQLYTSPIQLRWREVFAH |
| | gcgaagcacgtcgatttagtcaataacttctcgatgctatactcctgggcccagttcccgtcggcggcgaaccagcg | VSSPSTLSLPSYPFTKSKFWVSFK |
| | cgtcgcaattttcgagacgcccatctcgcagctcgggaagtccatcaagggccacagcgtcggagaccacccgct | EEAPGAVVSASTSVPLAKHVDLV |
| | ctgccccgcgtccgtgtaccacgagctcgcacttgcaggtatcgagatggctaggtcccatctgcaccttaagattg | NNFSMLYSWAQFPSAANQRVAI |
| | acgactgcttcgtcatgcttcgcgacatcgactacgcgaagccgctggtctacaacgagcacgttgcccggatggt | FETPISQLGKSIKGHSVGDHPLCP |
| | caggacatcgatcacacttgatgttgatggctcgggctctttcagcgttggctcgcaggtagatggctcgcccgagg | ASVYHELALAGIEMARSHLHLKI |
| | aggtccattgcttcgggaagttcaagcaccagtcgacatcgaaggcctcaaccaagttcgctcgcgttcttcccatc | DDCFVMLRDIDYAKPLVYNEHV |
| | gtaactcgtcaaatcagctccgtctcgtcgcctaacgatggtttcgcggagacgttctctactcgcacggcatatga | ARMVRTSITLDVDGSGSFSVGSQ |
| | ggtcatcttcccgcgcgtggtcgactacgcaaaggagtaccacacgatgaagacgctcaccgtcgcctcgaacgg | VDGSPEEVHCFGKFKHQSTSKAS |
| | tatggaagcatacgcgatcgtccagctgcctcgcgatcacgaccgcagcaagtttgtcgtccaccctgtcttcatgg | TKFARVLPIVTRQISSVSSPNDGF |
| | acaccatgcttcacgttgccggcttcgtcgcgaacatgcagggcggagtcaacgatgcctacatctgcagcaaggt | AETFSTRTAYEVIFPRVVDYAKE |
| | cgacacagtgaaggcgatcccggatttgatcgacaacgacgcggagtacggcgtgctgatcagcaacgcctgggt | YHTMKTLTVASNGMEAYAIVQL |
| | cgcggatgagggcgtcatgcttgcggaggcctacgccgtccagttgaagtctccaggtaagatcgtcgcgcacctc | PRDHDRSKFVVHPVFMDTMLH |
| | aagggcatgcacttccgcaaggtgcgcctcaatagcctgaagcgcggtctcgcgatggccgcgggcactagctcg | VAGFVANMQGGVNDAYICSKVD |
| | gcgcatgctgcgccgaagcgggcggaggcgcctgtgaagaaggctgcgcctgcttccccgatgtcctccaaggtc | TVKAIPDLIDNDAEYGVLISNAW |
| | gtttcctcaatcaccttcgtcgagccgcctcgagatgcagcacctactatcgacgtcctcgcggaggtcacgcgcat | VADEGVMLAEAYAVQLKSPGKIV |
| | cgtggccgagacttgcgacatcactgcatccacgatccagcccgatggggacctcgaggcgtacggcgtggattc | AHLKGMHFRKVRLNSLKRGLAM |
| | gctgatgtccattgagatcttcaccaagatgcagagcgccttcgctagtgctgaccttgacgcgaacgtcctttcctc | AAGTSSAHAAPKRAEAPVKKAA |
| | gtgccgcaatgttgcccagatcgtcgctgaggtgtcgtccaagttctcccaggaggactctggcccatccacgccgc | PASPMSSKVVSSITFVEPPRDAA |

| | | |
|---|---|---|
| | gcacactcgtcactgacgagaagcttggcgagcccagcgtcatcgcgaacttcgacgaggtcgacgtcaagccgc<br>tgctggcgtccgtcctcggcatcggcctccaggagatcaccgacgacgcggacttcgaatcgctcggcttagactc<br>gctgacatccatcgaggcgcattcggccctgcagagcgagtactcgctgacgctgccgaccaccctgttcgagaca<br>tacacaacggcgaaggccgtcaacgccttcctcacatcgcagctgcgtcctcggggcaaggccgtcgaggtcgtta<br>aggaggccgaggtgcacccttcggactacaaggactcggacatcgcgaccgccgcgaaggtcgcgcagatcgtc<br>gacggcaatctcaacccgctcgtcaccgcgctgcgcctcgactcggttcccgtcggtgcgcagaaggcgaagaca<br>cccggccgcgcgcccttattcctgattcacgatggtagcggccttgtgaactacatccagcgcctctccccgcttgac<br>cgcgacatttggggcattcataatccccacttcatcaccagccagccgtgggagagtgtcgtgtcgatggctgcgg<br>agtactctgagttcgcgacgaagacgacctctgagcctcttatccttggtggtgagtacccctctatcgctaccttatt<br>cttgctttgttgataattgtttatctcaggctggtcgttcggtggtgtcgttgccttcgaagctgcgcgtcagctcatga<br>agaagggcgttgccgtgaagggcgtcctcctcatcgactcccctactcctctcgcccacgttccgctctccgacgcc<br>ctcctcgagtccgtcgcgaagctggatggccgcgtcgacacgcaggttggcaagctcgtgaagacgcagttccag<br>atgaactcgcgtatgctcggccgctatgacccgcttgccgcgggcgggccgtttccctcgatcgtcctcctccgctcg<br>agagagggcttcaaacccgccggcgtcgccgacgtgccgaagtggcttgcggacaggagcgatgcgcagctggc<br>catatccgggtgggagcgcgtcgtgggcacgccgatcaaggccatcgacatccccggcaaccatttccagcctttc<br>cacacctctaatgtaagtacttgctgctgtctcttgagacgtcttgtcgctgaccatagatttttagatcgaagaagtt<br>tctcgtcgcattgctgaggggtgcgcacacctcgagagccttgctgct**tga**aggccgtactgtacactattcccttct<br>tttgtatcatgcgtcatgacatccgacatgtttctcctgcatcagtagaggttcacgattcccattgtattttactgcgc<br>tgctgctttacatgctgctccttgttacgattttcacgtagtctctag | PTIDVLAEVTRIVAETCDITASTI<br>QPDGDLEAYGVDSLMSIEIFTKM<br>QSAFASADLDANVLSSCRNVAQI<br>VAEVSSKFSQEDSGPSTPRTLVT<br>DEKLGEPSVIANFDEVDVKPLLA<br>SVLGIGLQEITDDADFESLGLDSL<br>TSIEAHSALQSEYSLTLPTTLFET<br>YTTAKAVNAFLTSQLRPRGKAV<br>EVVKEAEVHPSDYKDSDIATAAK<br>VAQIVDGNLNPLVTALRLDSVPV<br>GAQKAKTPGRAPLFLIHDGSGLV<br>NYIQRLSPLDRDIWGIHNPHFITS<br>QPWESVVSMAAEYSEFATKTTS<br>EPLILGGWSFGGVVAFEAARQL<br>MKKGVAVKGVLLIDSPTPLAHVP<br>LSDALLESVAKLDGRVDTQVGKL<br>VKTQFQMNSRMLGRYDPLAAGG<br>PFPSIVLLRSREGFKPAGVADVP<br>KWLADRSDAQLAISGWERVVGT<br>PIKAIDIPGNHFQPFHTSNIEEVS<br>RRIAEGCAHLESLAA* |
| *herB*<br><br>(g019600) | atgccgattcccgtgaatactttcgttcgcccacggctcaatctgggatacggccaagacagcagcgaaggcgtca<br>actcactcccagagttgatcgccttcaatgccgagcataatcccggccatgtcttcggcttgcagattcgcgctggc<br>gagaacatttcgccttgcaaaatgacattcgcagaacttcatgccgccgttgagcgtgcgtcggcatggctcatggc<br>atctggtgccaccgcggggcggacgtcgcgggacaccaaggtcgctcccgtcgctattttactggggagcgatatt<br>ggaatcttcatctatatggctgcattattgcggattggcacgccggtaattattctgtacctttcggactatctttaaag<br>gtgagctaacgacgaactttttcaggtccttctgctttccgcccgccttaccccaatcgccatcgcacatctcatcaa<br>ggcgacatctccttcgacgatcctcataaacgctcaagtctctcgatccgccaatgagaccgttgatcttttggcgac<br>tgatgacagcgccttcaccagaccgcaattcctccatgctttgggttacgaggatttcatcagcggcgaacatcccg<br>acctgcaaaatctttccatccctcccgtctacgacgctttcaagtatgaggatctcgacgcgatcatcatgcactcat<br>ccggcaccaccggtcttcccaagcccatctaccacgcgcaggcgtaccttctcatctacgctggctgccactgcatc<br>cccgagtcaagagagccgtcgcatttcaacgtttctagcttgcctctctatcatgtgcgtccgcagttttgcgcttttcc<br>acccgattctgacgttcaatattccacagggcttcggattattagcaccgtcgctttccctctctatcggcctcccattc<br>gtcttacccccctgcctctatcattccgacggcacggacggtattgaacagtctagagttaacgcgcgcacggtctat<br>gctgtccgtcccttccatattggaggatattgtcagactgcctggtgccgctggcttggaagcgctcaagaaactgg<br>acttcattgccatcggaggcgctcccatgaaagaagccgtcgcacaagaacttgtttcgaacggcgtcaatctgctc<br>aatcactggggtgcgtctttctgatgttcttaaaatatgtcgcagcatgttgactggcttgttcaatacaggagccac<br>ggagattggtgccatcgcacctgttcagcgcccgcctctgggatacgattggcattacctcatccctcgcactgaca | MPIPVNTFVRPRLNLGYGQDSSE<br>GVNSLPELIAFNAEHNPGHVFGL<br>QIRAGENISPCKMTFAELHAAVE<br>RASAWLMASGATAGRTSRDTKV<br>APVAILLGSDIGIFIYMAALLRIGT<br>PVLLLSARLTPIAIAHLIKATSPST<br>ILINAQVSRSANETVDLLATDDS<br>AFTRPQFLHALGYEDFISGEHPD<br>LQNLSIPPVYDAFKYEDLDAIIM<br>HSSGTTGLPKPIYHAQAYLLIYAG<br>CHCIPESREPSHFNVSSLPLYHGF<br>GLLAPSLSLSIGLPFVLPPASIIPT<br>ARTVLNSLELTRARSMLSVPSIL<br>EDIVRLPGAAGLEALKKLDFIAIG<br>GAPMKEAVAQELVSNGVNLLNH<br>WGATEIGAIAPVQRPPLGYDWH<br>YLIPRTDIGLEVIQLDDAGRTYRL |

| | | |
|---|---|---|
| | ttggcctcgaggtgatccagctcgatgacgctggccgcacttaccgccttatcgggcgggctcccggctggccgga<br>gccattcgtcgtccaagatcttctcgaagtacacccatctgatccgacgcagttcaagatccttggacgcgccgacg<br>atctgatcgtactcgccacgggcgaaaaagtccgccccacgagcatggaagctgccattgcagagcaccccgacg<br>taaaggccgtcctggcgttcggagatggccagttctcgctcggacttttggtcgagctggcgtcttcaaagtccggg<br>ctggatctttctctgcctgccaatgtggatgccgtgctggagacgatggagccccacctggagcgcggcaactcgtt<br>catggacaagcatgcaaaaatcacgaaggatatgattgtactcactcagcccgagattaagccgctcgtgcgcac<br>cgacaagggcagcctggcgcgaaaggcgacgttctttgcgttcgagaaggagatcaaagagtgctacgaacgtg<br>cggatgtggcgcgcgctgtacccttcccactctacagcgtcgacgagggtgcctctctcctgtcctcgttgcgggctc<br>ttgttgggtccaccctcggaattgatgacatcggcaccttcagcgacgagtccgatttcttcgaggctggcatggac<br>agtctgcaggccagccgtctacgtcgtgccattcagaacggcctccgcattaccaaagacatttcaggtcccgtccc<br>ggaactcgctcctgacttcgtcttcgagcactcctccgtgaaaaggatatgcaacgtcatggctcacatcatgctcg<br>ggacttatgaagccgacgccggcatggacaaggaggagcgaaggatagctgctatgggcgacatggtccaacgc<br>tacgcgaaggagcttacctggtacgccgaagacgcgcttctggctcgcgaagcaagggccgcatcttctcacaag<br>cgtgcctcgacgtcgaagtccactgtgctgttgacaggctctacgggcagtctgggttgcatgctcttggcacagct<br>cgccggcgaccccggcgtcgccaagatcatctgtttgaatcggccacagcagggcggtatcgacattcggaagcg<br>acaggcagatgcgatgaagaagcgcggcgcccatatcgacgctgaggggtgggataaggtcgtcatctacgagg<br>cggacataagccgcgcagacttcggtctcagtgacaacgaatttgaagaggtaagggaccatctctgtctctctgtc<br>ttctctgttcatttatgattctccccagctgttagaggtgacgcacatcattcacaacgcgtggcctgtcaacttcaat<br>cgcacgctggcttcgtttgattcccatgtcagagccctttgcaaccttgcgcggctatcccttctcagtgcagcgaag<br>ttccctacggatcgtcgaccgcggcgcatcttgttcgcatcgtcgatcgctgtggtcggccggttcccgctccttcatc<br>cggaggggcccttcgacgtgccagagactccccttgatgctgcgaacaccgctgagttcggataccctgaggcaa<br>aatgggtgtgcgagcgggtattgcaaacattctcagacctgtacggcaagcctggccatcggcaggaggaggccc<br>ttgtccagacgtcaagcgtccgcatcggacaaatgacgggtcctgaaggctctggcgcgtggaacgagaacgagc<br>acttccccatcattgtgcgcacctcgcagaagctcagggcattgcctgacattgacggcgtgagtatactctccctg<br>cgttgctttcaagtgactcataattgctccagtcattgtcatggatgcctgtcaaccgtgcggggtcggccatcgtgg<br>atttcctcttctcgaagaacttccgtccaatctaccacatggagaatccgtcgcgccagtcgtggtcaggtcttctag<br>agaaccttgcatcagttcttggcgatagagacgggcctctgcctgtgatcccgtacgacaagtggctgcagcgcgt<br>ccgcgacctcggctctgacccggagaagaacgtcgcgtacaaggtcatggacttcttggagaatgacttcgtgcgg<br>atggcgtccgggagcgtaatcttgggcaccgacgtcgcgaaacgcgattcactgacgatggtgaagagcgtcgca<br>ctggacaaaaagcacctggctgagtatgttgcttattggaggagtgttggtgcactgcagtag | IGRAPGWPEPFVVQDLLEVHPS<br>DPTQFKILGRADDLIVLATGEKV<br>RPTSMEAAIAEHPDVKAVLAFG<br>DGQFSLGLLVELASSKSGLDLSLP<br>ANVDAVLETMEPHLERGNSFMD<br>KHAKITKDMIVLTQPEIKPLVRT<br>DKGSLARKATFFAFEKEIKECYE<br>RADVARAVPFPLYSVDEGASLLS<br>SLRALVGSTLGIDDIGTFSDESDF<br>FEAGMDSLQASRLRRAIQNGLRI<br>TKDISGPVPELAPDFVFEHSSVK<br>RICNVMAHIMLGTYEADAGMDK<br>EERRIAAMGDMVQRYAKELTW<br>YAEDALLAREARAASSHKRASTS<br>KSTVLLTGSTGSLGCMLLAQLAG<br>DPGVAKIICLNRPQQGGIDIRKRQ<br>ADAMKKRGAHIDAEGWDKVVIY<br>EADISRADFGLSDNEFEELLEVT<br>HIIHNAWPVNFNRTLASFDSHV<br>RALCNLARLSLLSAAKFPTDRRP<br>RRILFASSIAVVGRFPLLHPEGPF<br>DVPETPLDAANTAEFGYPEAKW<br>VCERVLQTFSDLYGKPGHRQEEA<br>LVQTSSVRIGQMTGPEGSGAWN<br>ENEHFPIIVRTSQKLRALPDIDGS<br>LSWMPVNRAGSAIVDFLFSKNF<br>RPIYHMENPSRQSWSGLLENLAS<br>VLGDRDGPLPVIPYDKWLQRVR<br>DLGSDPEKNVAYKVMDFLENDF<br>VRMASGSVILGTDVAKRDSLTM<br>VKSVALDKKHLAEYVAYWRSVG<br>ALQ* |
| *herC*<br>(g074890) | atggccaacgagaagacgcccctcagagagcccatccagctcaagaaaaagacgctgcctttcccttatggcctct<br>tccccgcatcgctgcgtccatatgctgagctcatgcggttgcacagagtacgcaccacccttctctctcctcacaggc<br>gcgtctgacatttattcatcagccttccgggaccatcatgattttctggccgtatggtatggcgtctgttcgccatcat<br>gcatcggaagctcaatggcatttttcattttagcgtttggcgcgacgatggccgcttaccagacgggcttccctgtca<br>agcagtactgggcagagcttggcaagttcctcatcgctgcgttcttcgtgcgtagcgcgggctgcaccatcaacga<br>cattgccgaccgggagtttgacgctggcgtcggtgggtccttttaaatattcttttgtccagaataatattgacgccct | MANEKTPLREPIQLKKKTLPFPY<br>GLFPASLRPYAELMRLHRPSGTI<br>MIFWPYAFGATMAAYQTGFPVK<br>QYWAELGKFLIAAFFVRSAGCTI<br>NDIADREFDAGVERTKSRPLASG<br>RVTVFAAYVFCILQWICAIFVFLP |

| | | |
|---|---|---|
| | cttggtagagaggaccaagagtcggccattggctagcggacgcgtcaccgtgtttgcagcatatgtgttctgcattc<br>tccaatggatctgcgcgatcttcgttttcttgccctacaactcgacaacgtacgtacaaccgtccgtgcttcacttggc<br>ctgctttggctgacggcgtcatgcacaccacccagaatgatcgcagcaattctgcaatgcatcccgatgtcgacgg<br>cttatccgtatatgaagcgcatcacatactggccccaggcgtggctcggtctgacgatgtcgatgggcatctttatcg<br>gctgggccgcgatcgcagagacgccgaattggctcctgctcggttcgttcatgctcggcttcgtcgcttggacactg<br>cacttcggtacgcactttctctagcctccgtagccgcgcgcttactgacgcgtccattccatgcgcgcctagacacca<br>tatacgcctgccaagaccgcaaagacgacatcaaggtcggcgtcaagtcgaccgcagtgatcctgggtgacctcg<br>tcatcccgttcgcgatggtgtgctcgacgacgttcgtcggtgcgctcgcgttcgcggggtacctcaacggcgagacg<br>aaggcctactacttcgtcaccgtcgcgggcaccgcggcgcacttcgtgtggcagcttgcgaccgtcgacctcgagg<br>acggggatagctgcagccgtatgtatttcctcatatgtaatgacatagcgcttatctgtcaaacagtcaactttacac<br>gcaatggacaacttggatgggtcctctggggcggaatggcaatcgactatctgctcaagatgggcgtcattcagct<br>cggcgagggccggcttagcattgttgtatga | YNSTTMIAAILQCIPMSTAYPYM<br>KRITYWPQAWLGLTMSMGIFIG<br>WAAIAETPNWLLLGSFMLGFVA<br>WTLHFDTIYACQDRKDDIKVGV<br>KSTAVILGDLVIPFAMVCSTTFVG<br>ALAFAGYLNGETKAYYFVTVAGT<br>AAHFVWQLATVDLEDGDSCSLN<br>FTRNGQLGWVLWGGMAIDYLL<br>KMGVIQLGEGRLSIVV* |

a. Predicted introns are highlighted in grey.

b. Predicted aa sequences are based on structural and functional annotation performed with the GenSAS v6.0 webtools (additional file 4).

c. *herA* was cloned prior to re-annotation of the genome, as predicted by fungiSMASH (gene 34, Fig. S2), and therefore contains an extra stretch of 155 bp at its 3' end, highlighted in yellow in the nt sequence. The stop codon of the correctly annotated g019550 is underlined and bold.

### Table S2. Genomic location of the hericenone BGC and the non-clustered g074890 gene encoding a putative prenyltransferase.

| Organism & genome assembly (GenBank No.) | Scaffold/contig (GenBank No.) | BGC genes | Location of BGC genes (nt) |
|---|---|---|---|
| *Hericium erinaceus* strain 0605 (GCA_016906435.1) | contig 12 (JABWEG010000012.1) | HE-BGC4.1 (fungiSMASH) | 100,553 – 171,961 (total: 71,409 nt) |
| | contig 5 (JABWEG010000005.1) | g074890 | 960,050 – 961,393 (total: 1,344 nt) |

**Table S3. Comparison between the NRPS-like CAR gene preliminary annotated with Prodigal integrated into fungiSMASH or annotated with GenSAS v6.0 pipeline prior to fungiSMASH analysis.**

| Gene ID (annotation tool) | Location on contig (nt) | Length (nt) | Predicted protein length (aa) |
|---|---|---|---|
| Gene 39 (fungiSMASH) | 147,813 – 152,276 | 4,463 (2,889 w/o introns) | 962 |
| Gene g019600 (GenSAS v6.0) | 148,681 – 152,276 | 3,595 (3,315 w/o introns) | 1,104 |

**Table S4. Primers and PCR conditions used in this study.**

| Target | FW primer[a] (5' à 3') | RV primer[a] (5' à 3') | $T_{ann}$ | Extension time |
|---|---|---|---|---|
| *herA*[b] | *atcg*GCGGCCGCatgtcctcca tcgctgatacgca | *atcg*TTAATTAActagagact acgtgaaaatcgtaacaaggagc | 71 ˚C | 3 min 30 sec |
| *herB* | *atcg*GCGGCCGCatgccgattc ccgtgaatactttcg | *catg*TTAATTAActactgcag tgcaccaacactcc | 70 ˚C | 2 min |
| *herC* | *ctga*GCGGCCGCatggccaac gagaagacgcc | *atcg*TTAATTAAtcatacaac aatgctaagccggcc | 70 ˚C | 35 sec |

a. 4-bp cleavage overhangs are italicized; recognition sites for restriction enzymes NotI (FW) and PacI (RV) are capitalized.

b. *herA* was amplified based on the sequence of gene 34 as predicted by fungiSMASH, which stretches 155 bp further than the reannotated g019550 (see table S1).

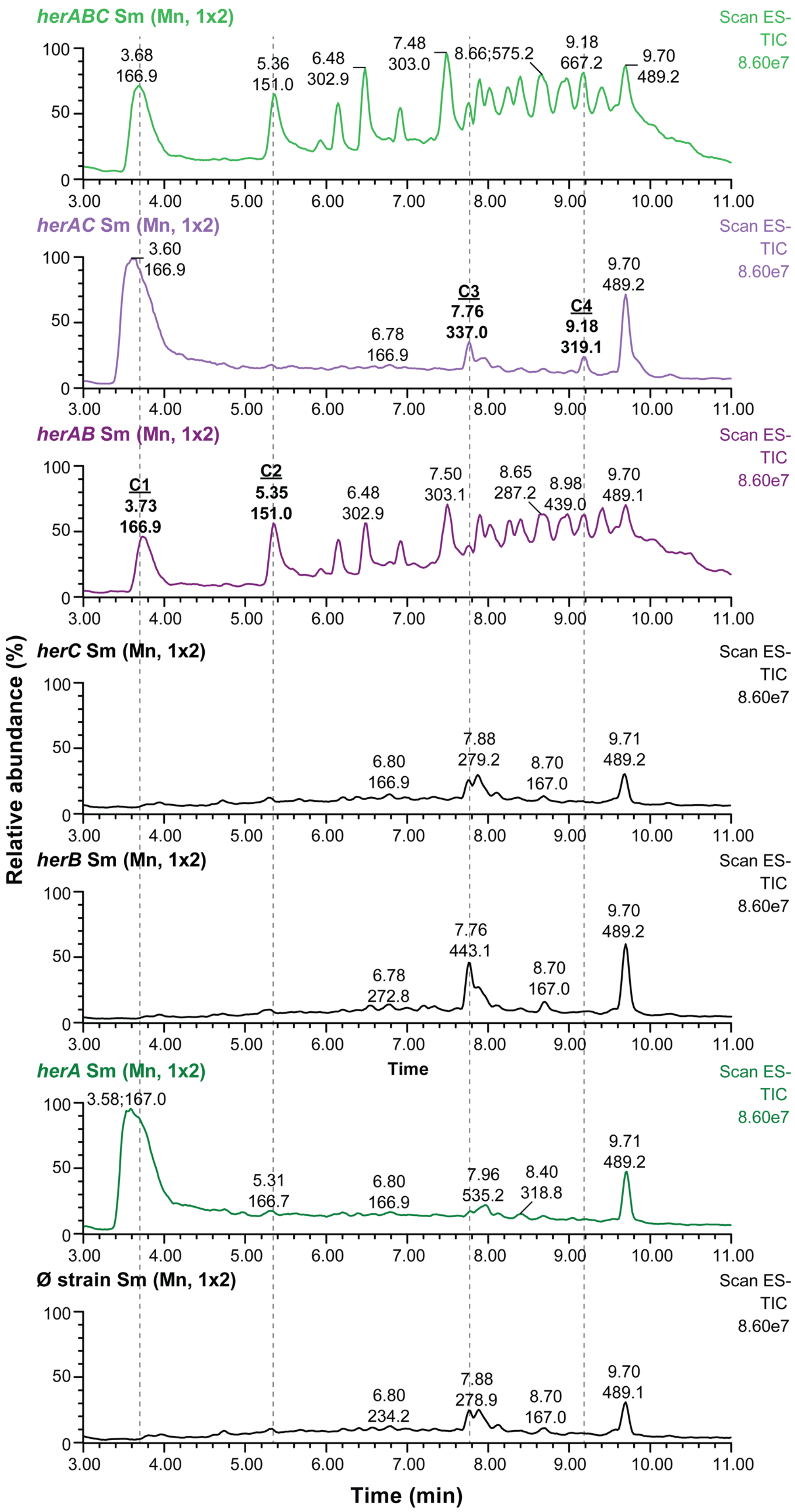


**Figure S1. Total ion chromatograms of fungal extracts from *A. oryzae* strains.** Extract of control strain *A. oryzae* NSAR1 is at the bottom. Despite small differences in signal intensity for control strains *herB* and *herC,* all the peaks are attributable to the background of the empty strain transformed with pTYsC or pTYadeA vectors. Compounds 1 to 4 are highlighted as in Figure 1. Peaks are annotated automatically by the proprietary software MassLynx v4.2 (Waters) based on the retention time at the highest point of the peak, and on the m/z value of the most abundant ion at that specific RT. Chromatograms are smoothed (window size (scans): ± 2; iterations: 1; method: mean) for visualization purposes.

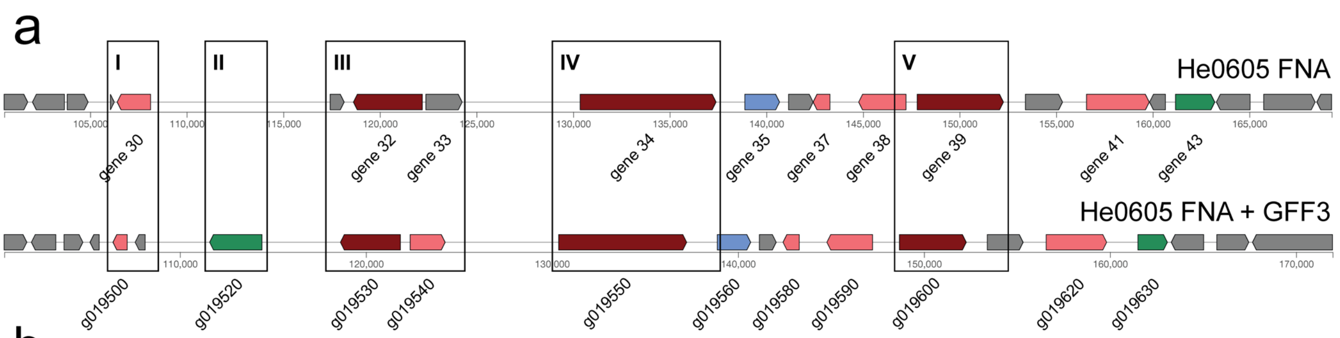


b

I - gene 30 flavoprotein + phocein-family domain (1,728 bp) → **flavoprotein g019500 (765 bp)** + phocein-family unknown g019510 (535 bp)

II - previously unannotated gene now identified: **gene 019520 (2,810 bp) putative fungal specific Zn(2)-Cys(6) transcription factor**

III - gene 32 NRPS-like (3,571 bp) → **g019530 (3,228 bp)**; gene 33 unknown (1,881 bp) → **aldehyde dehydrogenase g019540 (1,881 bp)**

IV - gene 34 PKS type I (7,033 bp) → **g019600 (6,878 bp)**

V - gene 39 NRPS-like (4,463 bp) → **g019600 (3,595 bp)**

**Figure S2. Hericenones BGC prediction by fungiSMASH. (a)** visual representation of hericenone BGC (HE-BGC4.1) as predicted by fungiSMASH v7.0 (1) using only the genomic FASTA file as input or genomic FASTA file plus annotation file generated with the web-tool GenSAS v6.0 (2). Several differences are noted, and in particular within five regions (I-V) where genes with a putative function are predicted. **(b)** key differences with respect to gene annotation/prediction in the highlighted genomic regions.

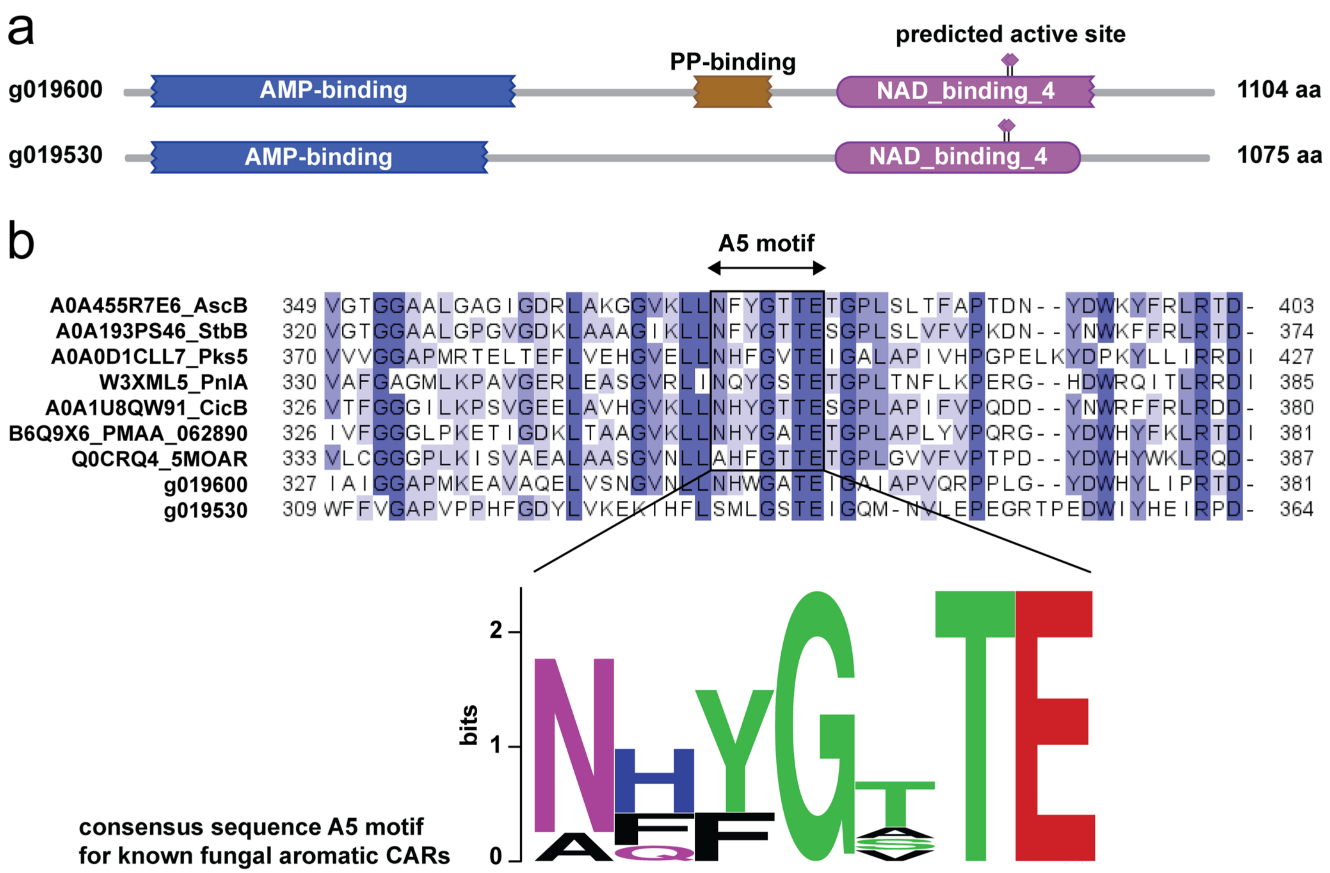


**Figure S3. Criteria for selection of CAR gene. (a)** domain prediction performed with the HMMER webserver [3,4] indicated that g019600 has all domains needed for CAR activity, whereas g019530 seems to be missing the phosphopantheteine attachment site required for the activated substrate to be transported to the reductase domain. **(b)** MSA analysis of the two candidate genes and known fungal aromatic CARs from previous studies [5] shows a high degree of conservation within the key A5 motif of the adenylation domain for g019600, but not for g019530. The sequence logo for the A5 motif was generated with the WebLogo tool [6].

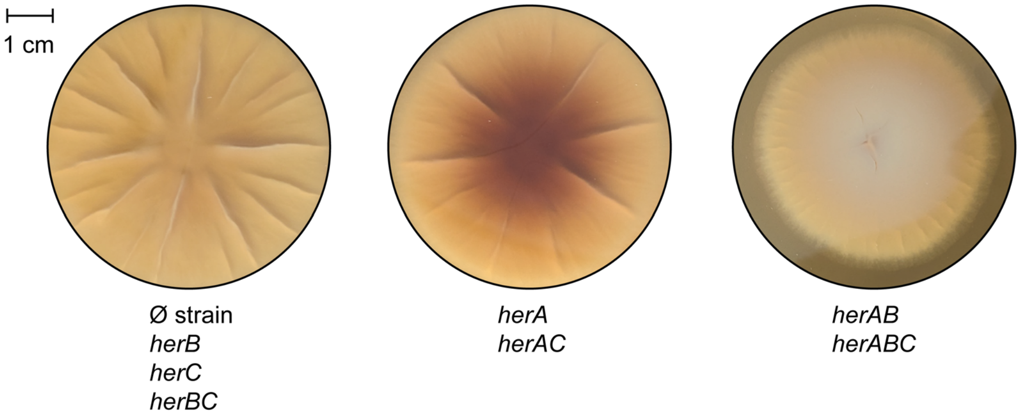


**Figure S4. Morphology of *A. oryzae* strains (reverse view).** Overexpression of *herA* leads to secretion of a dark brown pigment in the agar, likely a byproduct of orsellinic acid biosynthesis and/or degradation. Growth and appearance remain otherwise unchanged. Co-expression of *herA* and *herb* results in slower growth, and the mycelium appears velvety white (particularly on the agar side) and noticeably more compact.

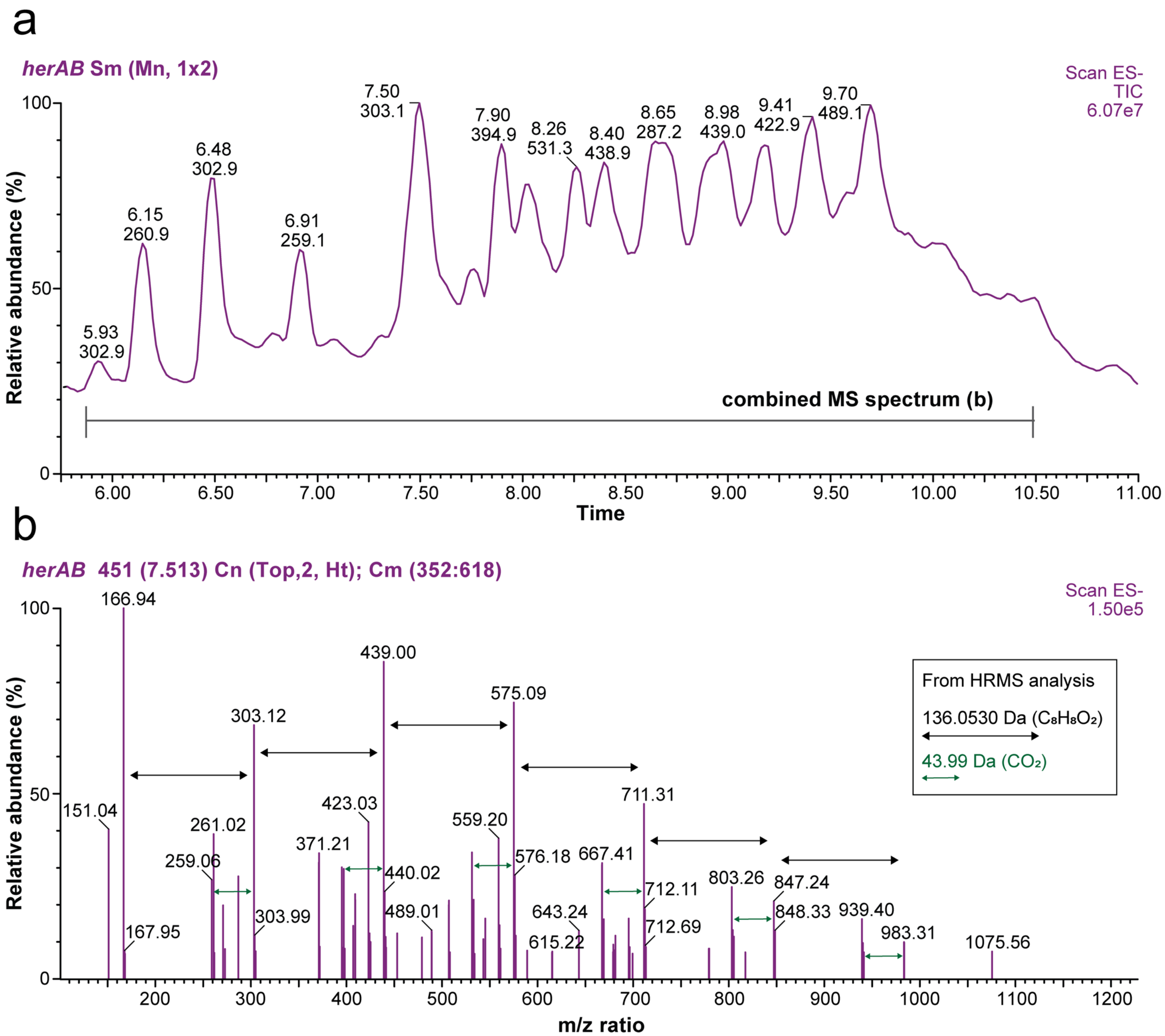


**Figure S5. LC-MS analysis of unidentified compounds from fungal extract from *herAB* overexpression strain. (a)** Chromatogram of fungal extract displaying TIC at RT range ~ 5.75 – 11.00 min, where the series of unknown peaks from the extract appears. **(b)** Analysis of the combined spectrum reveals that the most abundant ions detected show repeating units of 136 and 44 (136.0530 and 43.99, respectively, from corresponding HRMS analysis of select peaks). These are consistent with repeated additions of $C_8H_8O_2$ units—indicative of polymerization—and corresponding decarboxylation.

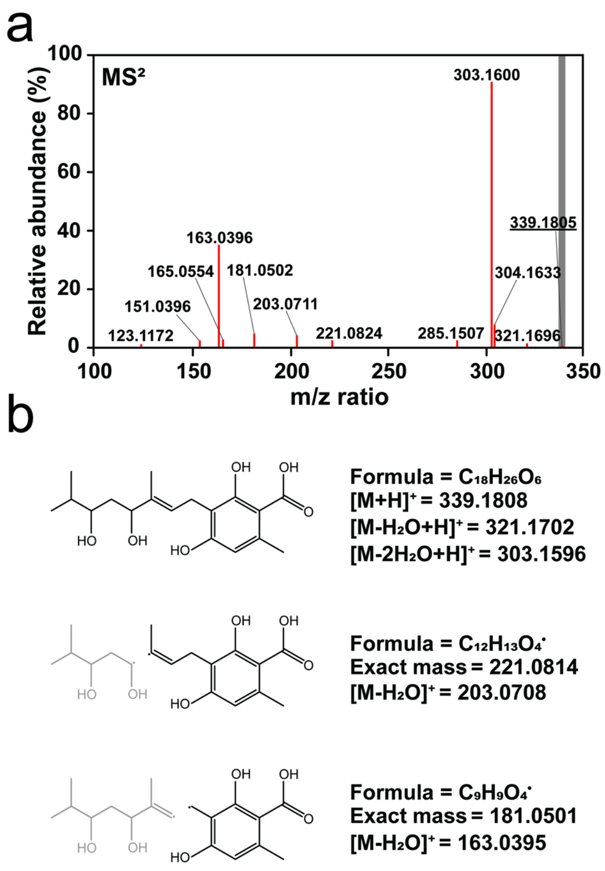


**Figure S6. HRMS-MS-based analysis of compound 3. (a)** MS2 spectrum of compound 3, showing two major fragments at m/z 303.1600 and 163.0396. **(b)** Proposed chemical structure of compound 3 and its fragments (in black) based on exact mass calculations and adduct formation. The analysis suggests that C3 might be a geranylated variant of orsellinic acid with two hydroxylations on the geranyl moiety. Chemical purification and NMR are required for confirmation.

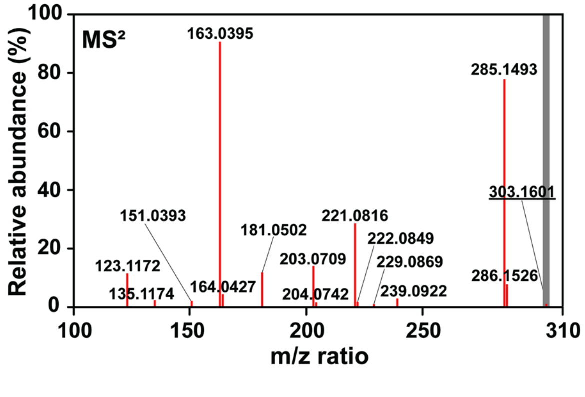


**Figure S7. MS2 spectrum of compound 4.** The spectrum shows similarity with that of compound 3, but no deductions could be made based on the m/z value of 303.1601, for the precursor ion. In low-resolution MS analysis (negative mode), we detected a m/z value of 319 $[M-H]^-$, possibly indicating a mono-hydroxylated orsellinic acid variant. We could not detect the corresponding m/z value of 321 $[M+H]^+$ in positive mode, neither with low-resolution MS nor with HRMS, thus we refrain from further propositions.

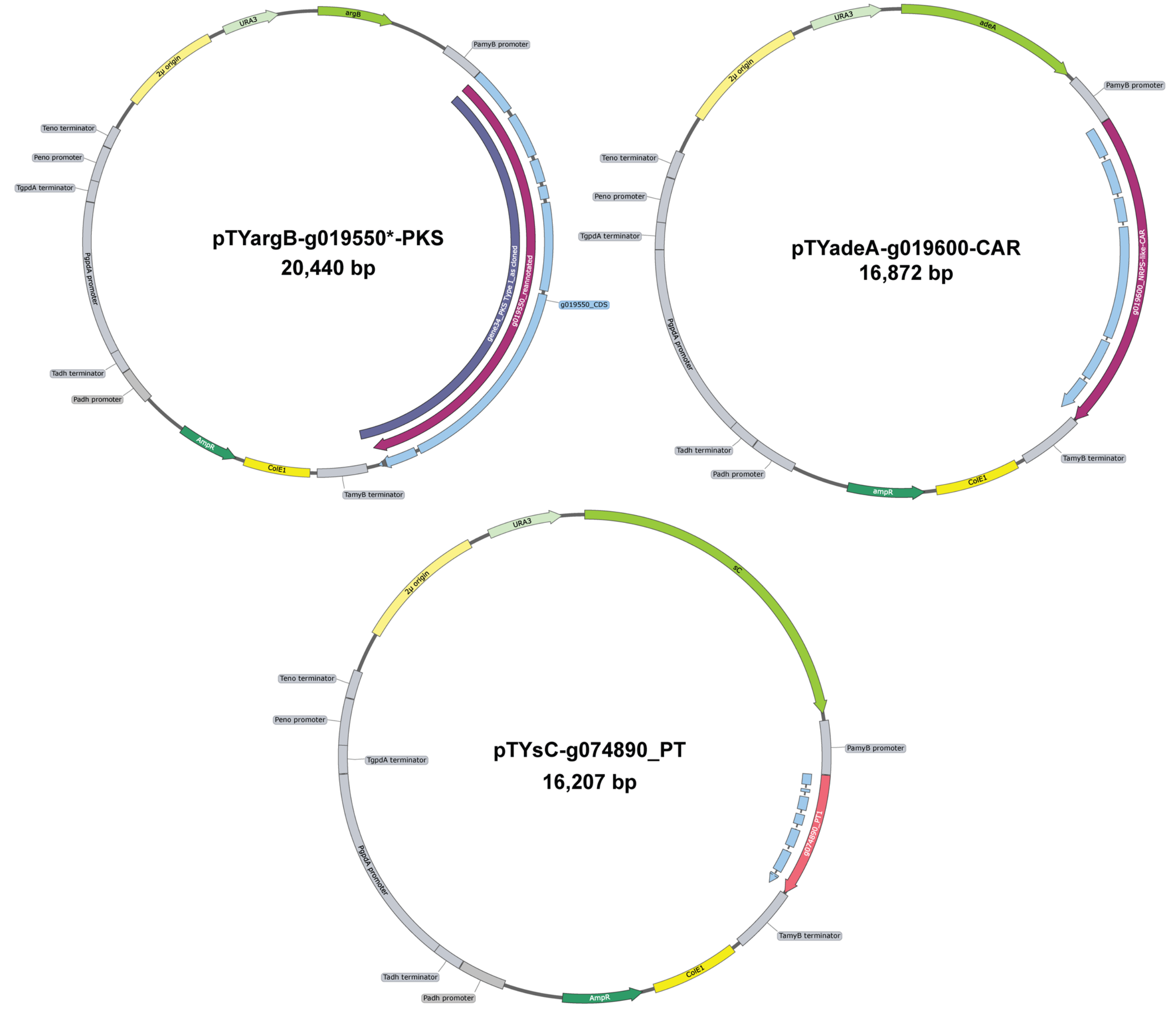


**Figure S8. Maps of *herA*, *herB*, and *herC* overexpression plasmids.** The corresponding features are displayed both as full genes (continuous arrows) or as intron-free coding sequences (interrupted arrows). *NB: g019550 was cloned prior re-annotation with GenSAS v6.0, and therefore contains an extra stretch of nt at the 3’, highlighted in Table S1. The full cloned features is annotated as a box, whereas the re-annotated gene and corresponding intron-free CDS as arrows. Plasmid maps generated with SnapGene software (www.snapgene.com).
