## Additional file 3 for "Identification and partial reconstitution of the biosynthetic pathway of bioactive meroterpenoids from *Hericium erinaceus* (Lion’s Mane mushroom)"

### Orsellinic acid. LC-MS m/z 167 [M-H]$^{-}$; HRMS m/z 169.0497 [M+H]$^{+}$

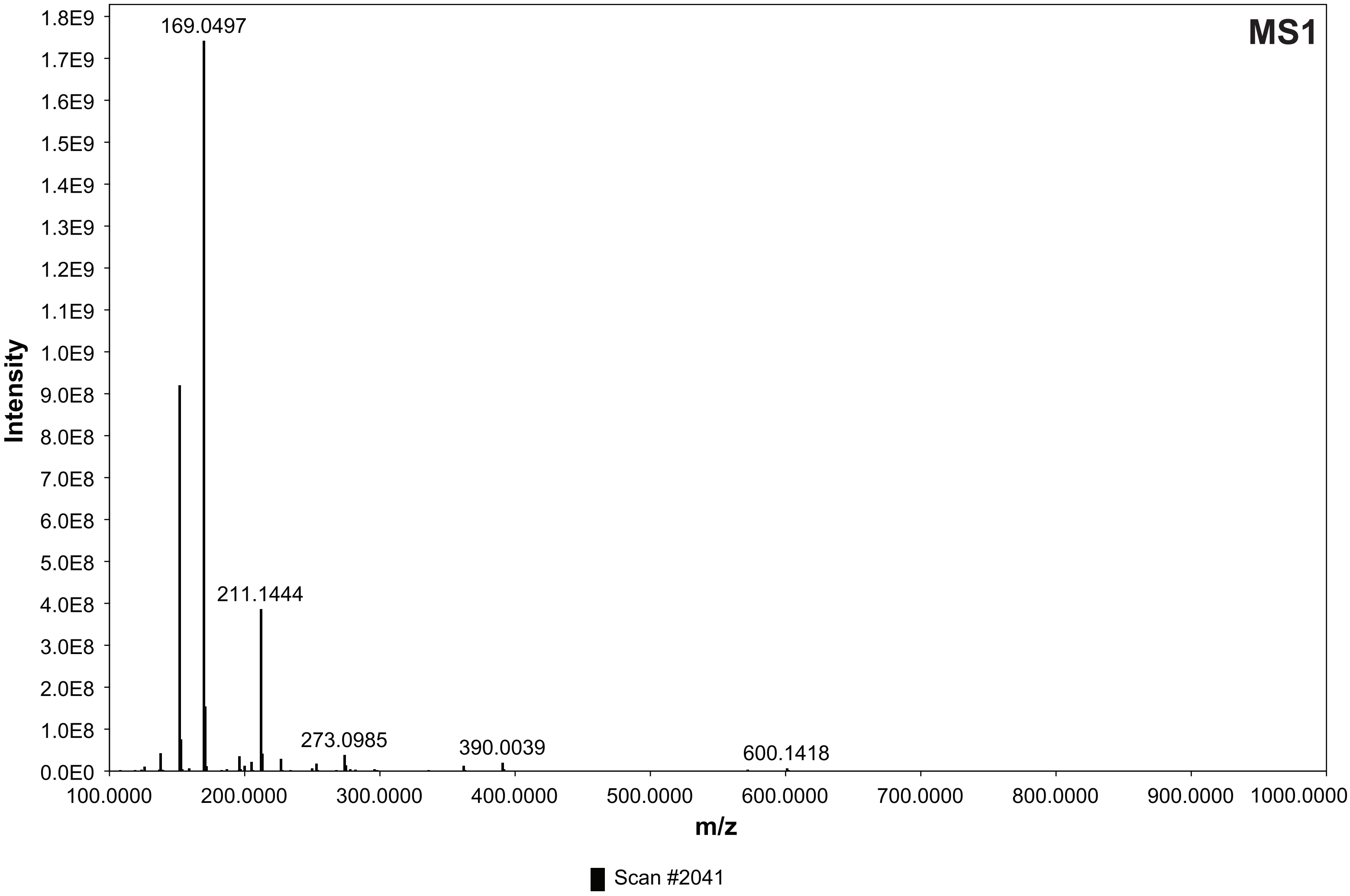


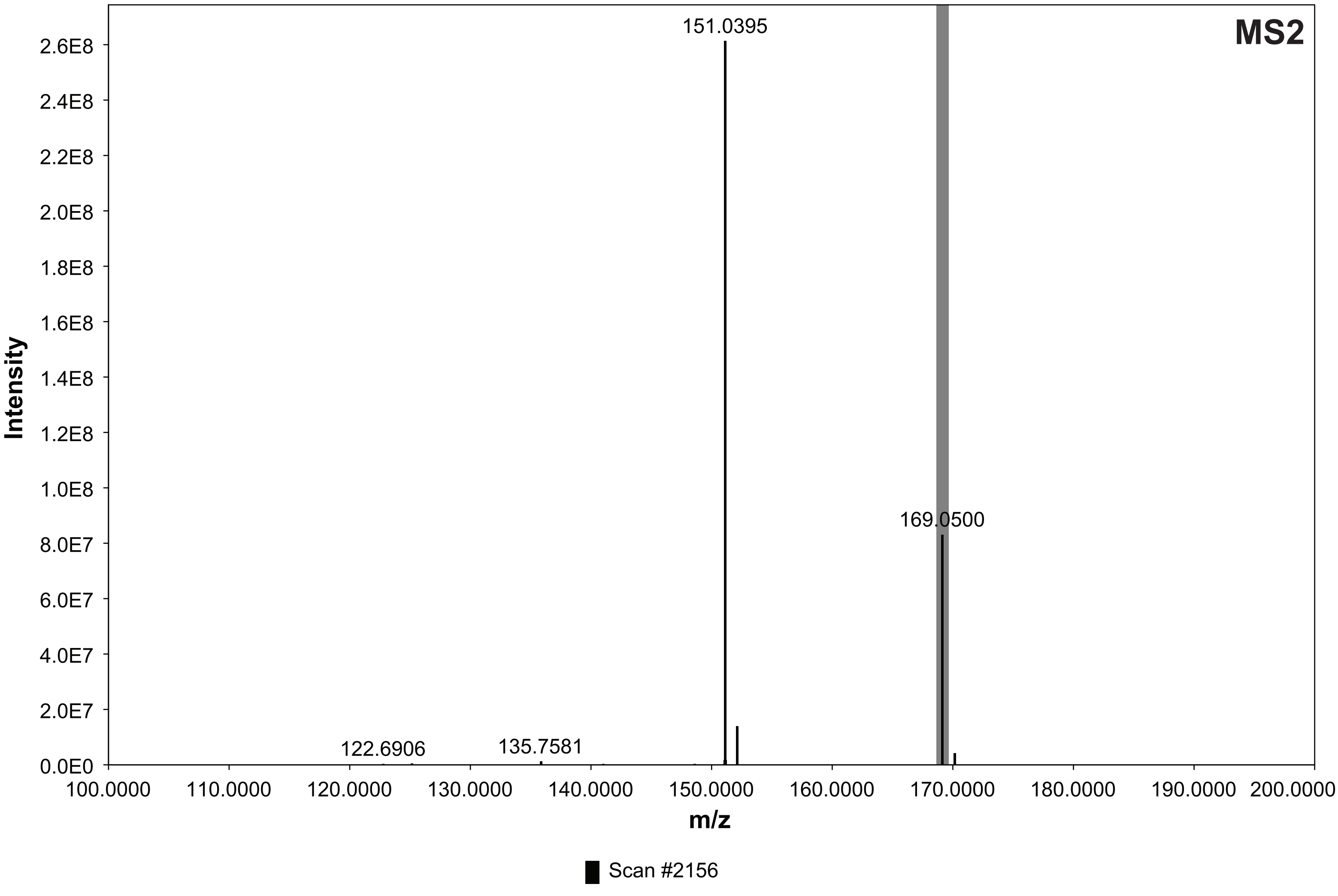

### Orsellinic aldehyde. LC-MS m/z 151 [M-H]$^-$; HRMS m/z 153.0547 [M+H]$^+$

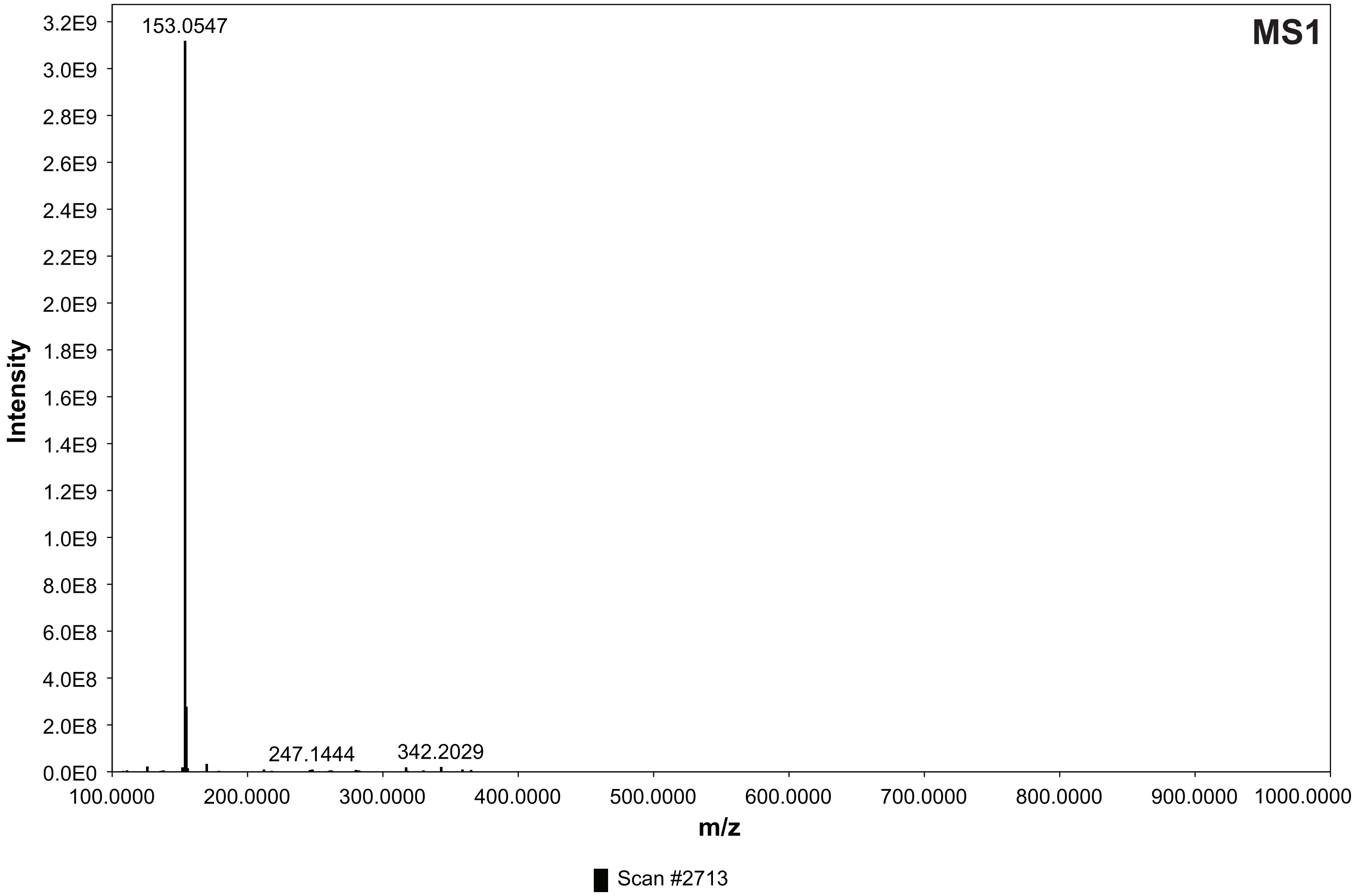


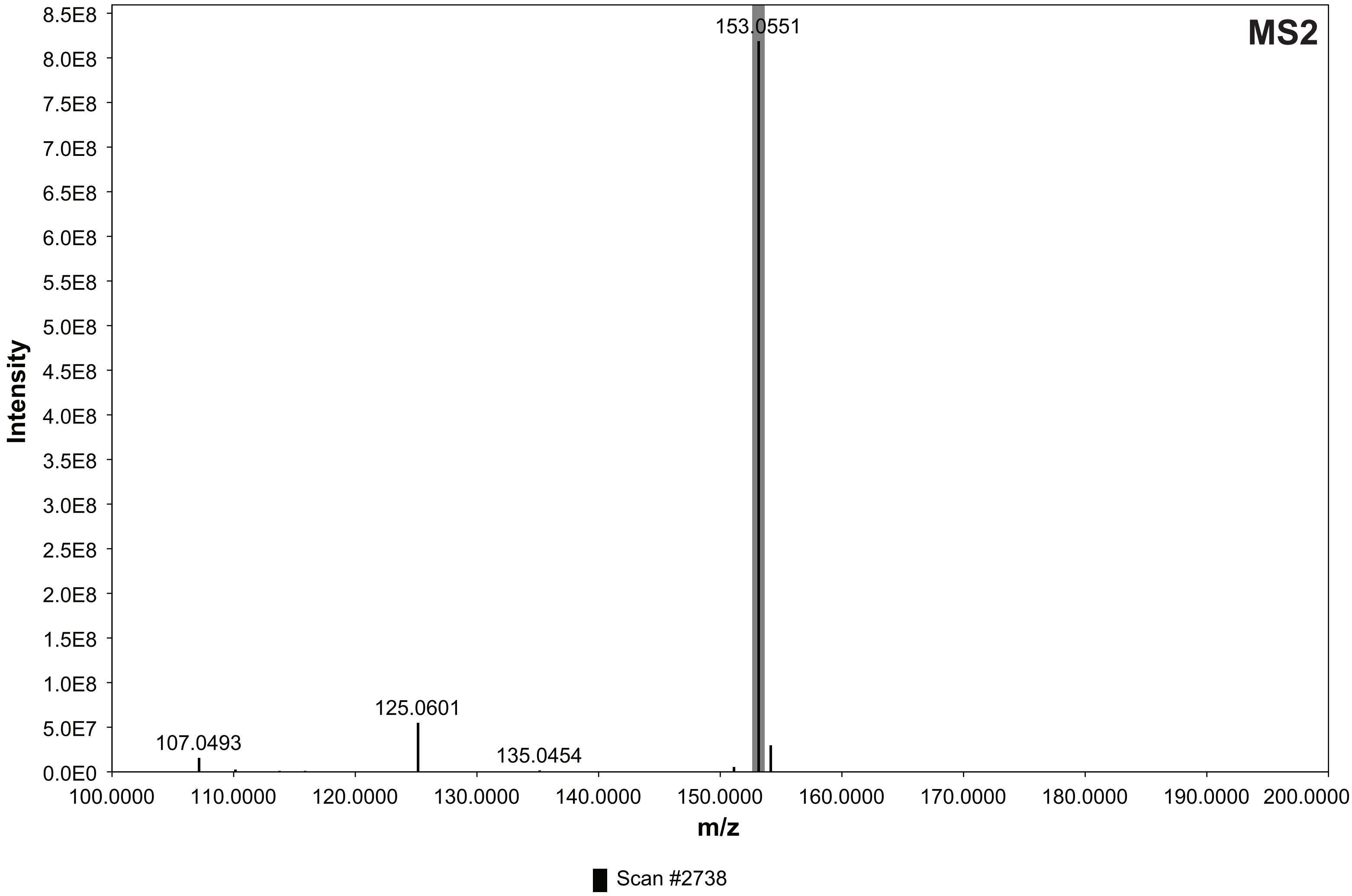

**1. LC-MS m/z 259 $[M-H]^-$; HRMS m/z 261.1124 $[M+H]^+$**

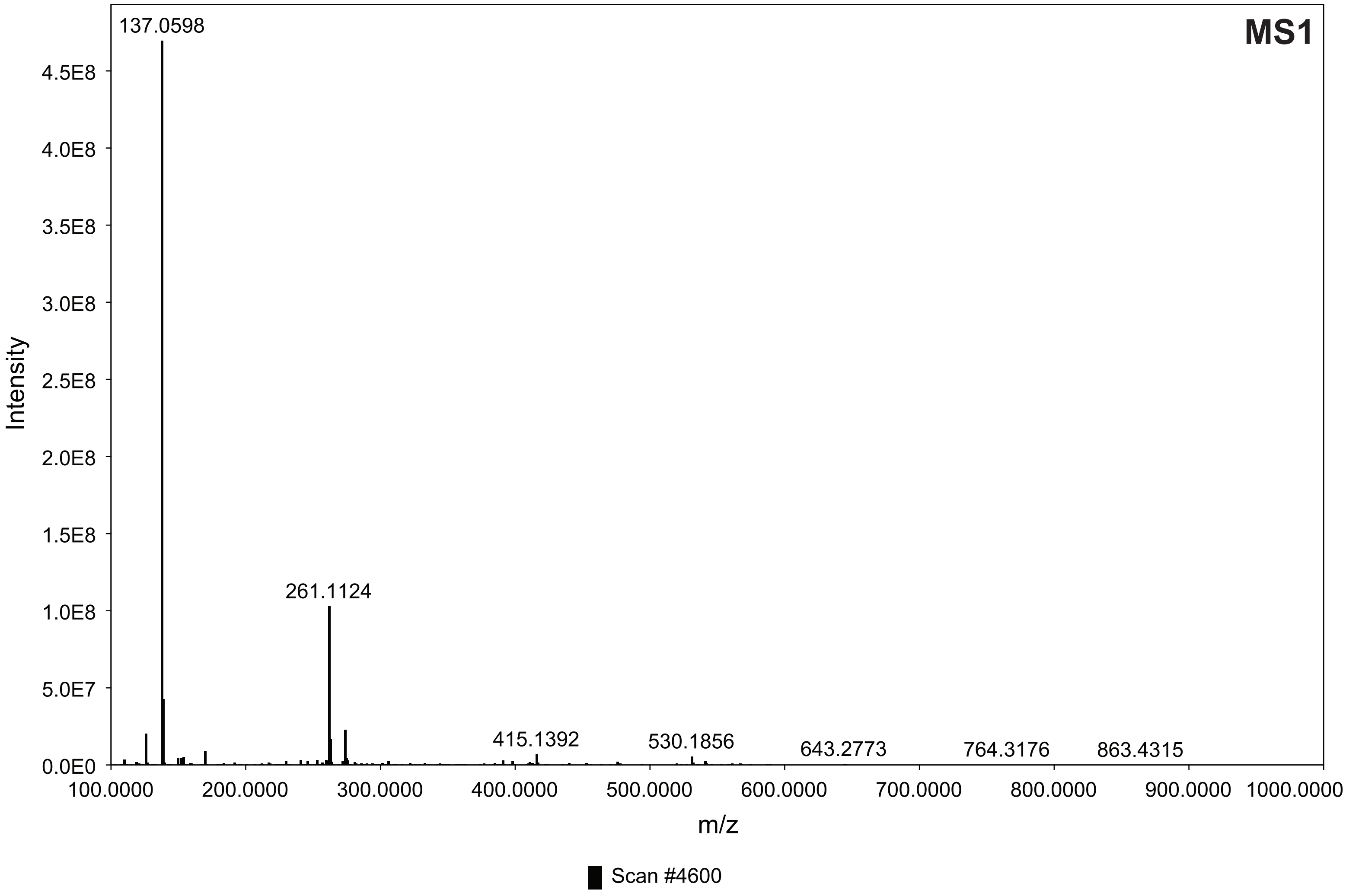


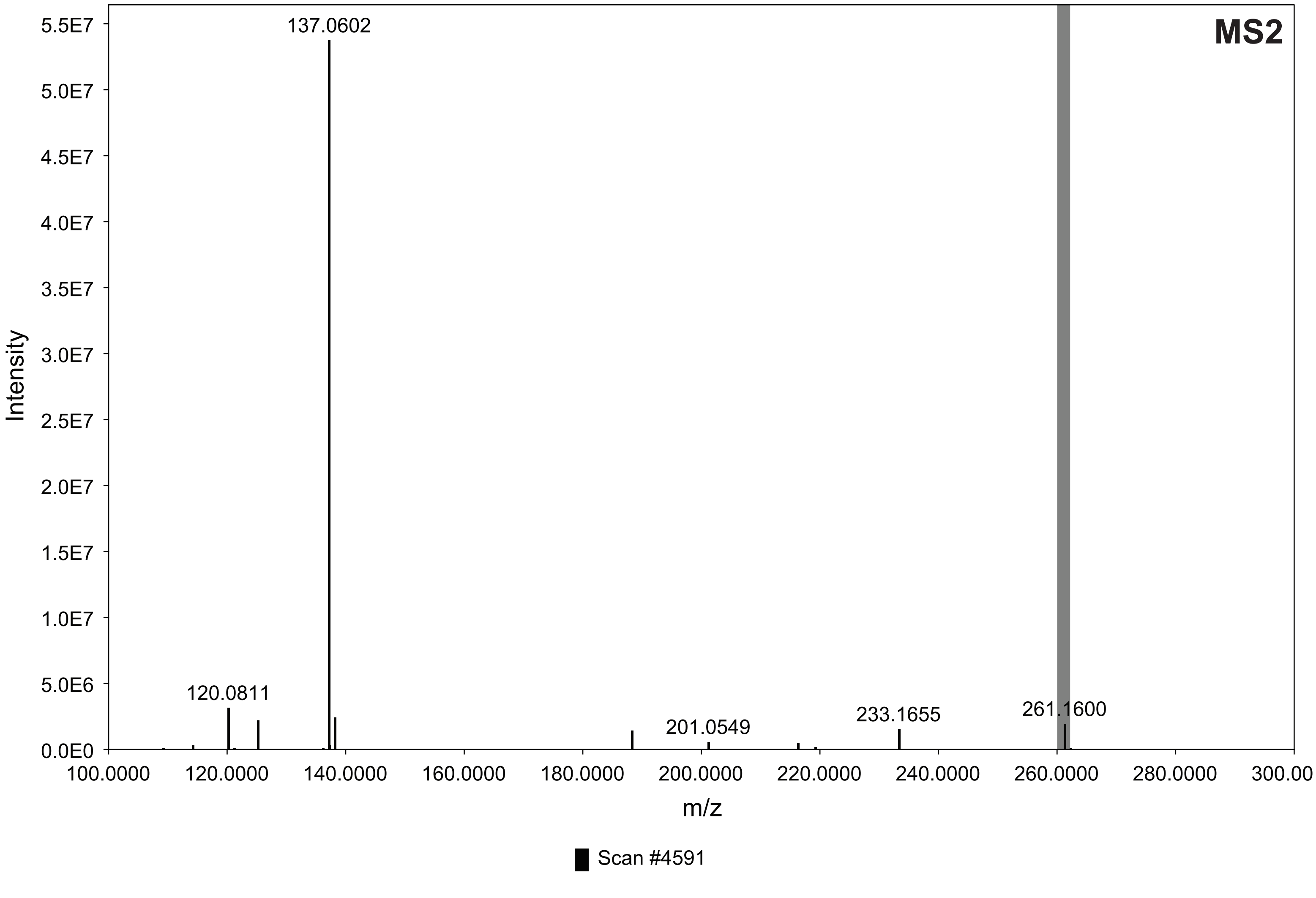

#### 2. LC-MS m/z 261 [M-H]$^-$; HRMS m/z 263.0915 [M+H]$^+$

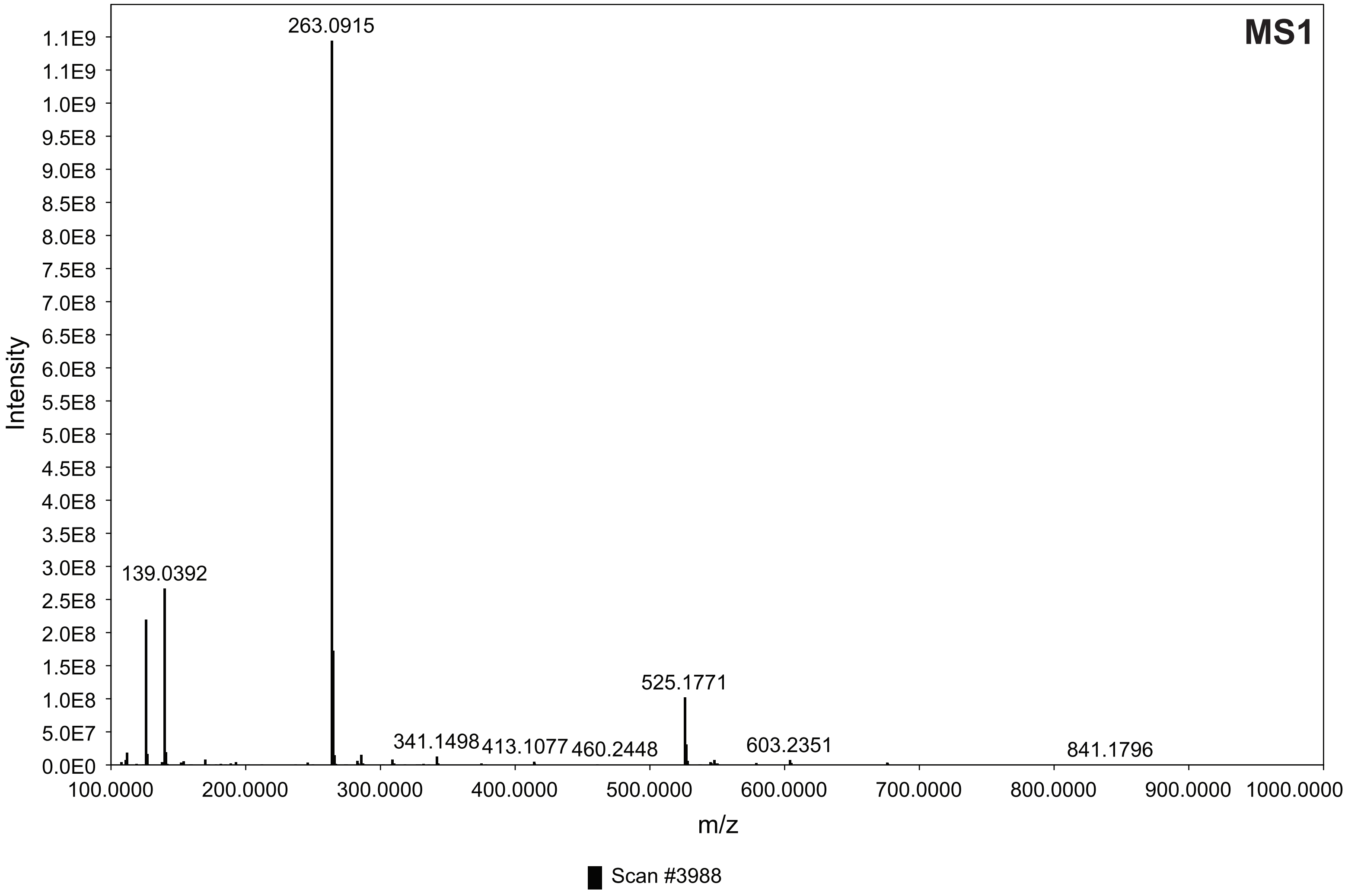


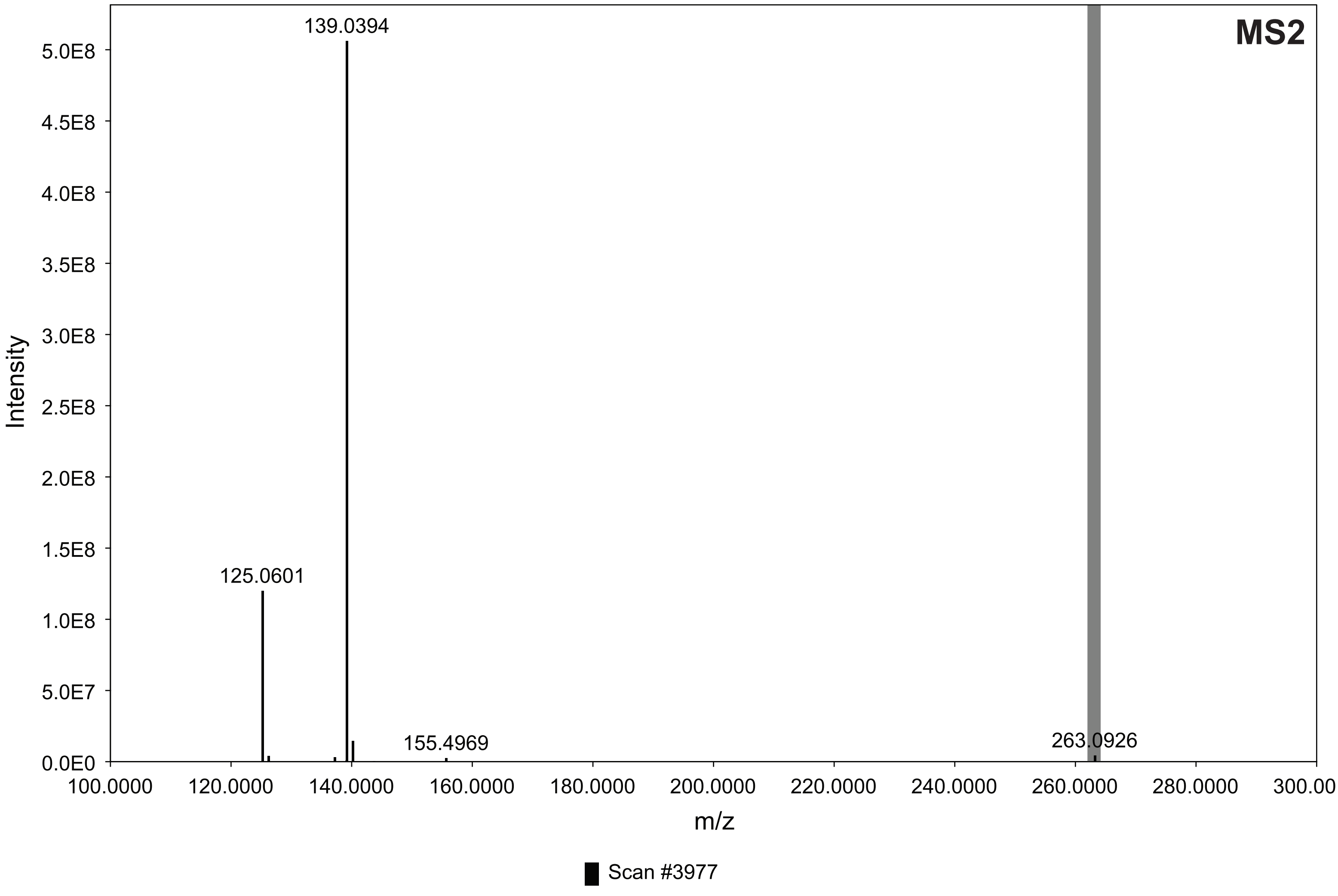

#### 3. LC-MS m/z 303 [M-H]⁻; HRMS m/z 305.1021 [M+H]⁺

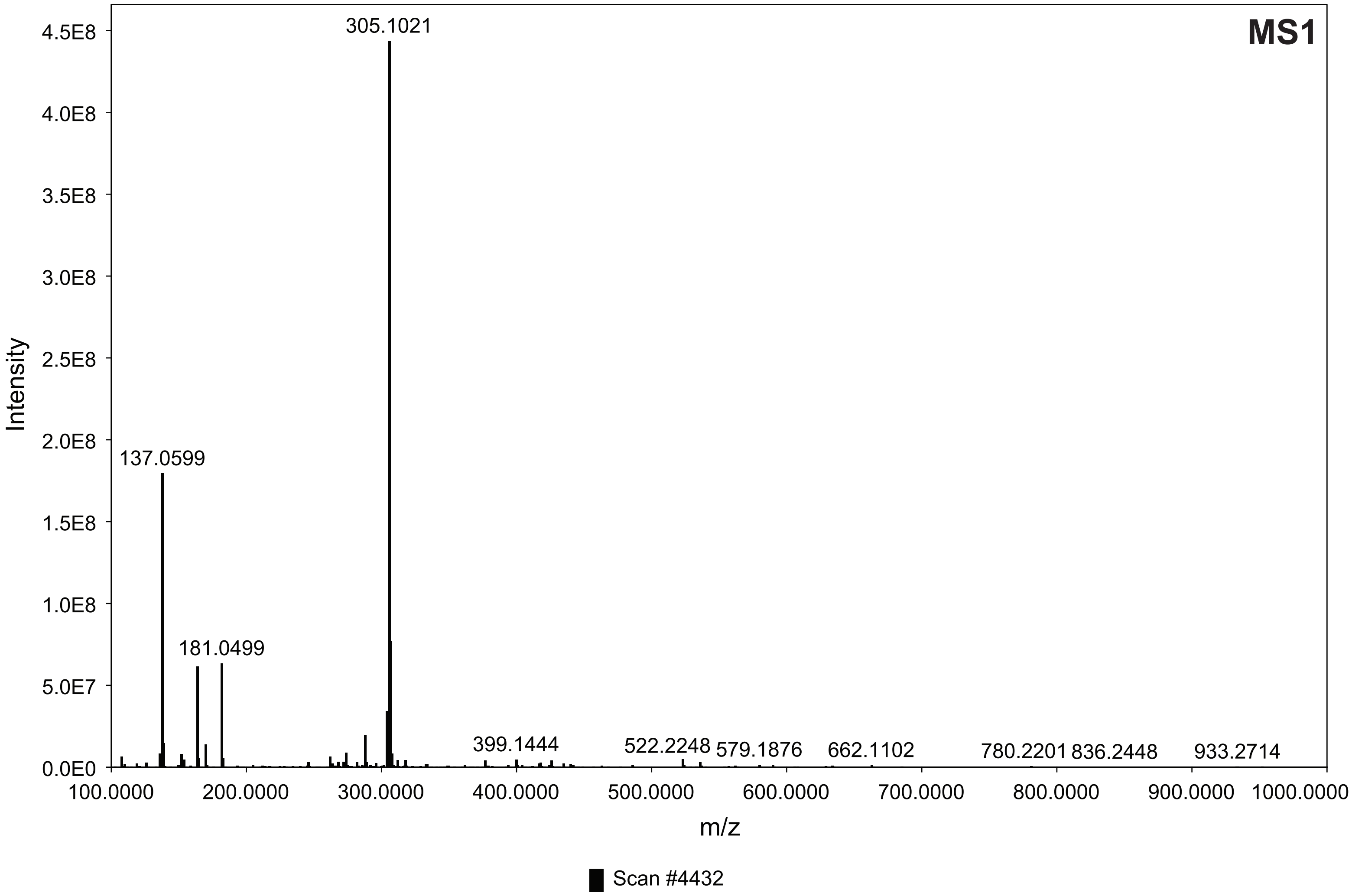


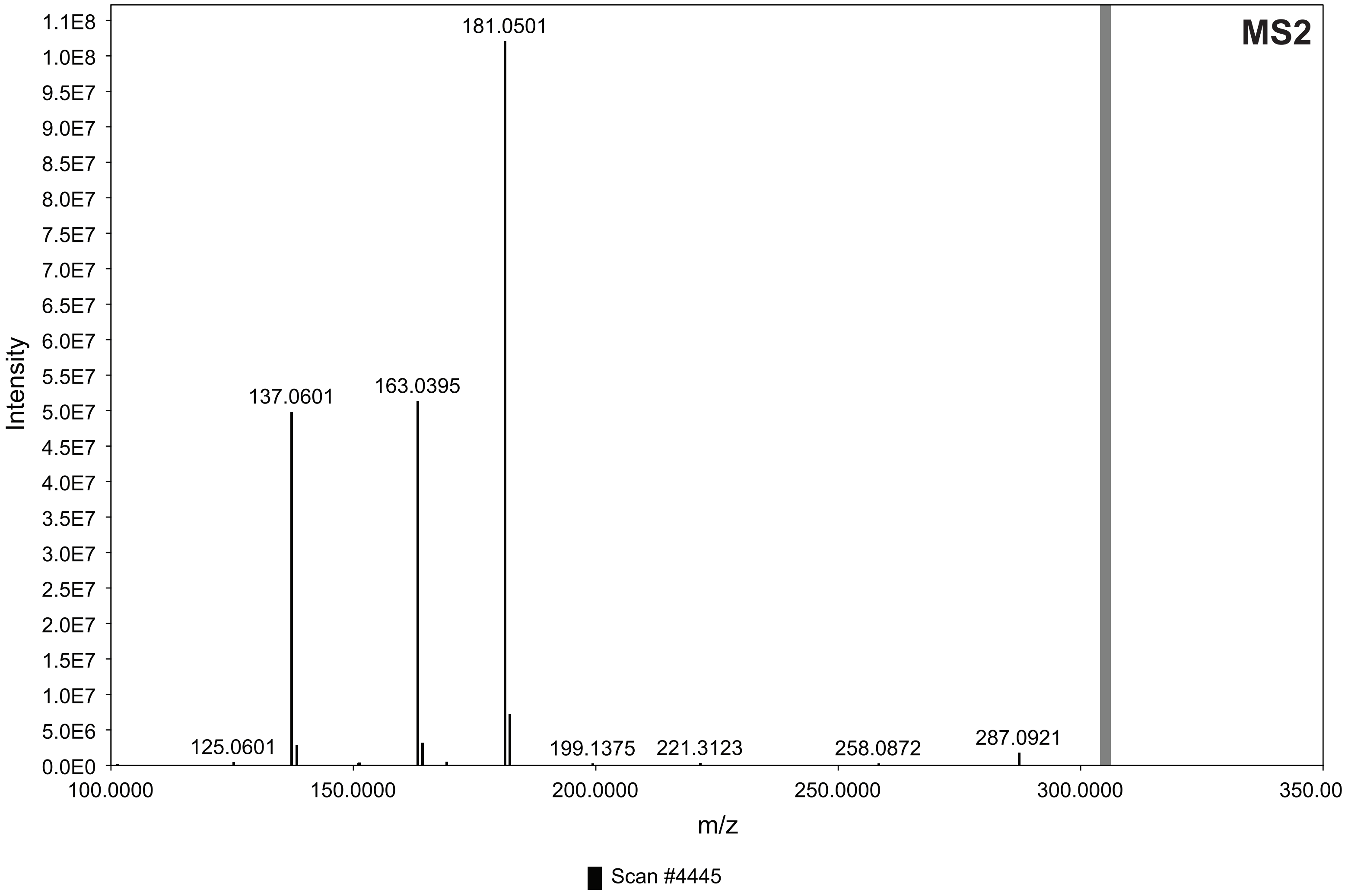

#### 4. LC-MS m/z 395 [M-H]$^-$; HRMS m/z 397.1648 [M+H]$^+$

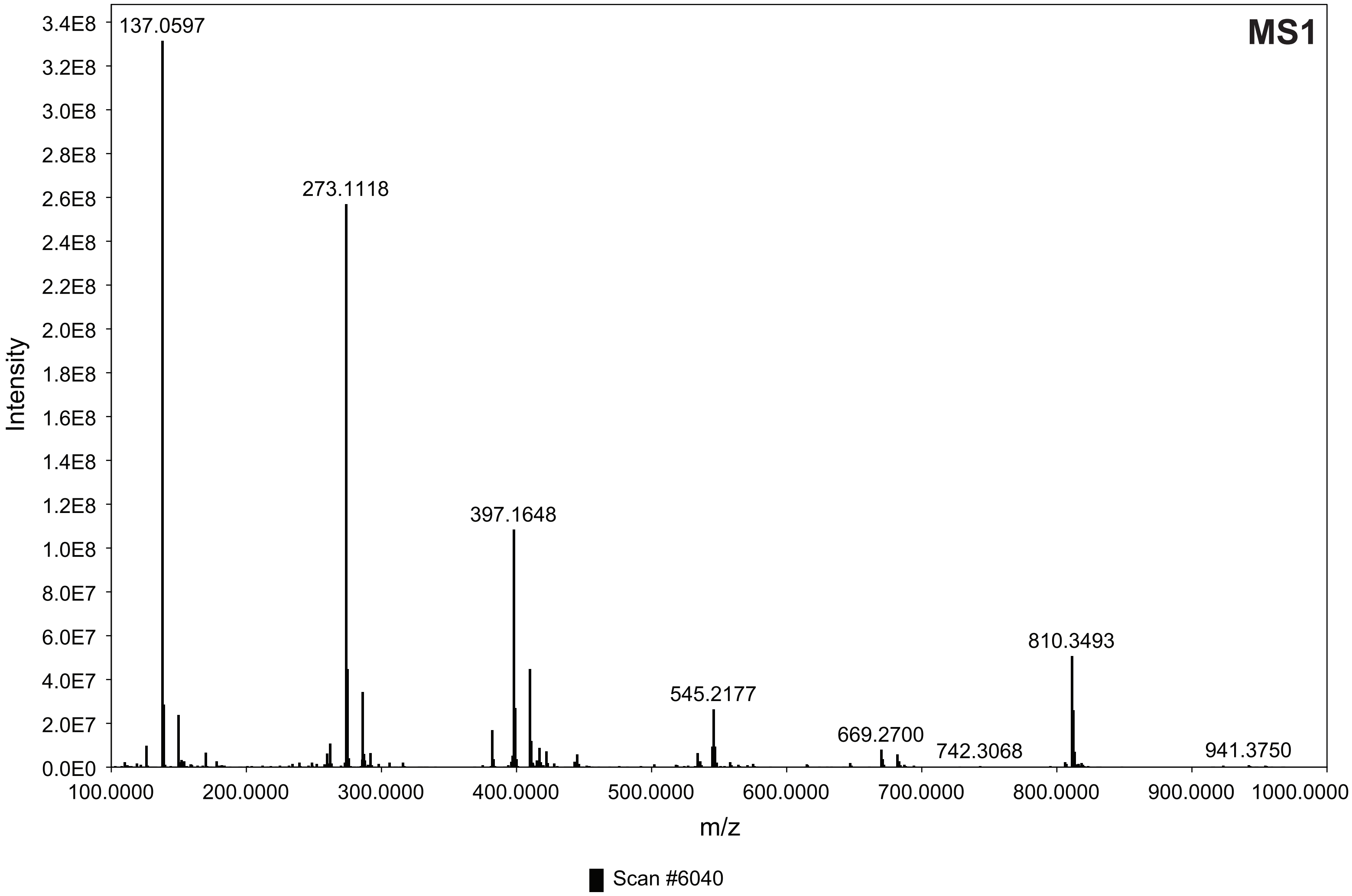


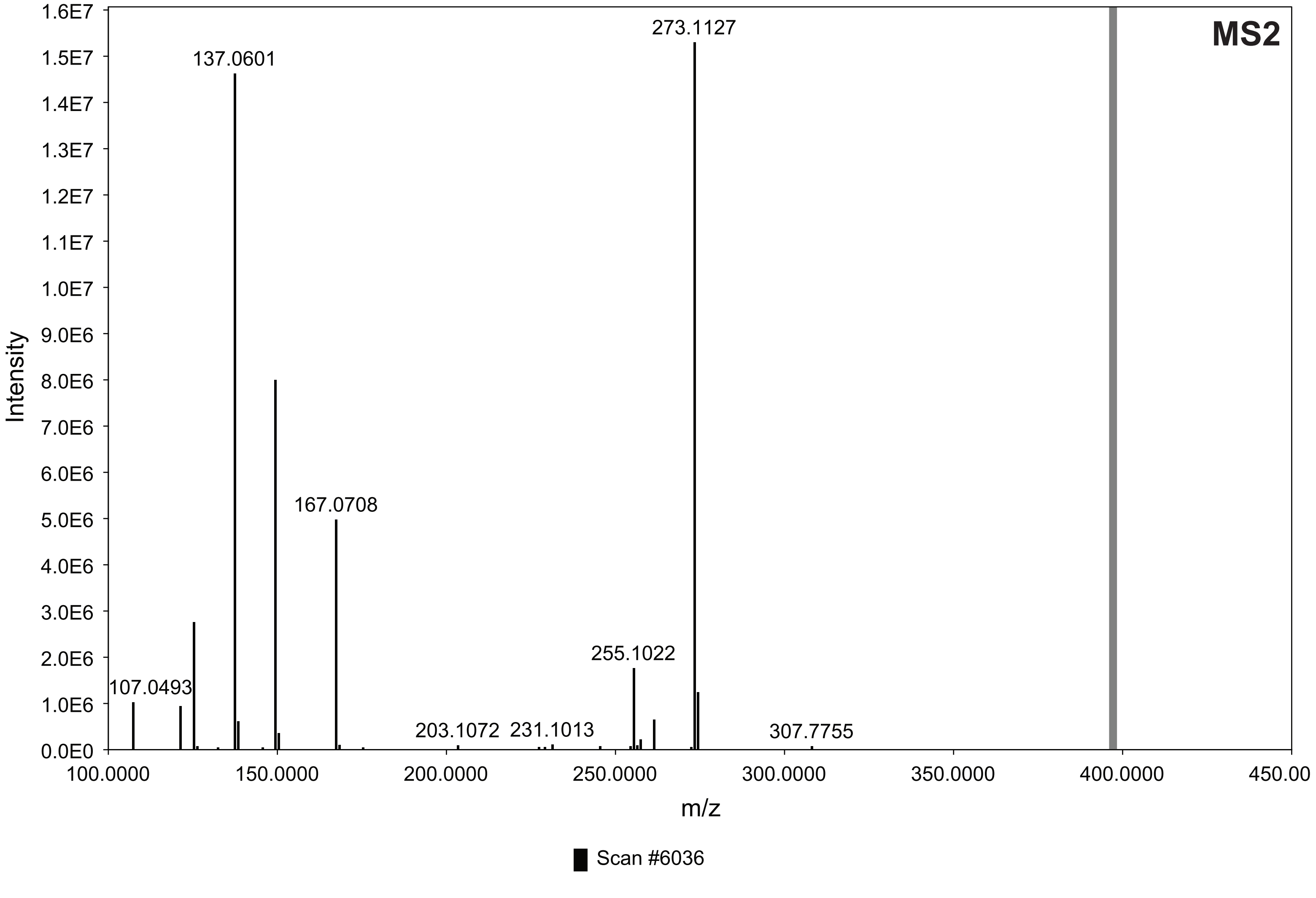

**5. LC-MS m/z 423 [M-H]⁻; HRMS m/z 425.1599 [M+H]⁺**

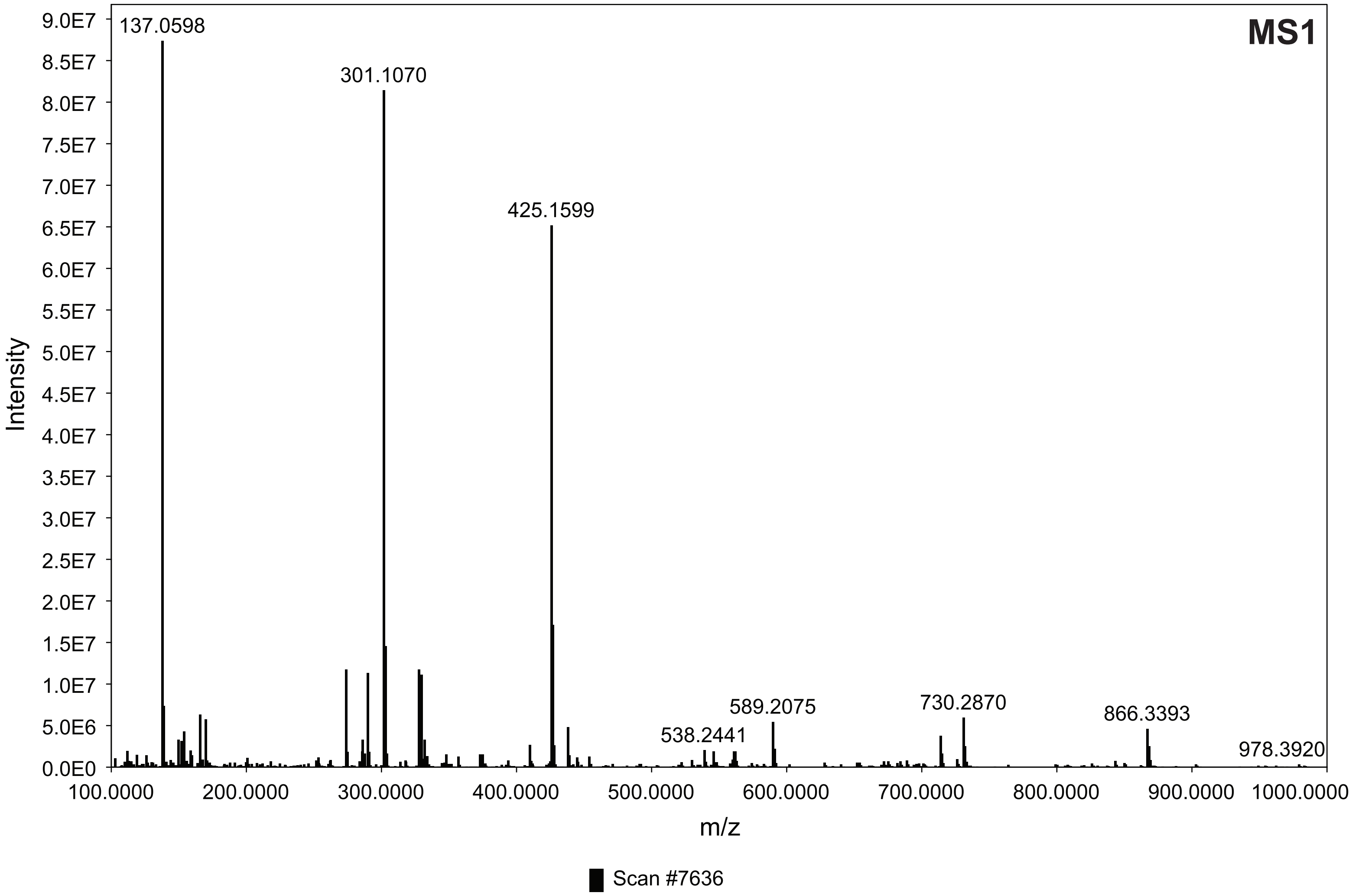


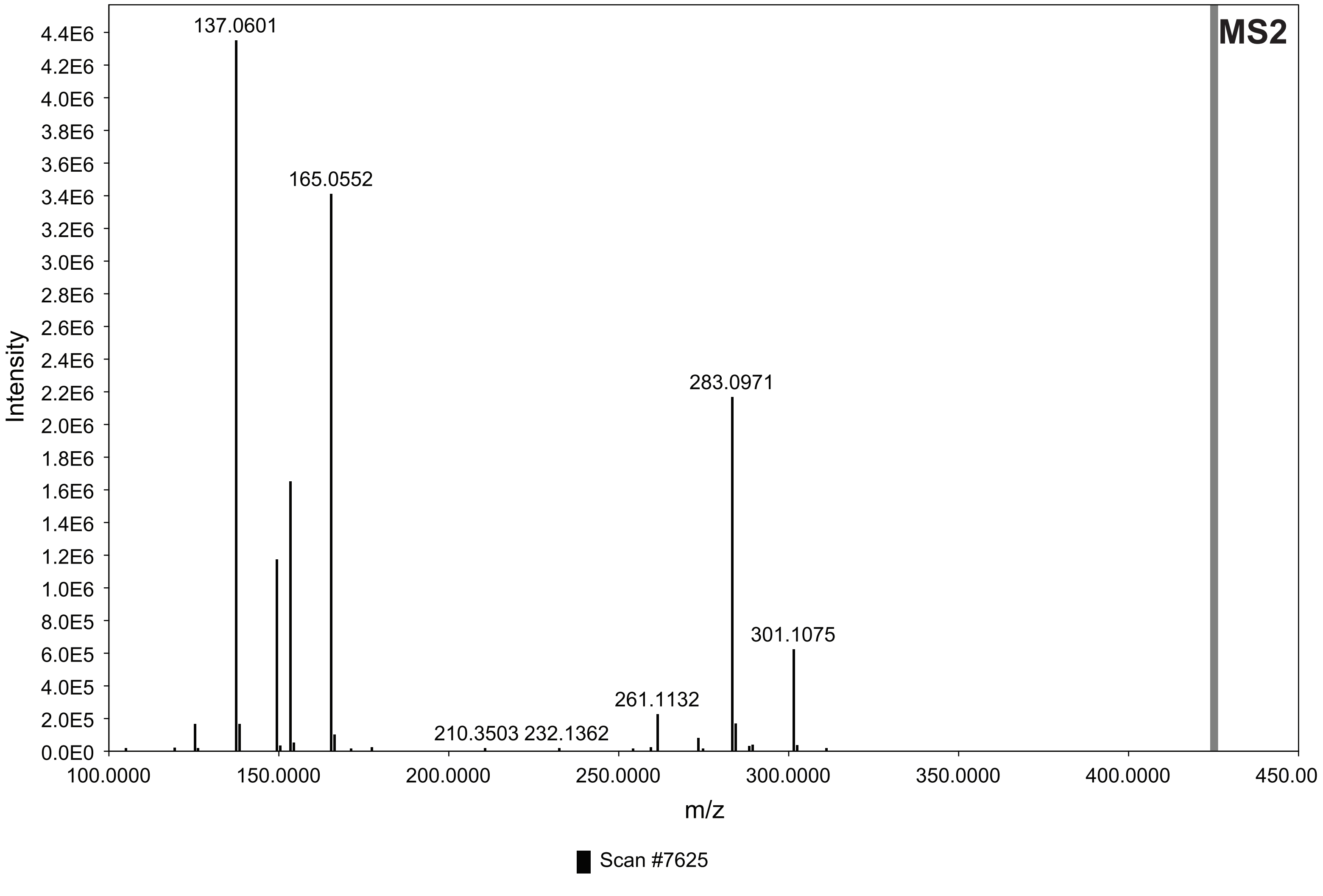

#### 6. LC-MS m/z 439 $[M-H]^-$; HRMS m/z 441.1549 $[M+H]^+$

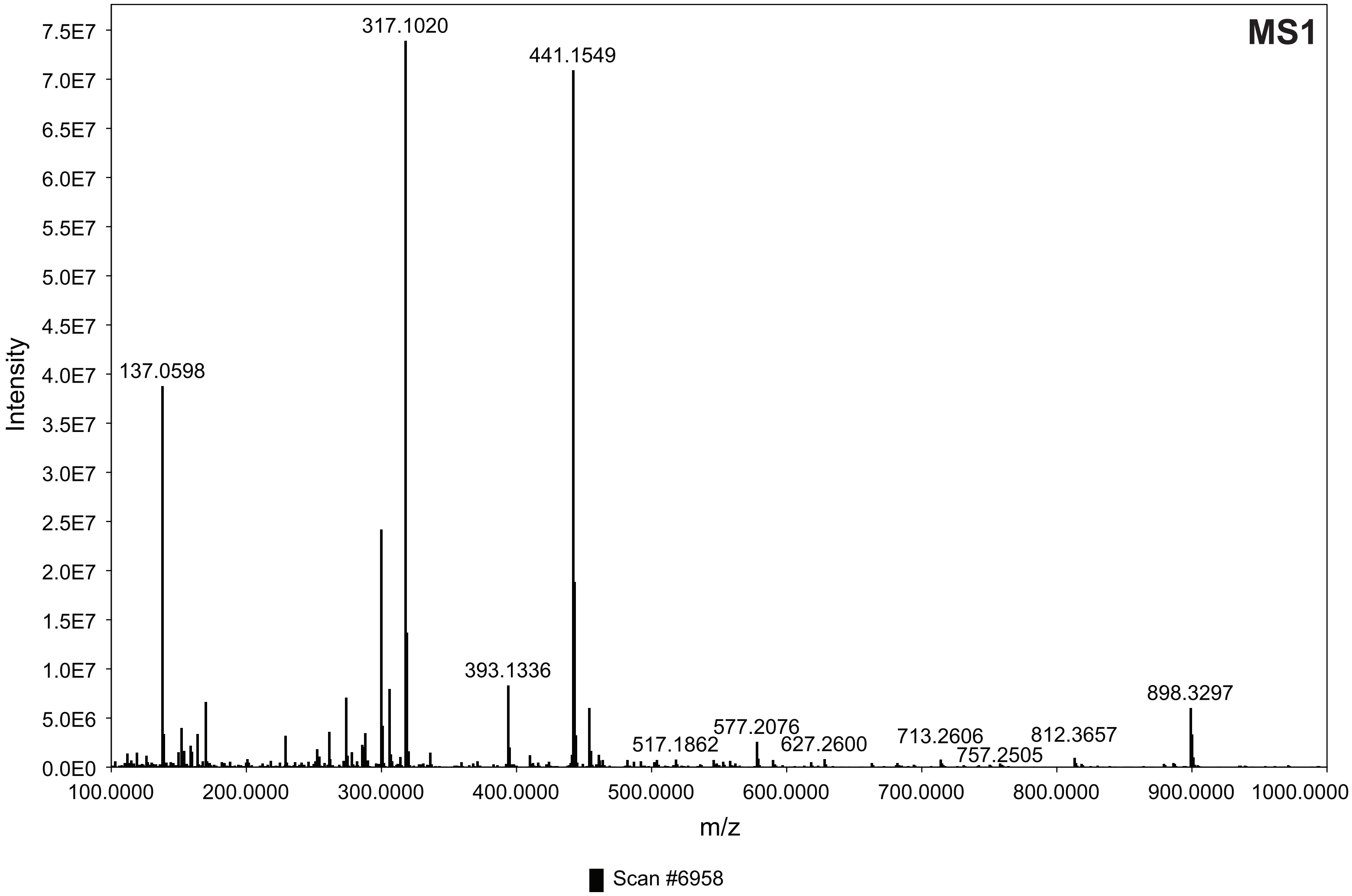


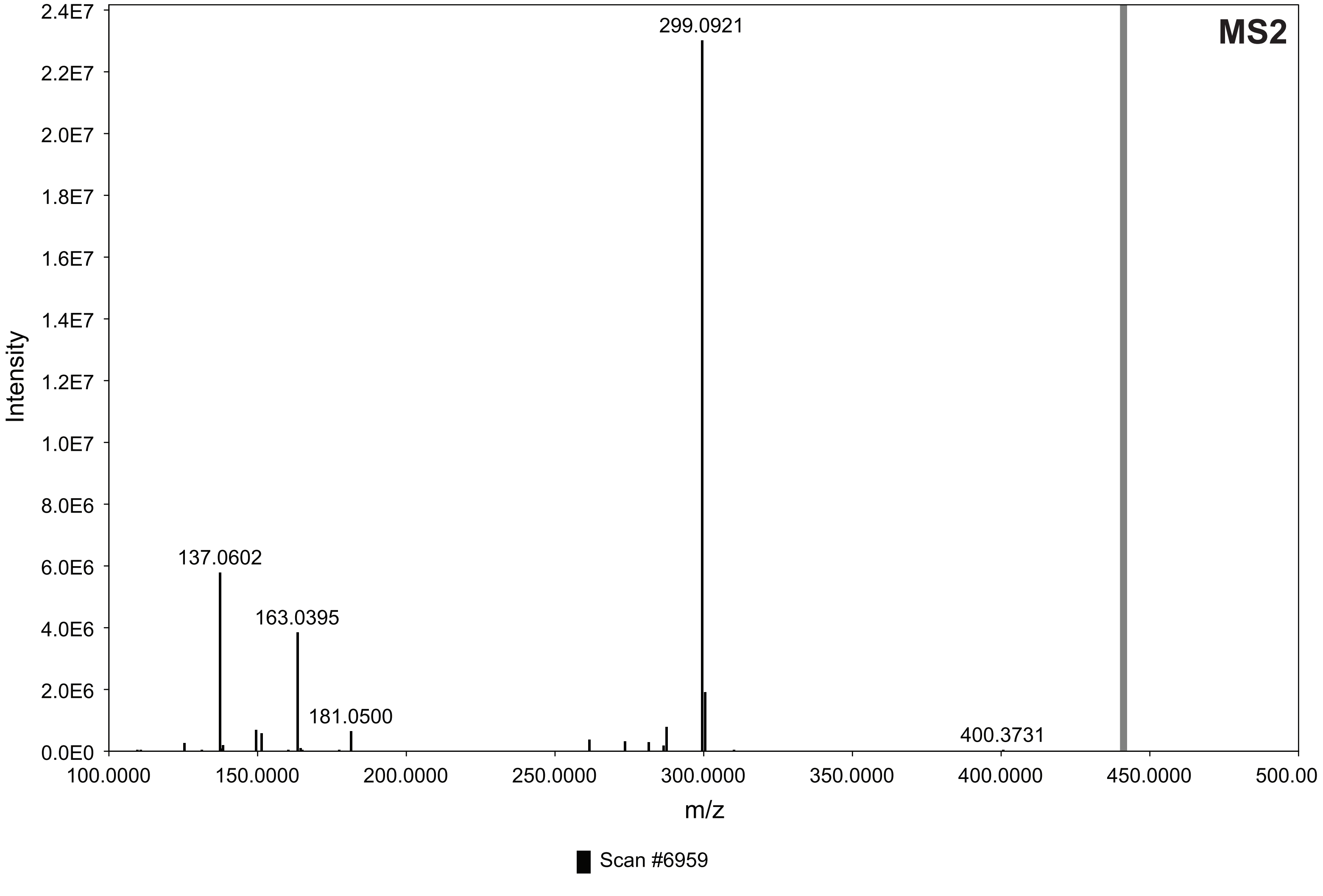

**7. LC-MS m/z 531 [M-H]⁻; HRMS m/z 533.2176 [M+H]⁺**

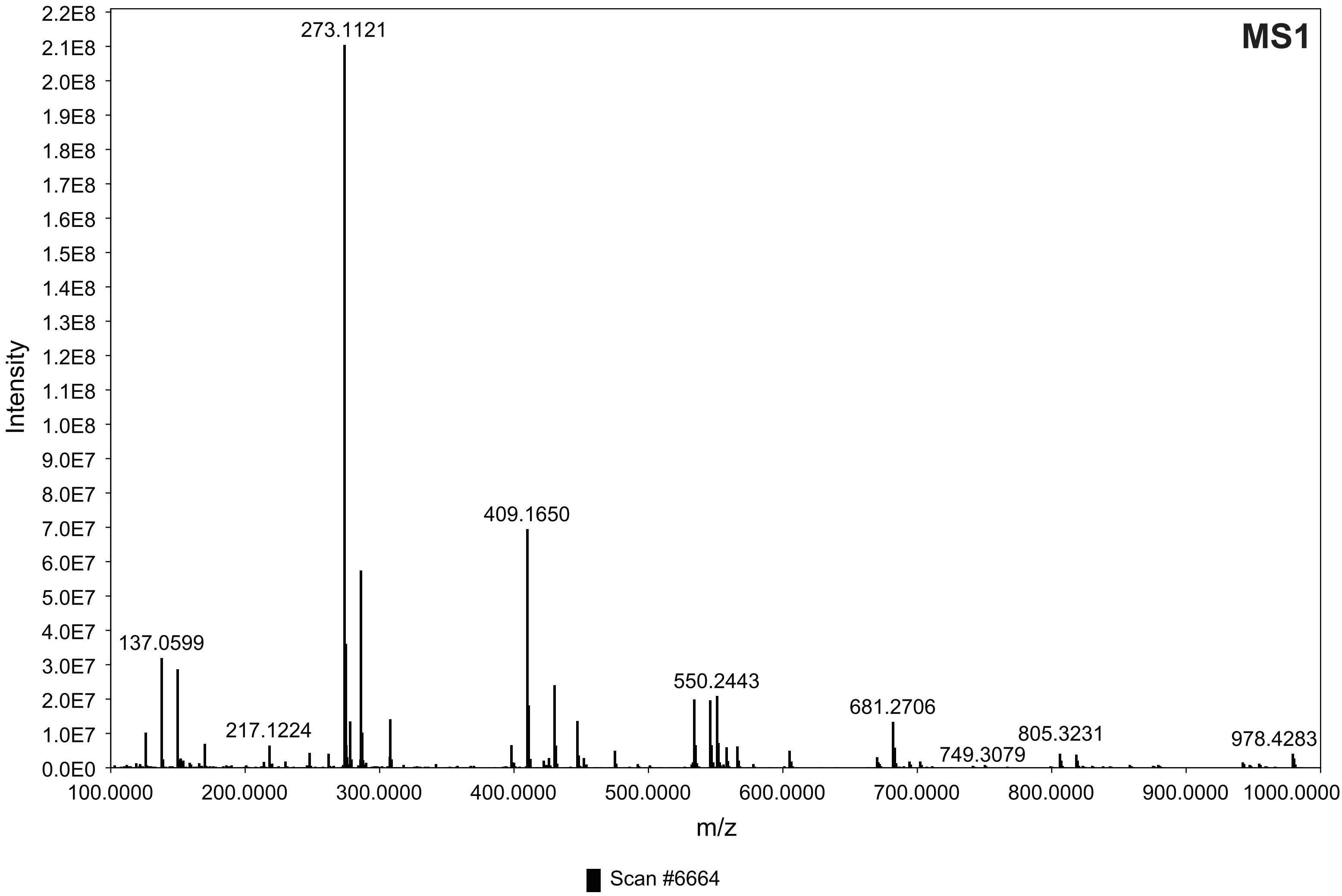


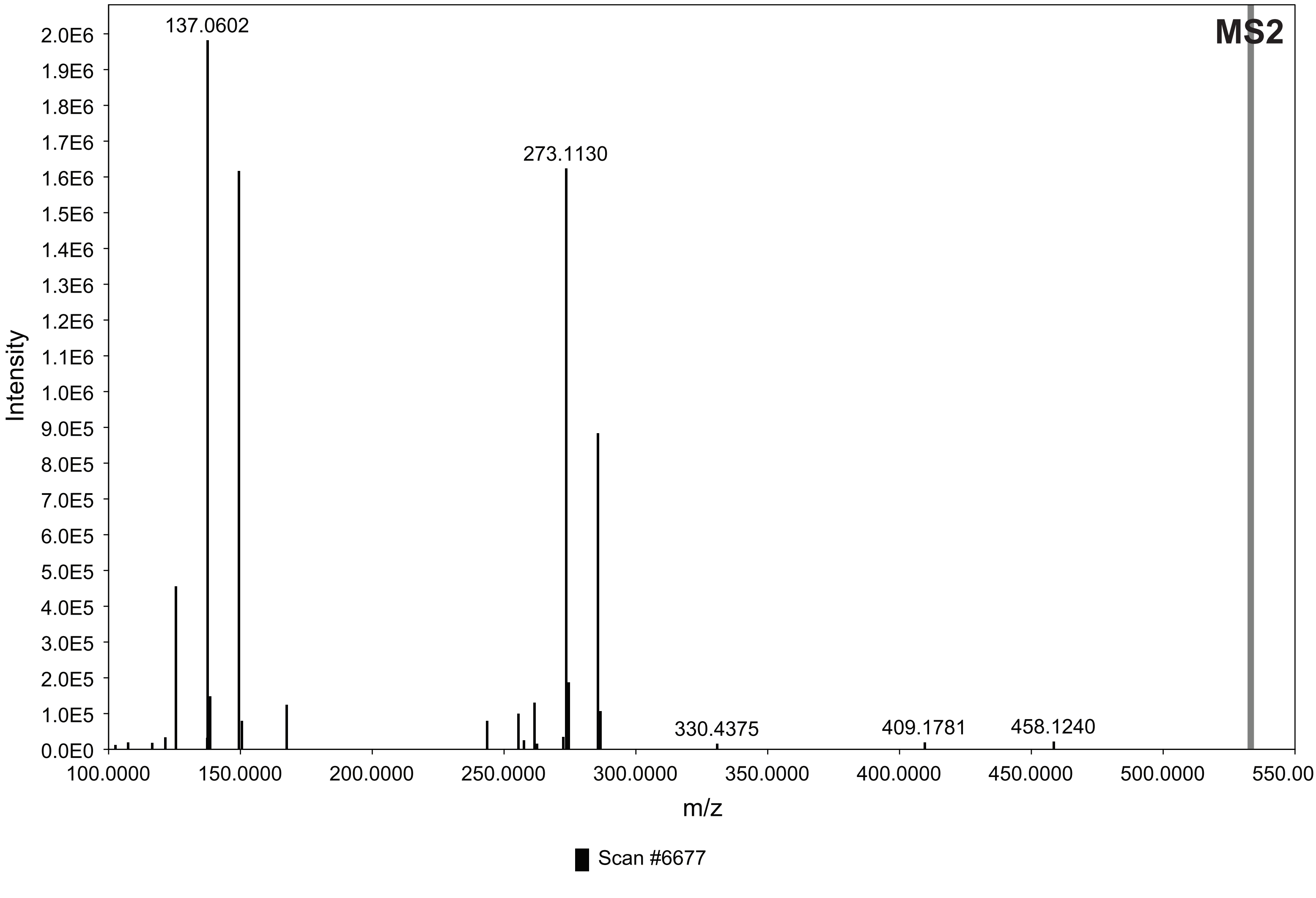

### 8. LC-MS m/z 575 [M-H]⁻; HRMS m/z 577.2078 [M+H]⁺

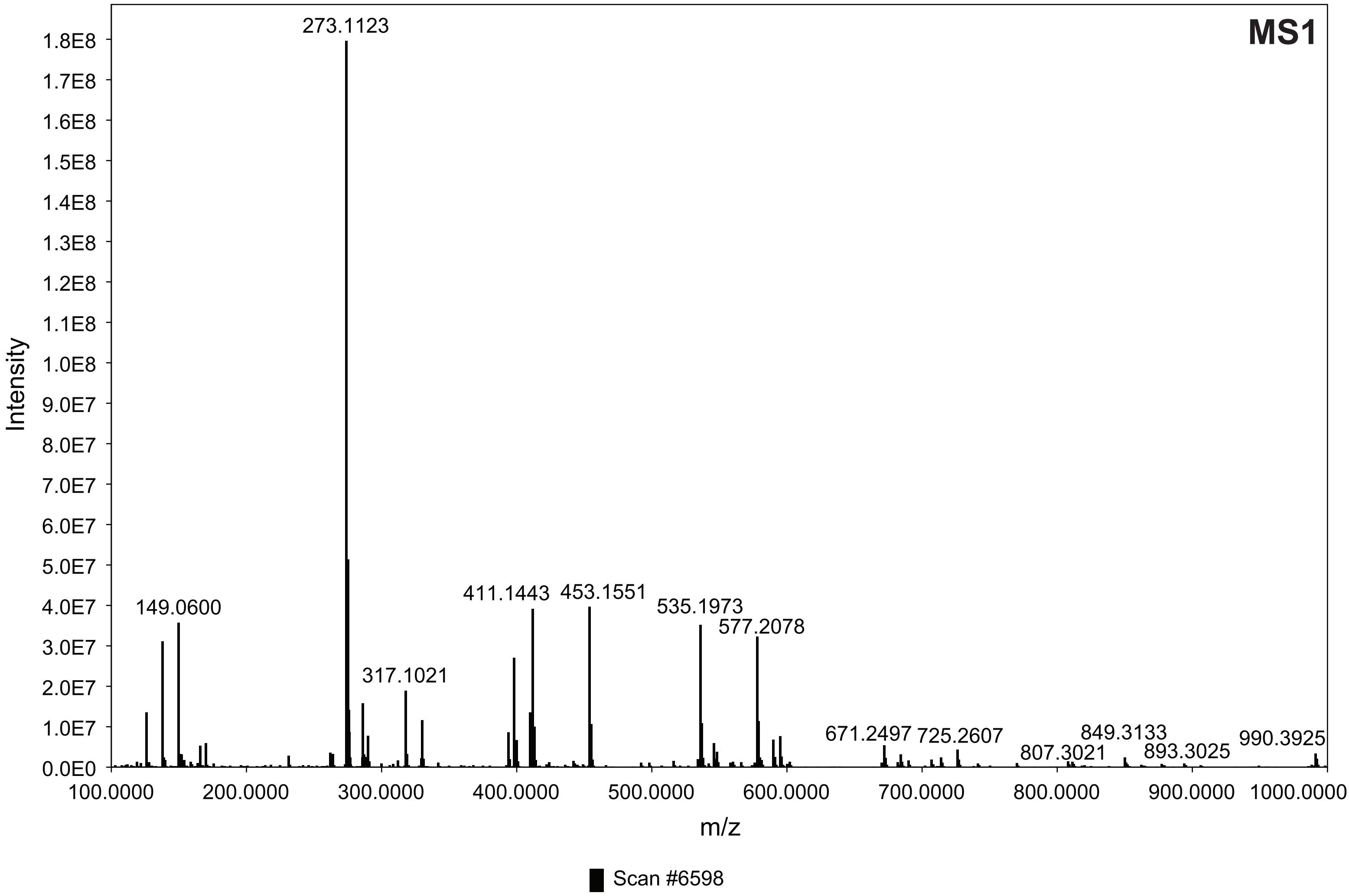


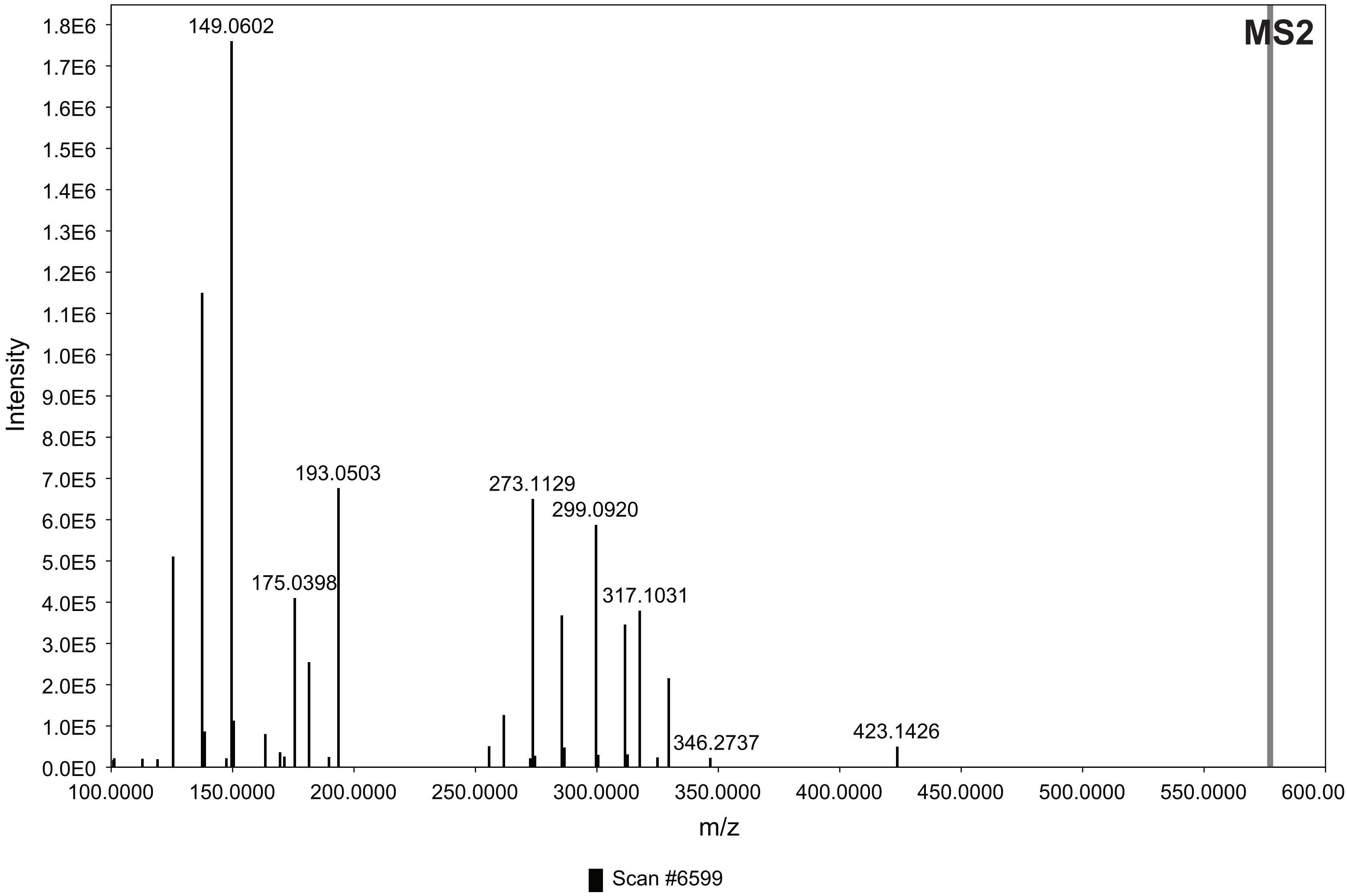

**9. LC-MS m/z 667 [M-H]$^{-}$; HRMS m/z 669.2701 [M+H]$^{+}$**

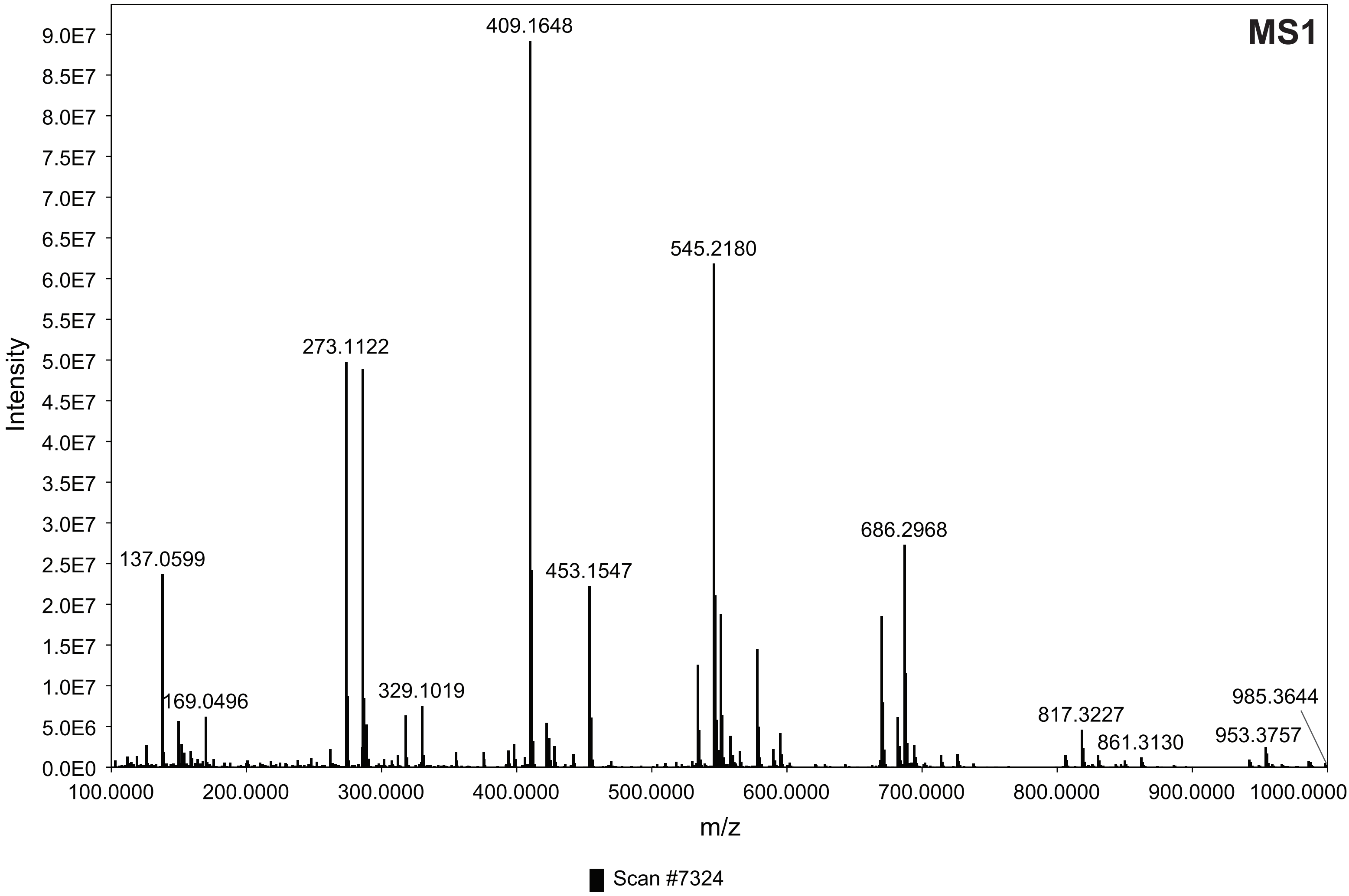


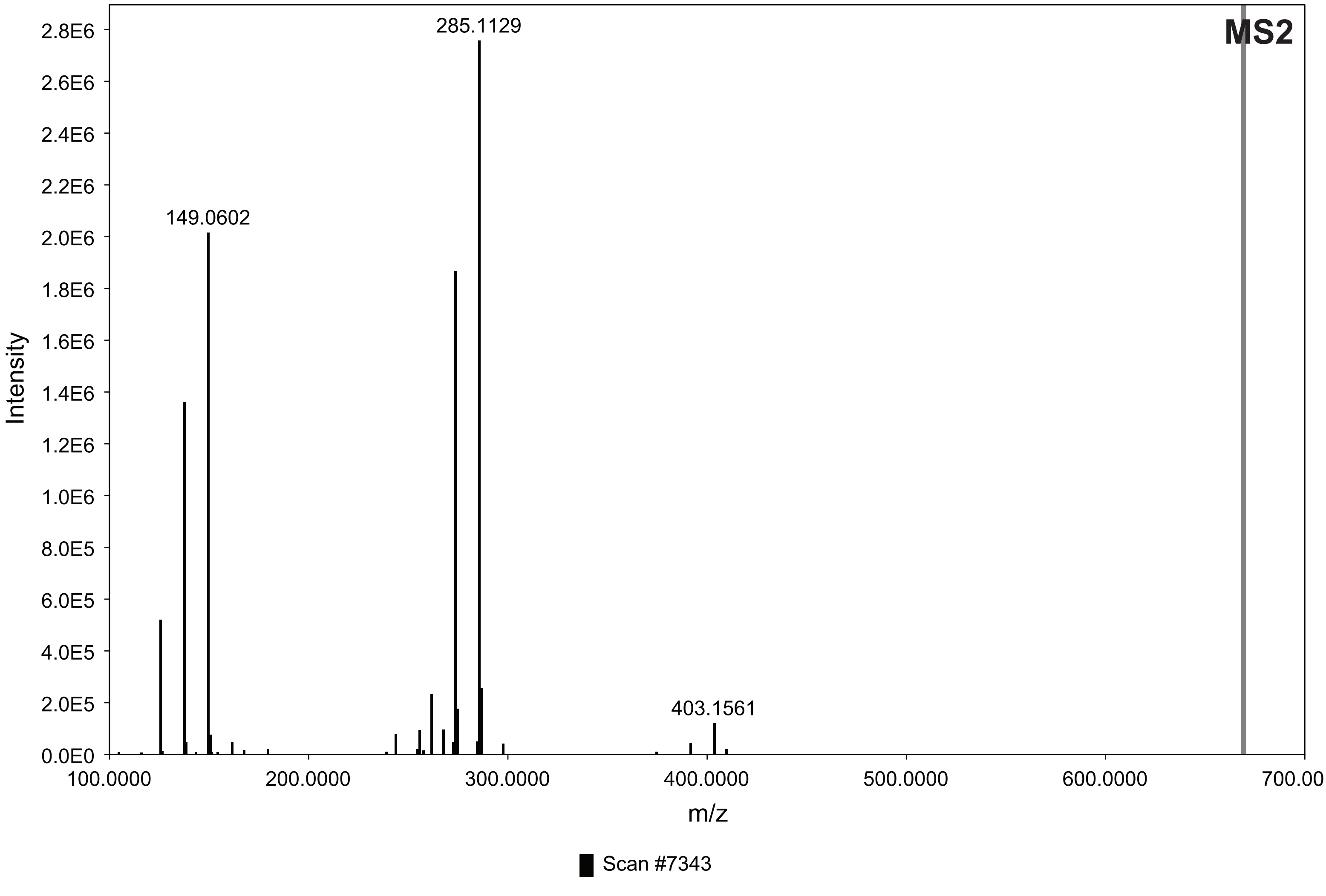

**10. LC-MS m/z 711 [M-H]⁻; HRMS m/z 713.2601 [M+H]⁺**

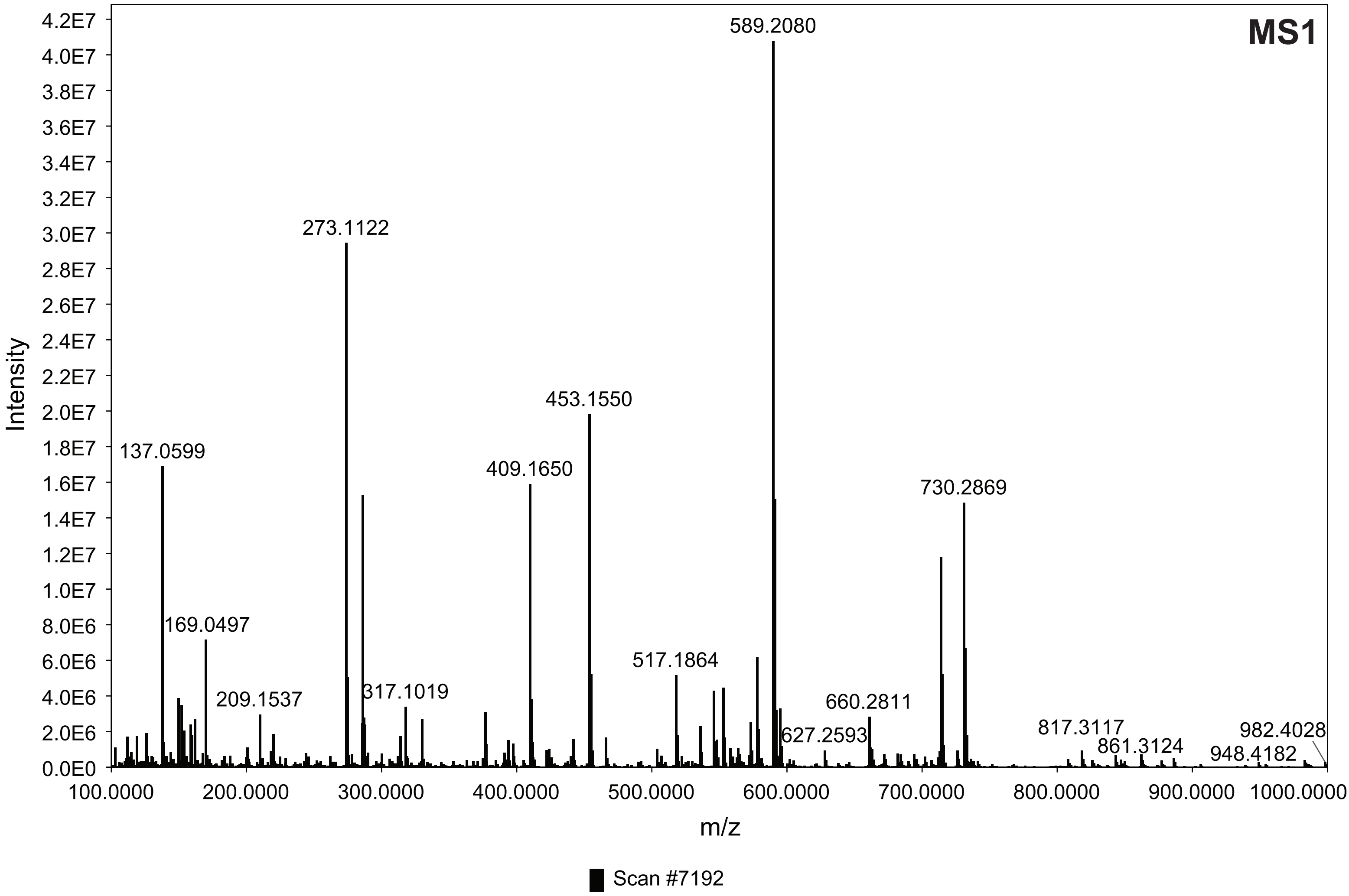


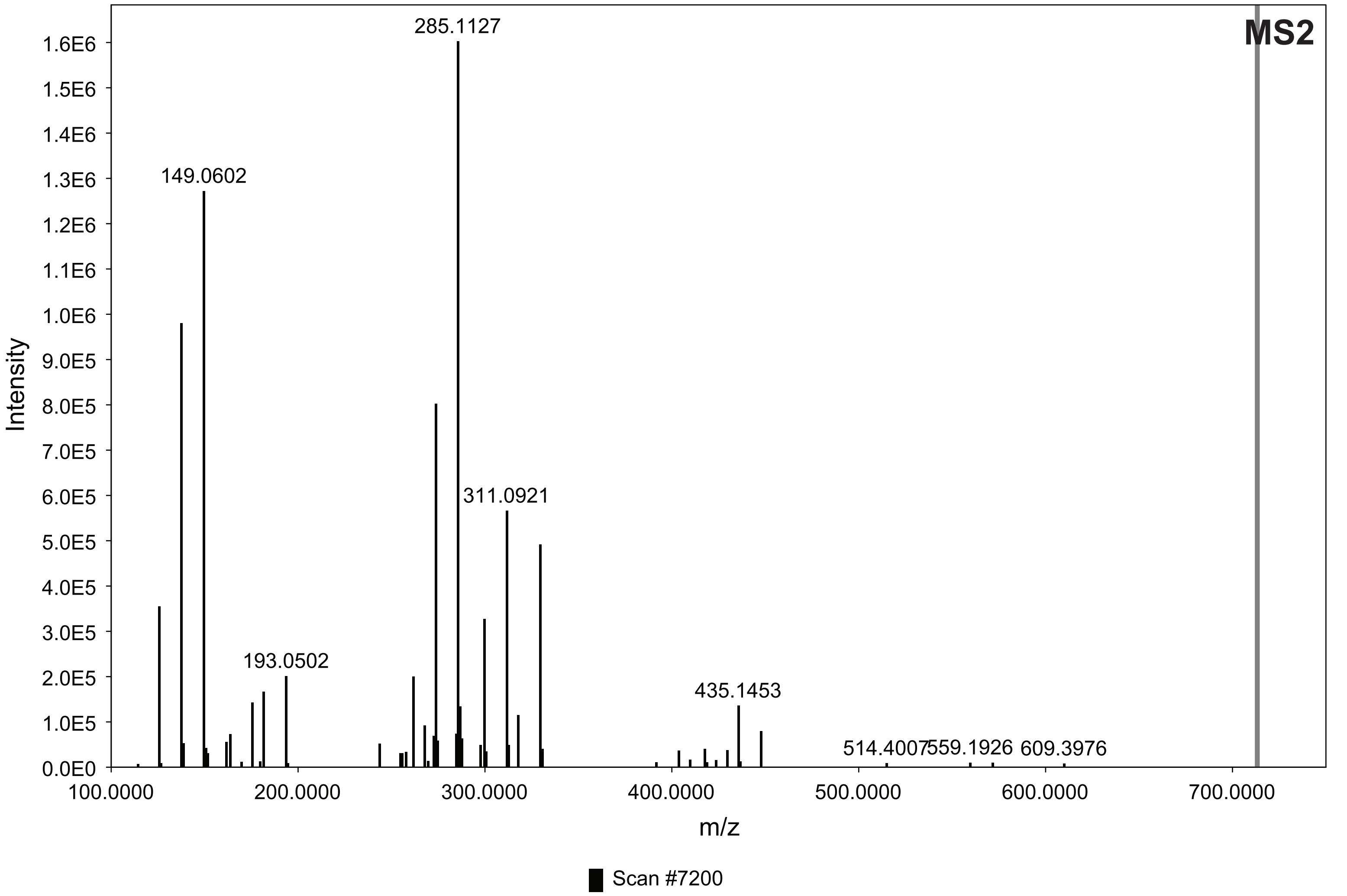

#### 11. LC-MS m/z 803 $[M-H]^-$; HRMS m/z 805.3232 $[M+H]^+$

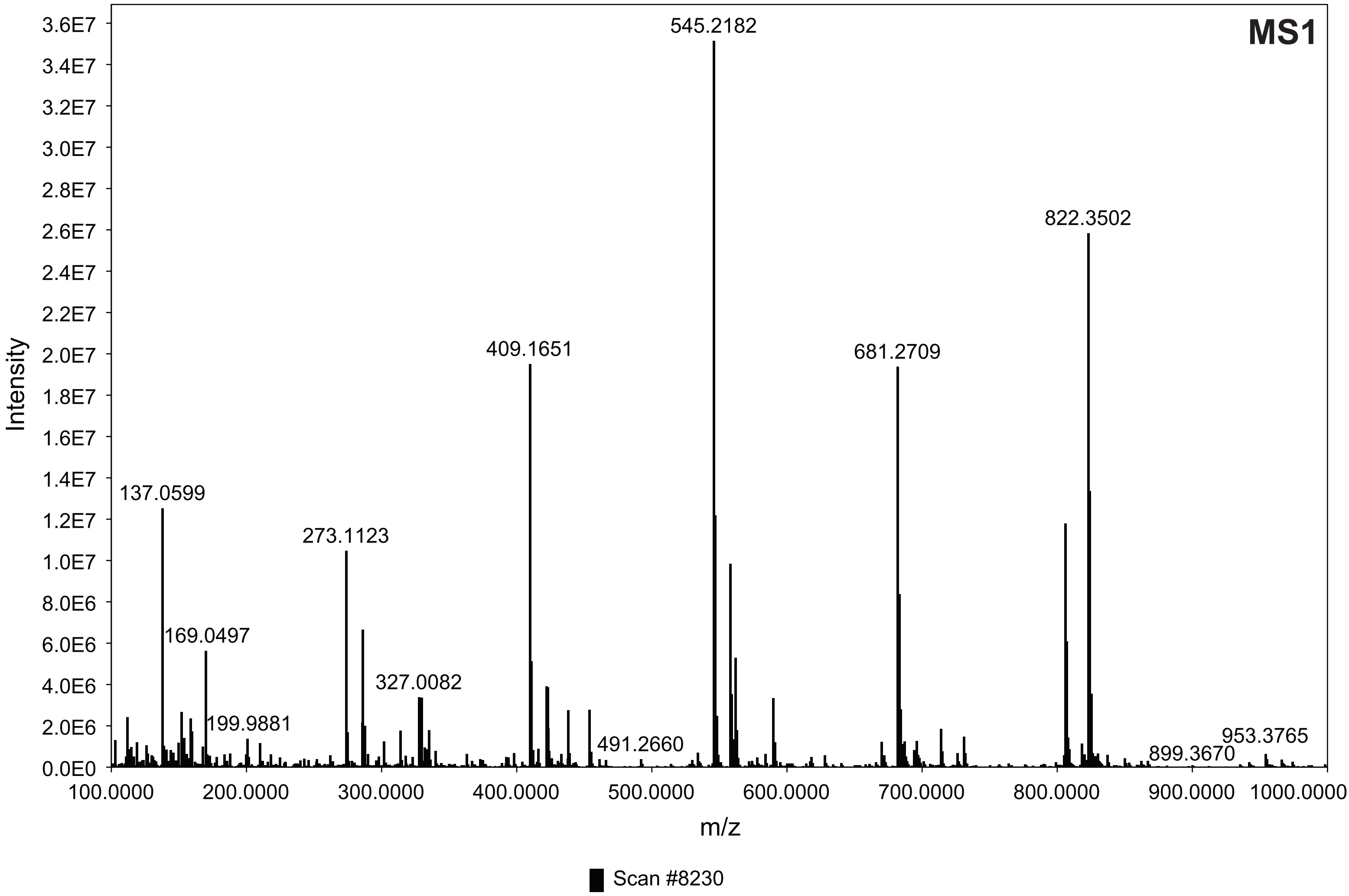


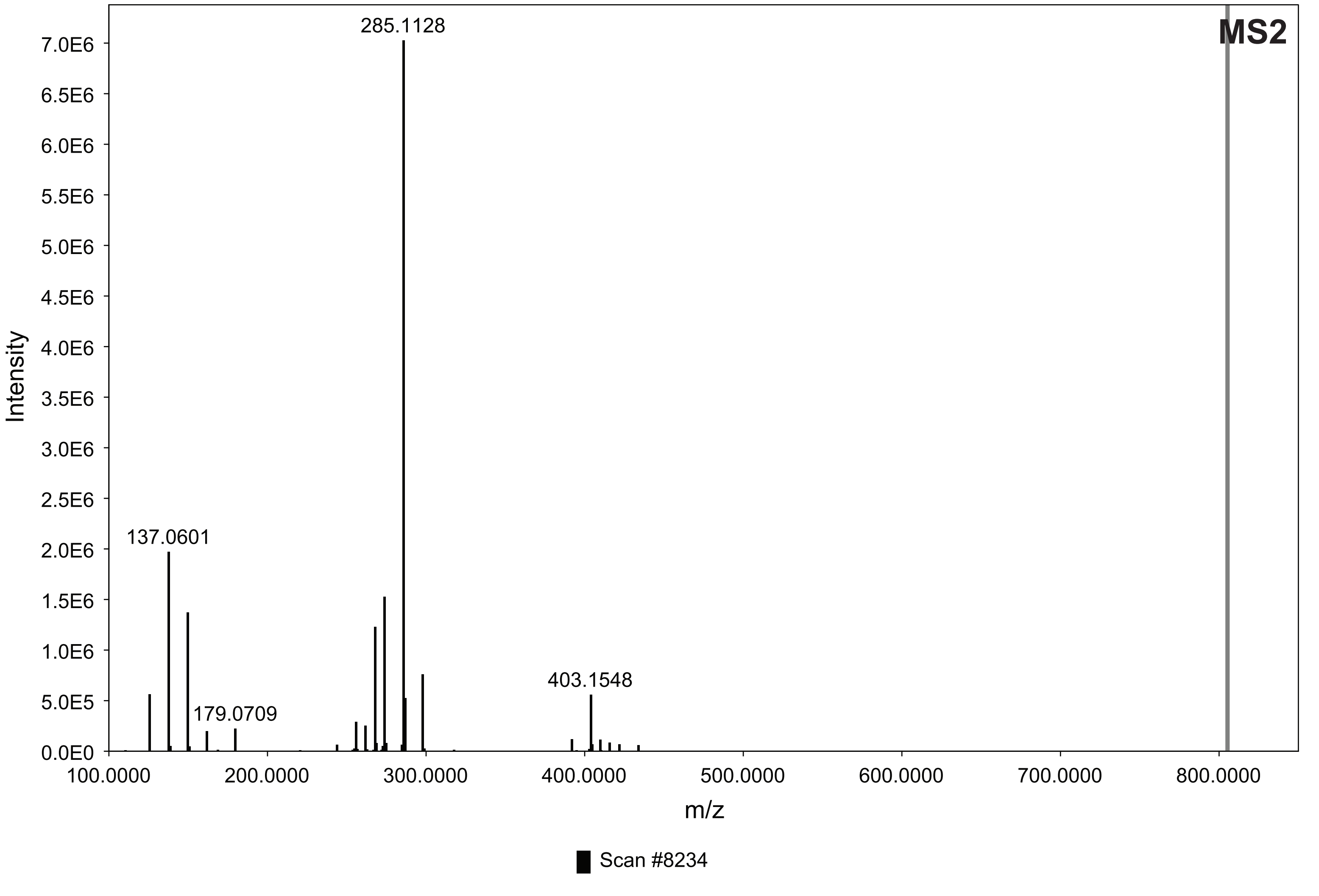

**12. LC-MS m/z 847 $[M-H]^-$; HRMS m/z 849.3132 $[M+H]^+$**

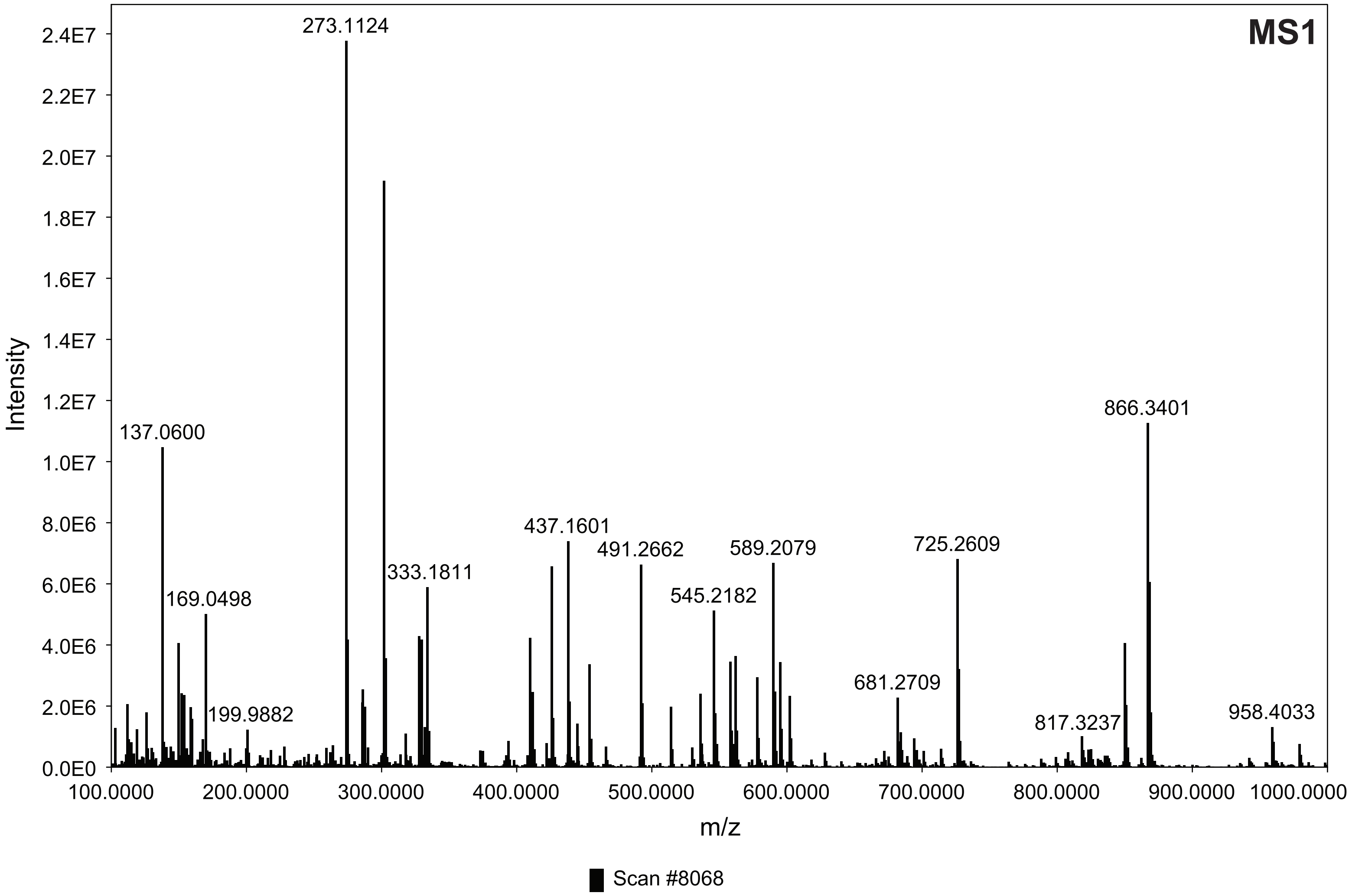


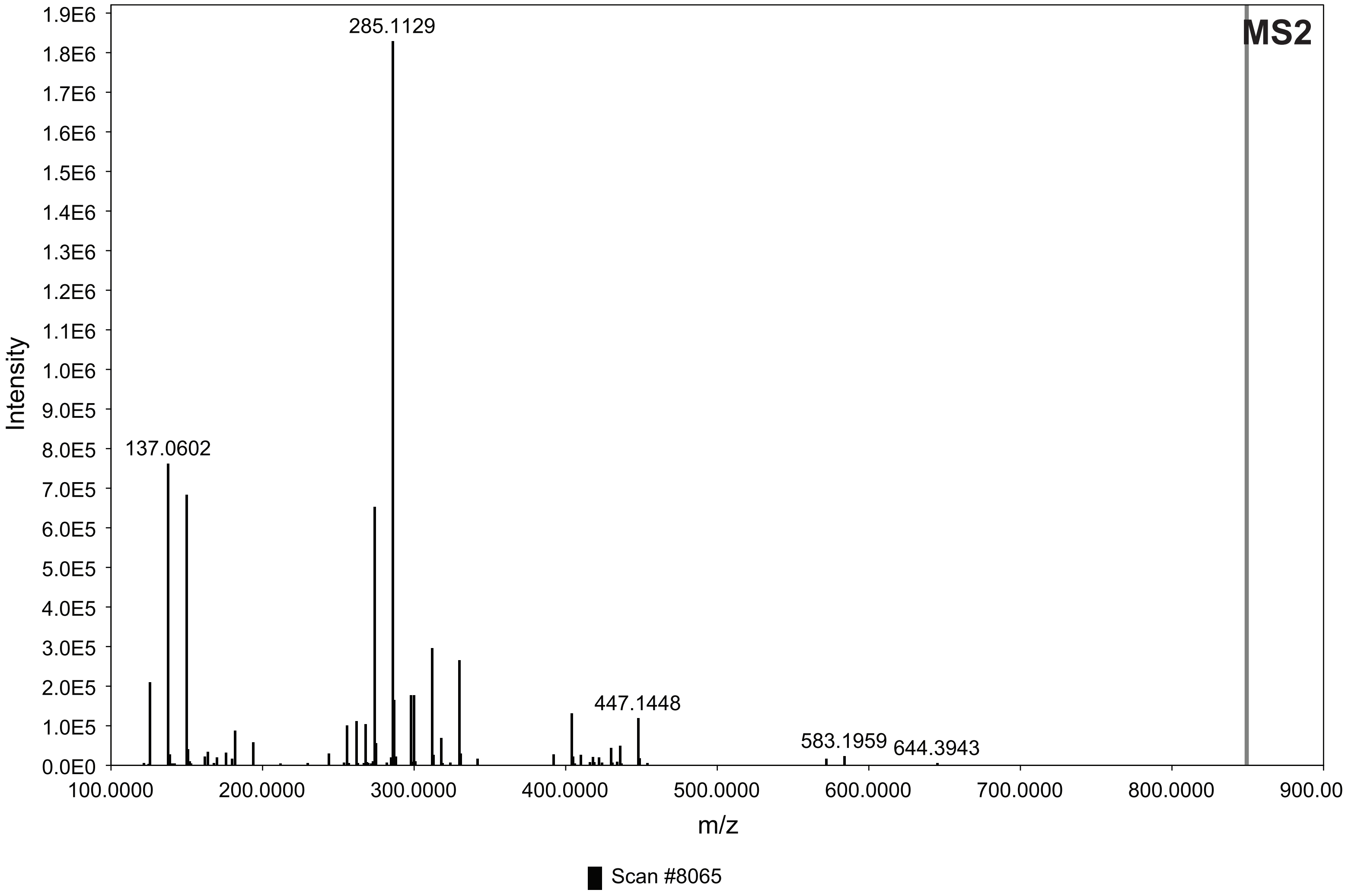

### 13. LC-MS m/z 847 [M-H]⁻; HRMS m/z 849.3132 [M+H]⁺

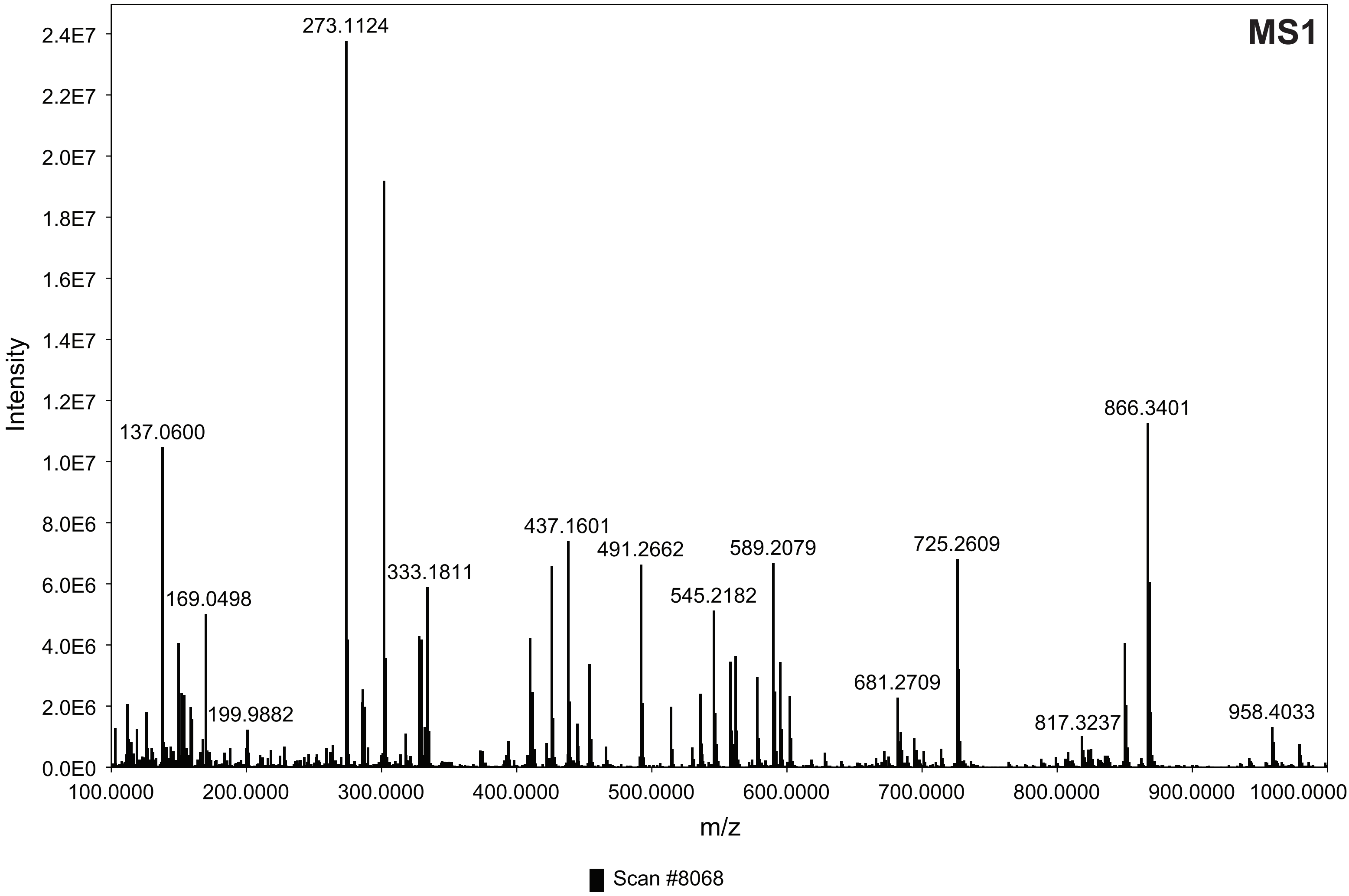


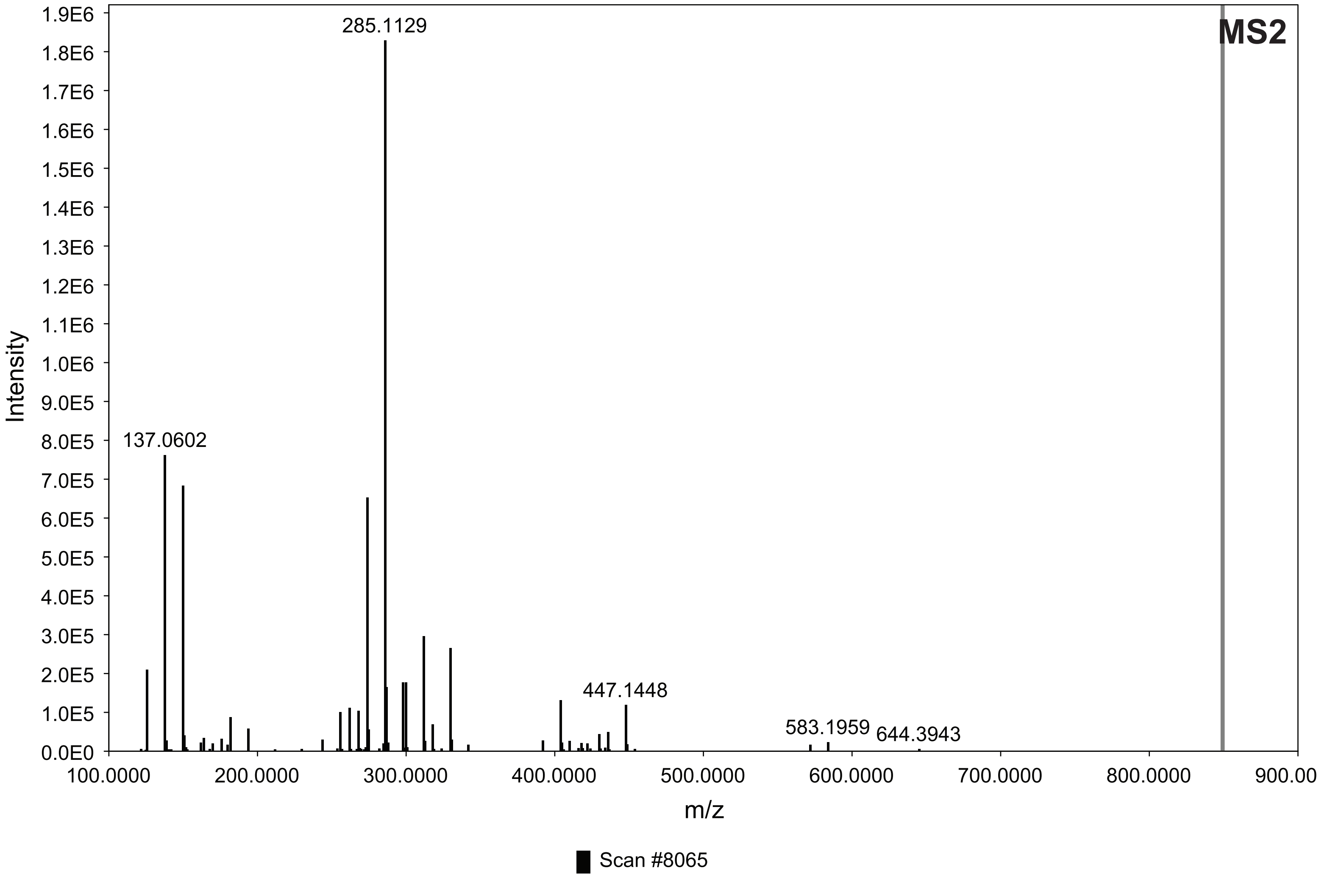

**14. LC-MS m/z 939 [M-H]$^-$; HRMS m/z 941.3759 [M+H]$^+$**

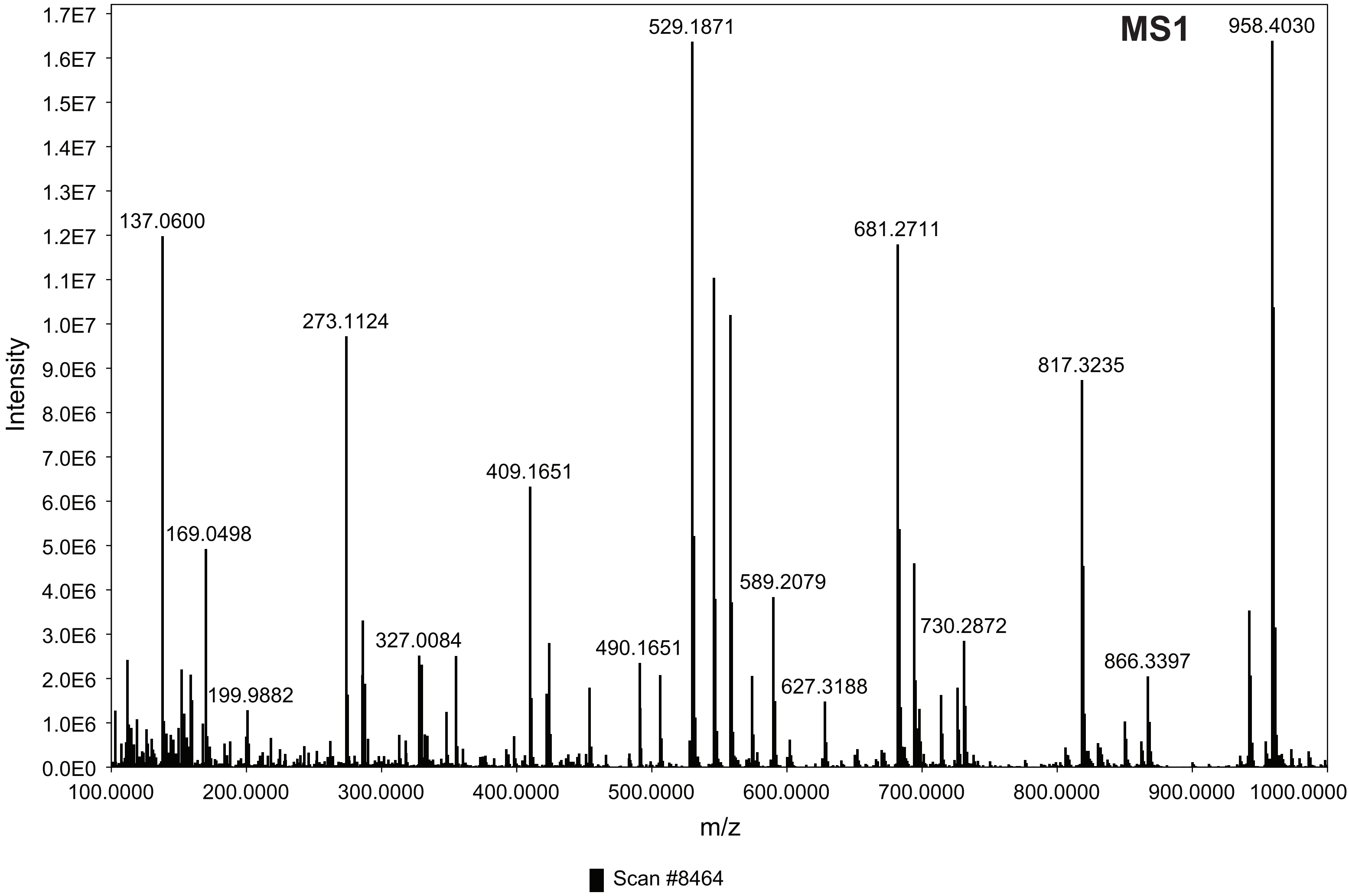


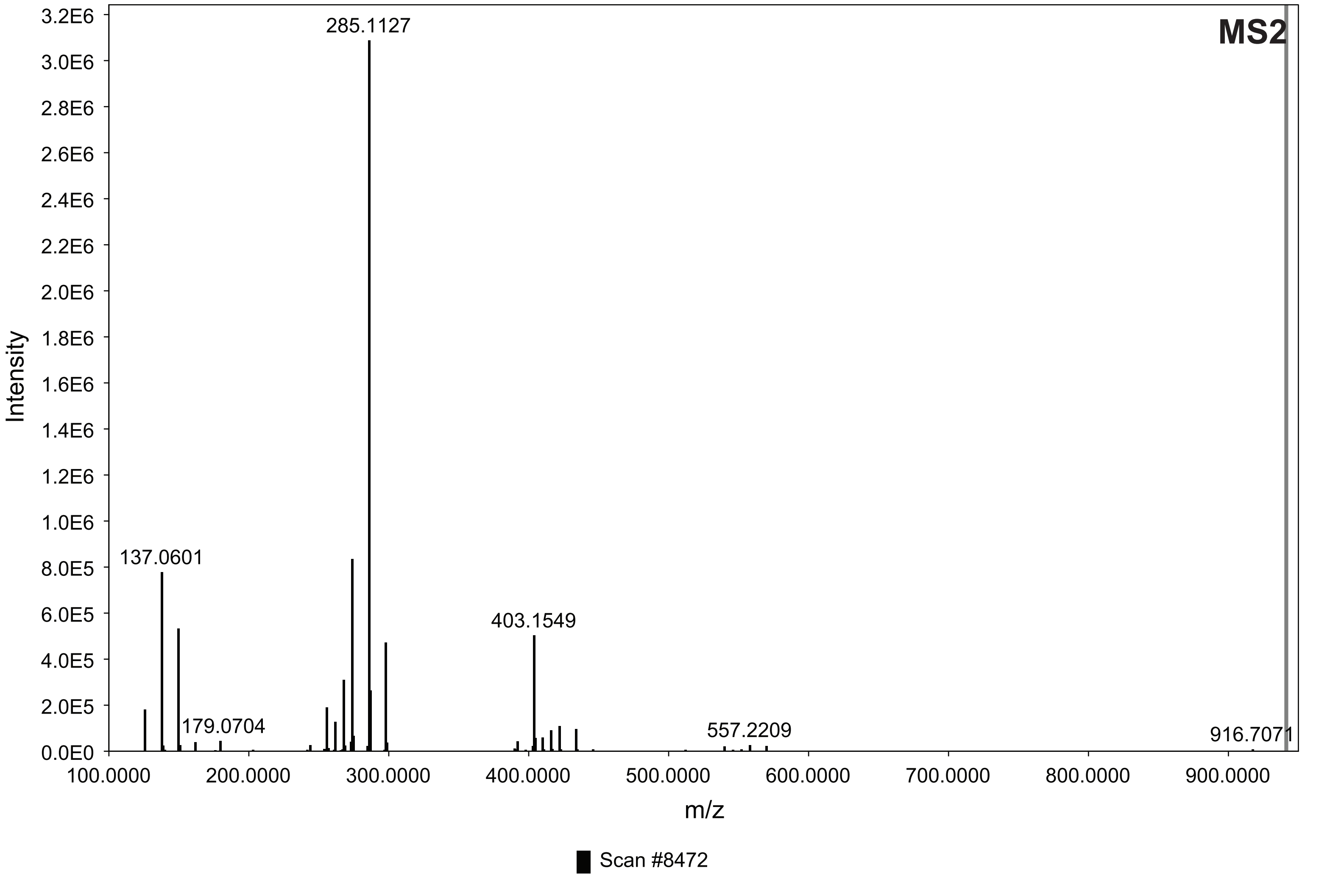

**15. LC-MS m/z 983 [M-H]⁻; HRMS m/z 985.3662 [M+H]⁺**

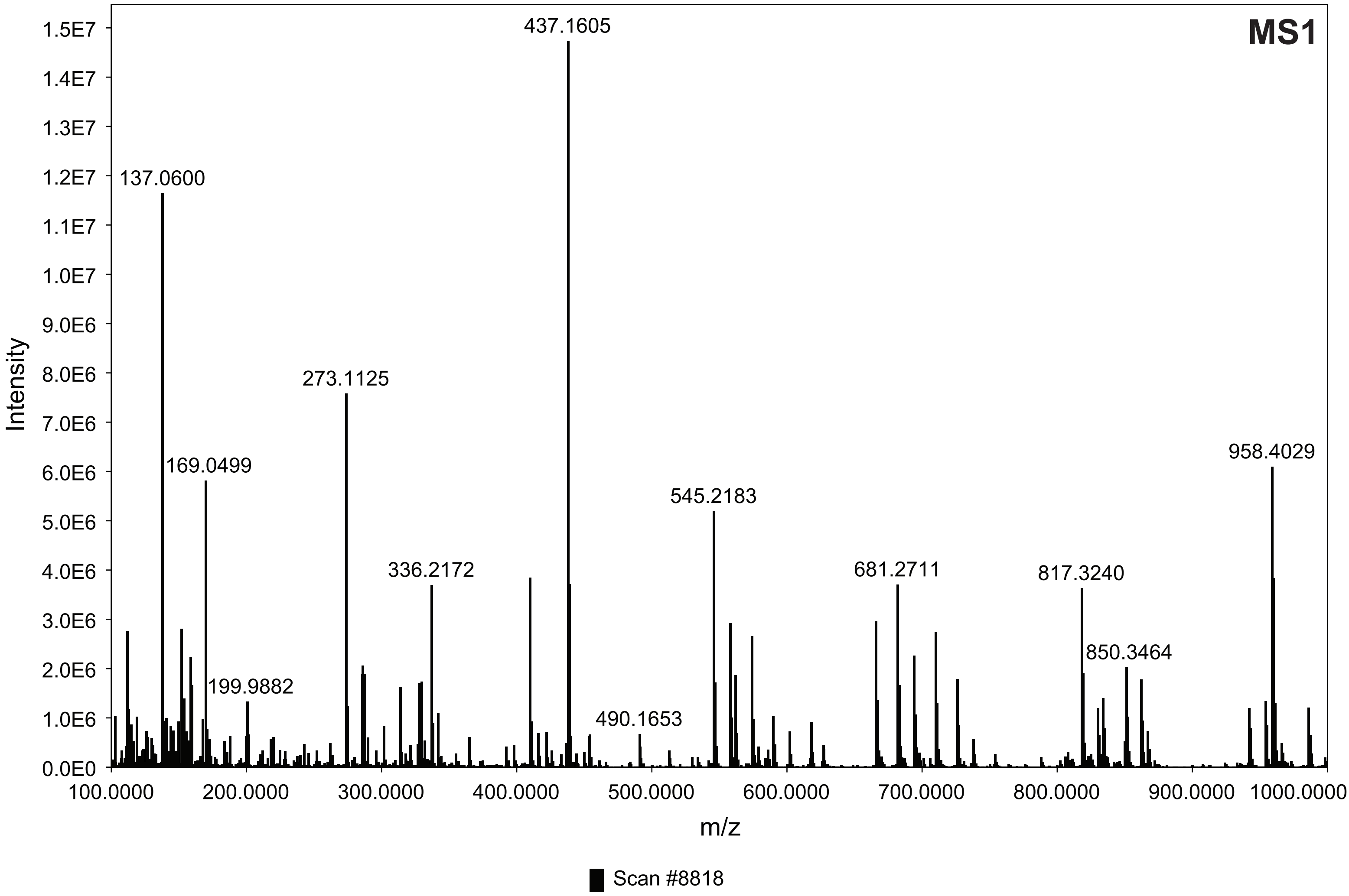


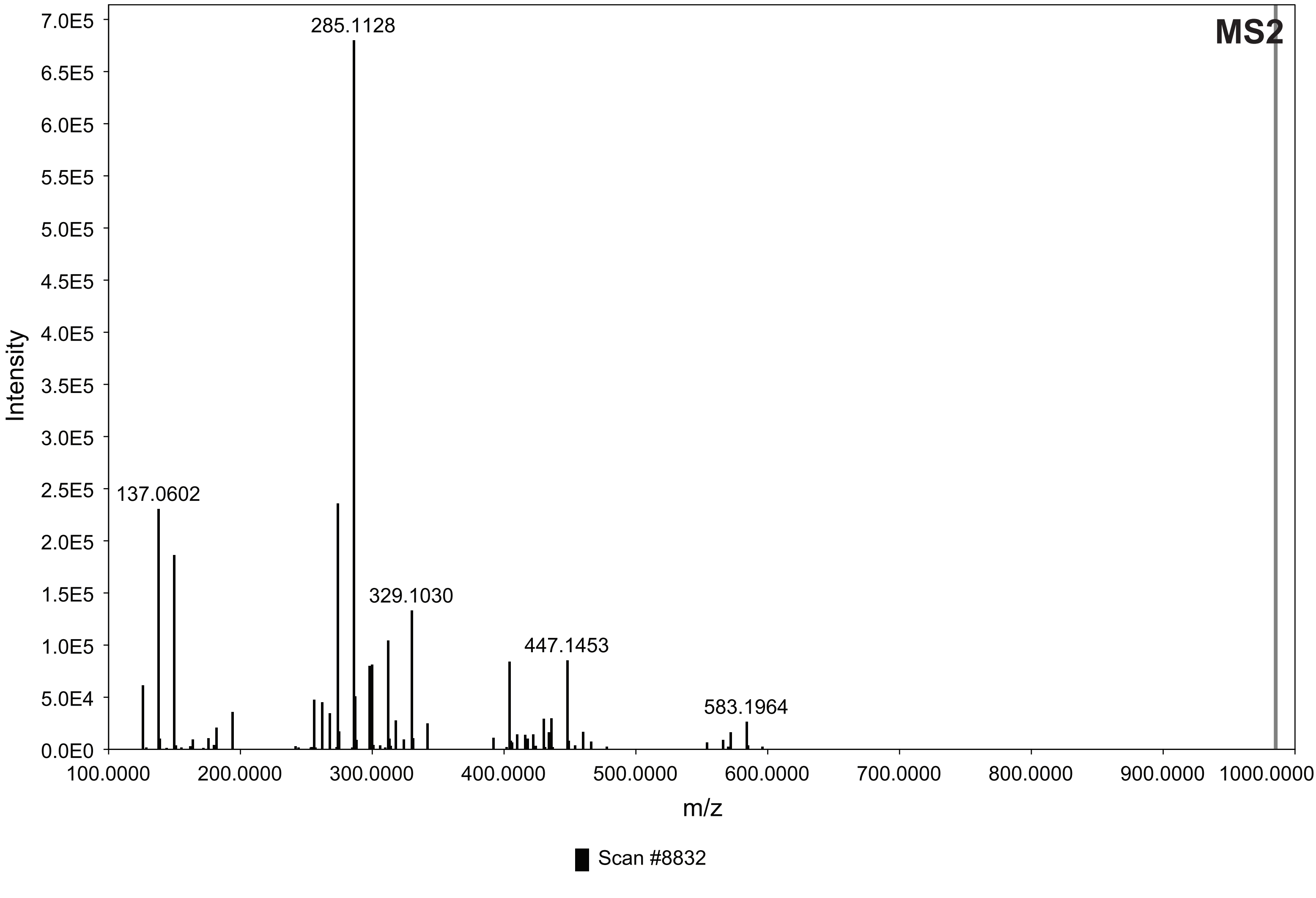
